# Identifying the core genome of the nucleus-forming bacteriophage family and characterization of *Erwinia* phage RAY

**DOI:** 10.1101/2023.02.24.529968

**Authors:** Amy Prichard, Jina Lee, Thomas G. Laughlin, Amber Lee, Kyle P. Thomas, Annika Sy, Tara Spencer, Aileen Asavavimol, Allison Cafferata, Mia Cameron, Nicholas Chiu, Demyan Davydov, Isha Desai, Gabriel Diaz, Melissa Guereca, Kiley Hearst, Leyi Huang, Emily Jacobs, Annika Johnson, Samuel Kahn, Ryan Koch, Adamari Martinez, Meliné Norquist, Tyler Pau, Gino Prasad, Katrina Saam, Milan Sandhu, Angel Jose Sarabia, Siena Schumaker, Aaron Sonin, Ariya Uyeno, Alison Zhao, Kevin Corbett, Kit Pogliano, Justin Meyer, Julianne H. Grose, Elizabeth Villa, Rachel Dutton, Joe Pogliano

**Affiliations:** School of Biological Sciences, University of California San Diego, La Jolla, CA, USA; Department of Cellular and Molecular Medicine, University of California San Diego, La Jolla, CA, USA; Department of Microbiology and Molecular Biology, Brigham Young University, Provo, UT, USA; Howard Hughes Medical Institute, University of California San Diego, La Jolla, CA, USA

## Abstract

We recently discovered that some bacteriophages establish a nucleus-like replication compartment (phage nucleus), but the core genes that define nucleus-based phage replication and their phylogenetic distribution were unknown. By studying phages that encode the major phage nucleus protein chimallin, including previously sequenced yet uncharacterized phages, we discovered that chimallin-encoding phages share a set of 72 highly conserved genes encoded within seven distinct gene blocks. Of these, 21 core genes are unique to this group, and all but one of these unique genes encode proteins of unknown function. We propose that phages with this core genome comprise a novel viral family we term Chimalliviridae. Fluorescence microscopy and cryo-electron tomography studies of *Erwinia* phage vB_EamM_RAY confirm that many of the key steps of nucleus-based replication encoded in the core genome are conserved among diverse chimalliviruses, and reveal that non-core components can confer intriguing variations on this replication mechanism. For instance, unlike previously studied nucleus-forming phages, RAY doesn’t degrade the host genome, and its PhuZ homolog appears to form a five-stranded filament with a lumen. This work expands our understanding of phage nucleus and PhuZ spindle diversity and function, providing a roadmap for identifying key mechanisms underlying nucleus-based phage replication.

## INTRODUCTION

The ability to establish and maintain subcellular organization is fundamental to cellular function. Even many viruses remodel their host cells, setting up their own complex compartments to suit their unique needs for viral replication (Kieser et al., 2020; Knipe et al., 2022; Labarde et al., 2021; Paszkowski et al., 2016; Tomer et al.,2019; Trinh et al., 2020). We recently discovered that some bacteriophages replicate by creating a nucleus-like proteinaceous structure (the phage nucleus) that compartmentalizes the bacterial host cell during phage infection much in the same way a membranous nucleus compartmentalizes a eukaryotic cell (Birkholz et al., 2022; Chaikeeratisak, Nguyen, Egan, et al., 2017; Chaikeeratisak, Nguyen, Khanna, et al., 2017; Knipe et al., 2022; Laughlin et al., 2022). Although the phage nucleus is structurally different from the eukaryotic nucleus, it performs many similar functions. The phage nucleus, which is made of a protein called chimallin (ChmA), separates transcription from translation, exports mRNA, selectively imports proteins, and shields the phage DNA from cytoplasmic nucleases (Birkholz et al., 2022; Chaikeeratisak, Nguyen, Egan,et al., 2017; Chaikeeratisak, Nguyen, Khanna, et al.,2017; Laughlin et al., 2022; Malone et al., 2020;Mendoza et al., 2020; Nguyen et al., 2021).

Nucleus-forming phages belong to a larger group of phages that encode rifampicin-resistant multi-subunit RNA polymerases (msRNAP) (Ceyssens et al., 2014; Krylov et al., 2021; Matsui et al., 2017; Orekhova et al., 2019; Sokolova et al.,2020; Thomas et al., 2008, 2012; Yakunina et al.,2015). Nucleus forming phages, typified by *Pseudomonas aeruginosa* phage ΦKZ, encode two msRNAP composed of 4-5 subunits. One of these msRNAPs is the virion RNAP (vRNAP), which is packaged within the capsid and is likely injected into the cell along with the DNA upon infection (Ceyssens et al., 2014; Thomas et al., 2008, 2012). The non-virion RNAP (nvRNAP) is expressed by the vRNAP during infection (Ceyssens et al., 2014;Orekhova et al., 2019; Yakunina et al., 2015). While all nucleus-forming phages studied to date encode these unique msRNAPs, this feature is also observed in several jumbo phages that do not form phage nuclei (Sokolova et al., 2020).

Another feature shared by currently characterized nucleus-forming phages is a phage-encoded tubulin homolog called PhuZ. Members of the PhuZ family studied thus far share a common filament structure and assembly mechanism, producing a dynamic three-stranded filament (Aylett et al., 2013; Zehr et al., 2014). In *Pseudomonas* phages 201ϕ2-1, ΦKZ, and ΦPA3, the phage nucleus is centered at the host midcell and rotated by a bipolar PhuZ spindle (Chaikeeratisak et al., 2019; Chaikeeratisak,Nguyen, Egan, et al., 2017; Chaikeeratisak, Nguyen,Khanna, et al., 2017; Erb et al., 2014; Kraemer et al., 2012), and in *Escherichia coli* phage Goslar, the PhuZ filaments form a vortex that rotates the nucleus without positioning it at midcell (Birkholz et al., 2022). Capsids assemble on the membrane and traffic along PhuZ filaments to reach the nucleus in the *Pseudomonas* phages (Chaikeeratisak et al., 2019; Chaikeeratisak,Nguyen, Egan, et al., 2017; Chaikeeratisak, Nguyen, Khanna, et al., 2017). Capsids then dock to initiate DNA packaging at the surface of the phage nucleus (Birkholz et al., 2022; Chaikeeratisak et al., 2019; Chaikeeratisak, Nguyen, Egan, et al., 2017; Chaikeeratisak, Nguyen, Khanna, et al., 2017). Filled capsids assemble with tails, forming cytoplasmic bouquet structures at rates varying from phage to phage prior to cell lysis (Birkholz et al., 2022;Chaikeeratisak et al., 2022).

These recent discoveries prompt intriguing questions: How widespread is nucleus formation among phages that infect different hosts? Do all nucleus-forming phages share a common set of core genes? If so, which viral genes are part of the core genome versus the accessory genome? To answer these questions, we began by studying *Erwinia* phage vB_EamM_RAY (hereafter RAY), which encodes a chimallin homolog. RAY infects *Erwinia amylovora,* an important agricultural pathogen and the causative agent of fire blight (Sharma et al., 2019; Vanneste, 2000). Many chimallin-encoding *Erwinia* phages are distantly related to each other and to known nucleus-forming phages, making *Erwinia amylovora* an enticing host for studying possible nucleus formation in diverse phages (Arens et al., 2018;Esplin et al., 2017; Sharma et al., 2019).

Core genomes of phage and bacteria define their conserved components, while accessory genomes provide genes that are specific to the individual species, strains, or variants (Cazares et al., 2014; Comeau et al., 2007; Konstantinidis & Tiedje, 2005; Mathee et al., 2008; Park et al., 2019). While genes required for making viral particles such as virion structural components, polymerases, and chimallin can be identified bioinformatically by comparison to sequence and structural homologs, the genes that comprise the nucleus-forming phage core genome have not been previously identified. Studying the core genome will allow us to identify and understand the genes required for the nucleus-based replication mechanism, including nuclear shell assembly, PhuZ spindle formation, mRNA export, selective protein import, capsid docking, and bouquet formation. While previous work identified some core genome components common of chimallin-encoding *Pseudomonas* phages 201ϕ2-1, ΦKZ, ΦPA3, EL, and OBP, and *Vibrio* phage JM-2012, this work predates the discovery of the phage nucleus replication mechanism, and only 6 phage genomes were included at the time (Jang et al., 2013). Additionally, core genomes of large groups of jumbo phages have been analyzed, but did not focus on viral replication mechanisms to identify the core genome of phage that replicate by forming a nucleus (Iyer et al., 2021).

Here we investigated the conservation of nucleus-based phage replication by focusing on *Erwinia* phage RAY and comparing it to previously studied nucleus-forming phages. We analyzed its phylogenetic relationships to other phages, identified the core and accessory genes, and used fluorescence microscopy and bioinformatics to characterize the putative functions of fifteen core and nine accessory RAY proteins. We show that RAY is a nucleus-forming phage by fluorescence microscopy and cryo-electron tomography and describe how its replication mechanism varies compared to previously characterized nucleus-forming phages.

## RESULTS

### Defining the Core Genome of Chimallin-Encoding Phages

To identify the key, conserved genes required to replicate via the phage nucleus pathway, we first identified all phages in the NCBI database that encode a homolog of the major nuclear shell protein chimallin (ChmA). We found 66 unique phages encoding chimallin homologs and made whole genome trees to compare them (Fig. 1A,B), showing that all 66 phages form a monophyletic group (Fig. 1A) when compared with phages that encode msRNAPs and other related phages (see methods). Members of this clade infect a wide range of Gram-negative bacteria and one Gram-positive *Bacillus* species, and they vary greatly in genome size from 167kb to 322kb. This suggests that the chimallin-encoding group of phages arose only once from a common ancestor, has a wide host range, and is not restricted only to “jumbo” phages larger than 200kb. We created phylogenetic trees based on six conserved proteins. The whole genome tree (Fig. 1B) is generally congruent with all of the protein-based trees, including chimallin (Fig. 1C), major capsid protein (Fig. 1D), terminase large subunit (Fig. S1A), DNA polymerase (Fig. S1B), replicative helicase (Fig. S1C), and an RNA polymerase subunit (Fig. S1D). Each of these trees contained 16 groups of related phage species composed of the same individuals (color coded in Fig. 1, Fig. S1, Fig. S2). The phylogeny of the groups closely matched the whole genome tree (Fig. S2). The similarity across different phylogenetic trees suggests that these proteins have been co-evolving with little evidence of horizontal gene transfer between divergent phages for these proteins. This supports previous findings that the phage nucleus may help to limit recombination between nucleus-forming phages (Chaikeeratisak et al., 2021). We then determined the set of core genes that are conserved among chimallin-encoding phages and whether this core genome is conserved in other msRNAP-encoding phages.

**Figure 1.**
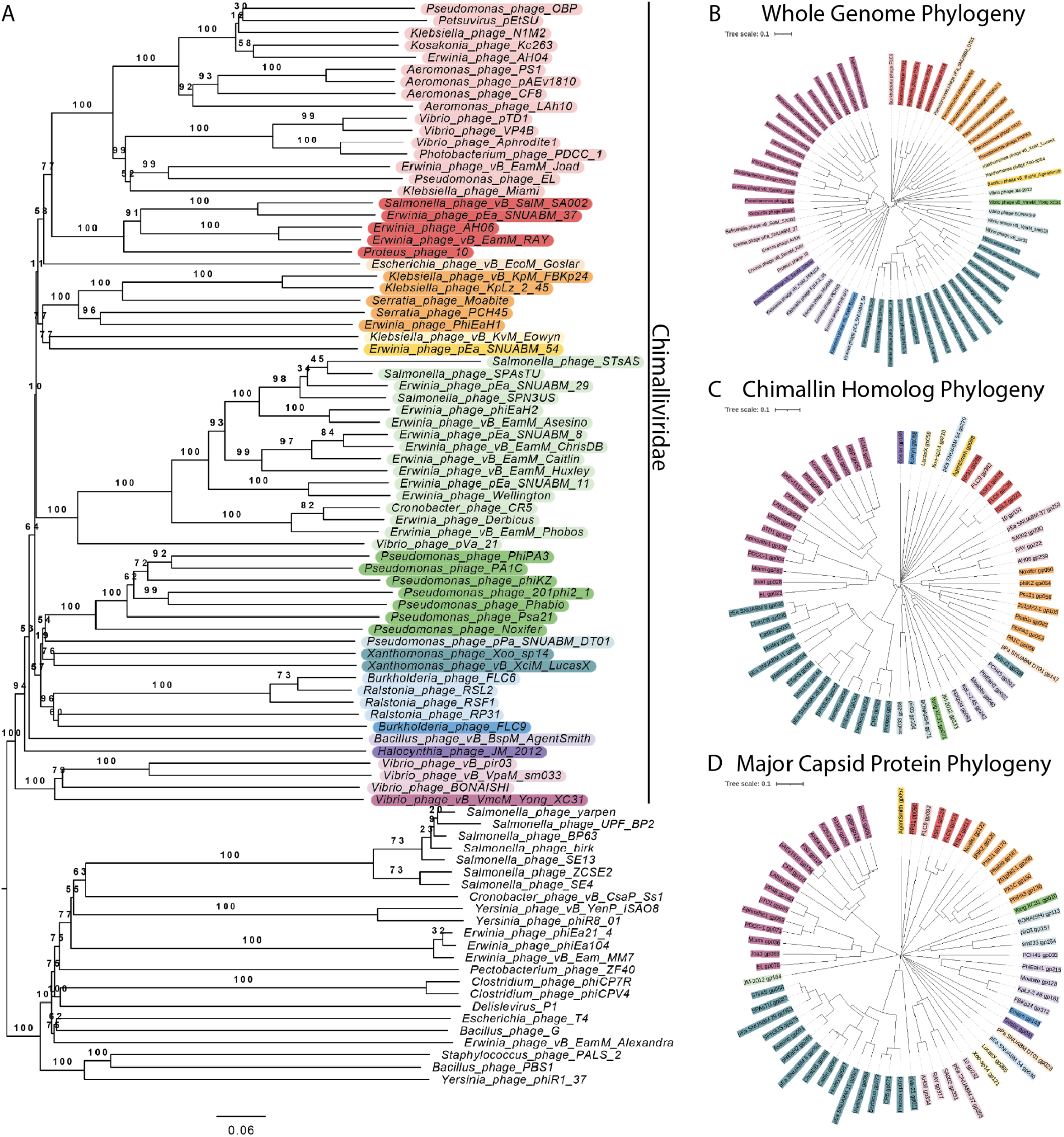
Phylogenetic comparison of Chimalliviridae. (A) Phylogenetic tree of the Chimalliviridae and potentially related phages. The Chimalliviridae appear to form one clade, separate from all potentially related groups included (msRNAP-encoding phages, ViPTree-predicted relatives, and other phages with large genomes), which has been color coded in rainbow colors by predicted genus to improve readability. (B) Phylogenetic tree based on whole genome comparison of 66 chimallin-encoding phages (Table 1). (C) Phylogenetic tree based on the protein sequences of the chimallin homologs from the 66 phages used in our analysis. (D) Phylogenetic tree based on the major capsid proteins from the 66 phages used in our analysis. Trees were colored in iTOL (Letunic & Bork, 2021) by predicted genus (Table 2) where predicted genus 28 is light red, 18 is red, 24 is light orange, 27 is orange, 3 is light yellow, 9 is yellow, 25 is light green, 20 is green, 6 is light teal, 8 is teal, 17 is light blue, 2 is blue, 7 is light purple, 13 is purple, 29 is light pink, and 4 is pink. Some of these predicted genera are broken up as currently classified by the ICTV, but for maximum readability of the colors, only the genera predicted by VICTOR are color-coded.

We classified a gene as core if it is present in more than 90% of the chimallin-encoding phages to accommodate potential sequencing errors and variability due to sampling (van Tonder et al., 2014) and found 72 highly conserved genes (Fig. 2A-C) (see methods for details). The 72 core genes included the major capsid protein, terminase large subunit, and msRNAP subunits, but surprisingly, the majority of the highly conserved core genes (53, 73.6%) had no predicted function. A PhuZ homolog was present in only 66.2% of chimallin-encoding phages, which is notable given its well-characterized role in nucleus-based phage replication yet consistent with PhuZ function being dispensable for replication (Birkholz et al., 2022;Chaikeeratisak et al., 2019, 2021; Guan et al., 2022;Kraemer et al., 2012). The core genes identified here likely contain many of the key proteins required for nucleus-forming phages to replicate.

**Figure 2.**
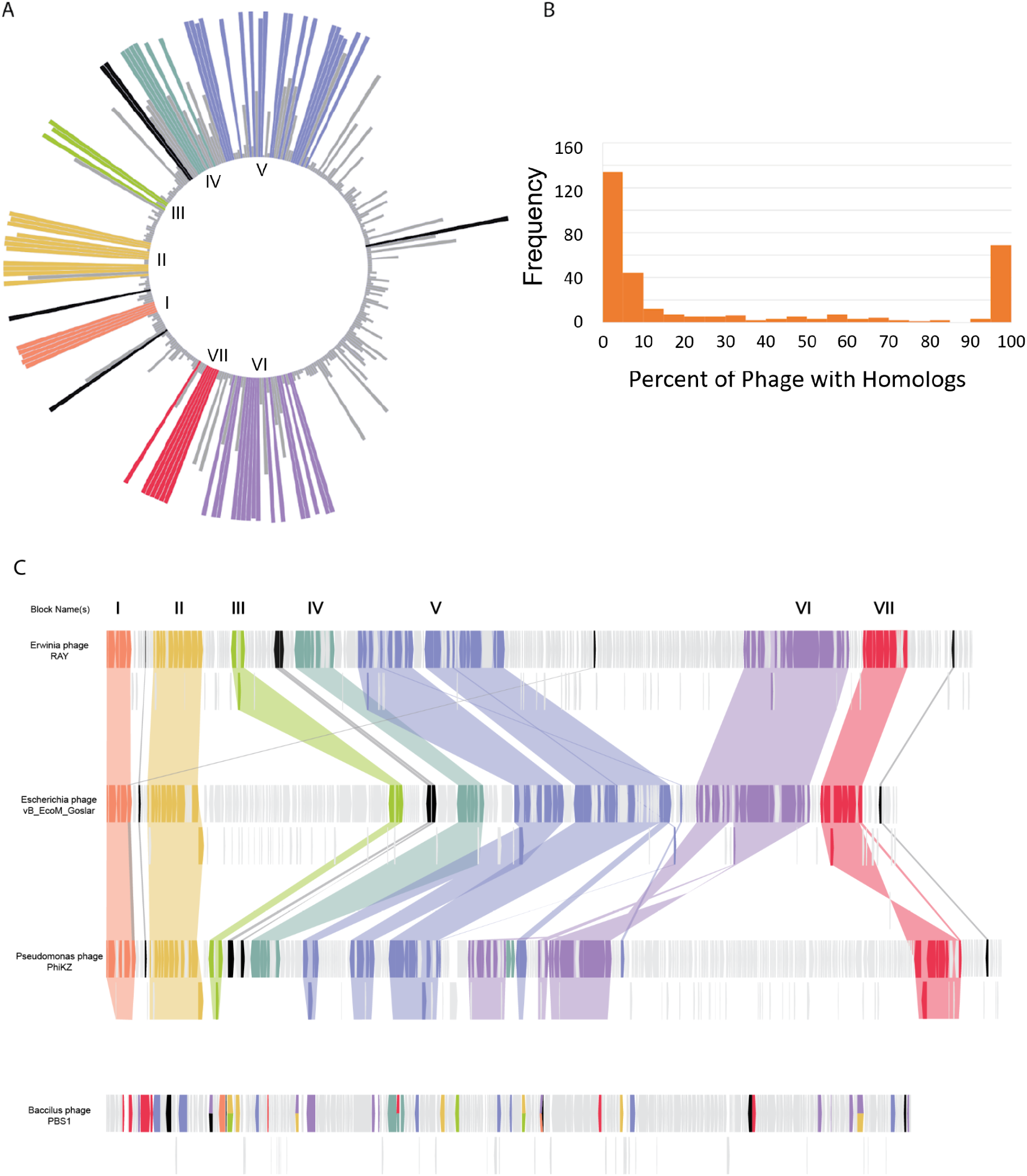
Core genome determination. (A) A circular bar plot of RAY’s genome with heights of bars corresponding to the percentage of phage that contain genes homologous to each RAY gene. Certain regions are more variable (right of circle), while other regions are highly conserved. The highly conserved regions with more than 3 genes are annotated in colors as different blocks. Core genes that are not part of blocks are colored black. (B) Histogram showing the frequency of conservation of each RAY protein among chimallin-encoding phages. The left peak indicates RAY-specific proteins (conserved in less than 10% of Chimallin-encoding phages) and the right peak indicates core proteins (conserved in 90% or more Chimallin-encoding phages). (C) A genome map of chimallin-encoding *Erwinia* phage RAY, *Escherichia* phage Goslar, *Pseudomonas* phage ΦKZ, and *Bacillus* phage PBS1. PBS1 was included for comparison since it encodes the unique msRNAP but does not encode chimallin. The PSI-BLAST homologs for each RAY protein are colored accordingly in the three other phages. While in chimallin-encoding phage the genes are conserved in seven blocks (colored as in A), in PBS1 some of the core genes are present but are dispersed across its genome rather than being encoded in blocks. Goslar gp188 and ΦKZ gp055 are homologous to each other and to two RAY genes, gp223 (orange) and gp070 (black). The full list of genes and blocks can be found in Table 3.

The core genes occur in seven distinct blocks (of three or more genes) whose general order is conserved across *Pseudomonas aeruginosa* phage ΦKZ, *Escherichia coli* phage vB_EcoM_Goslar (Goslar), and *Erwinia amylovora* phage RAY (Figure 2C). These three phages were chosen for our analysis because they represent a diverse group of chimallin-encoding phages and are well-studied (ΦKZ), infect highly tractable hosts (Goslar), or formed the basis of our bioinformatic studies (RAY). The core genome blocks are often rich with certain types of genes, such as Block 7 which contains the terminase and a handful of virion structural genes, and Block 5 which contains nineteen genes, including seven structural genes, the major capsid protein, one helicase, and an RNA polymerase beta subunit (Table 3). Gene order within the blocks is also conserved across chimallin-encoding phages. Upon mapping the level of conservation across the RAY, ΦKZ, and Goslar genomes, we found that there is a conserved region that is dense with core genome blocks and a variable region with putative accessory genes specific to each phage (Fig. 2A,C). The majority of RAY’s 317 genes are either part of this core conserved genome (23%, 72 genes) or are accessory genes only detectable in 10% or fewer of the phages we used in our analysis (178 genes, 56%) (Fig. 2B). This is a similar organization to other previously reported phage core genomes (Cazares et al., 2014; Comeau et al., 2007).

We next identified which of the core genes are specific to chimallin-encoding phages and likely represent the key genes required to replicate via the phage nucleus pathway, versus core genes that are broadly conserved among phage in general. We found that 51 out of the 72 genes are also present in other phages lacking chimallin homologs (Table 1). These broadly conserved proteins include DNA polymerases, RNA polymerase subunits, the major capsid protein, and other virion structural proteins. The remaining 21 genes include chimallin and 20 hypothetical proteins with no predicted structure or function. This unique set of 21 genes likely encodes the proteins specifically required to form and maintain a phage nucleus. We propose that all phages encoding chimallin and the associated core genome belong to one viral family, Chimalliviridae, named after the shared protein that makes up the phage nucleus lattice.

### Phage Nucleus Formation by *Erwinia* Phage RAY

Since RAY is a member of the Chimalliviridae and shares the core genome, we predicted that it replicates by forming a phage nucleus. We tested this prediction by imaging infected *Erwinia* cells in the presence of DAPI to determine whether phage DNA accumulates in the center of the cell. We observed a bright zone of DAPI fluorescence that was often positioned at midcell, similar to *Pseudomonas* nucleus-forming phages (Fig. 3A). We created an N-terminal fusion of GFPmut1 to gp222 (RAY chimallin homolog, ChmA_RAY_) (Fig. S3) and examined its localization in the absence and presence of phage infection. Fluorescence from GFP-ChmA_RAY_ was uniformly distributed throughout uninfected cells but formed a ring in the center of the cell enclosing DNA during RAY infections (Fig. 3B). Zones of DAPI fluorescence consistent with host chromosomal DNA were also present outside of the nucleus in every RAY infection, unlike previously characterized nucleus-forming phages which degrade the host DNA (Birkholz et al., 2022; Chaikeeratisak, Nguyen,Egan, et al., 2017; Chaikeeratisak, Nguyen, Khanna,et al., 2017) but similar to *Serratia* phage PCH45 (Malone et al., 2020). To determine whether the extranuclear DAPI staining was due to host DNA, we tagged the *Erwinia amylovora* H-NS protein with GFP, which coats DNA, and used the H-NS-GFP fusion as a marker to visualize host DNA during phage infection, as H-NS is not imported into the phage nucleus. When we expressed H-NS-GFP in *Erwinia* cells and infected them with RAY, the H-NS-GFP fluorescence co-localized with the DAPI outside of the phage nucleus (Fig. 3C). In time lapse microscopy, we observed that the bacterial DNA was coated with H-NS before infection and was then pushed to the poles and compacted upon infection as the phage nucleus formed (Fig. 3D, Movie 1). These results suggest that RAY forms a nucleus-like structure without detectably degrading the host DNA.

**Figure 3.**
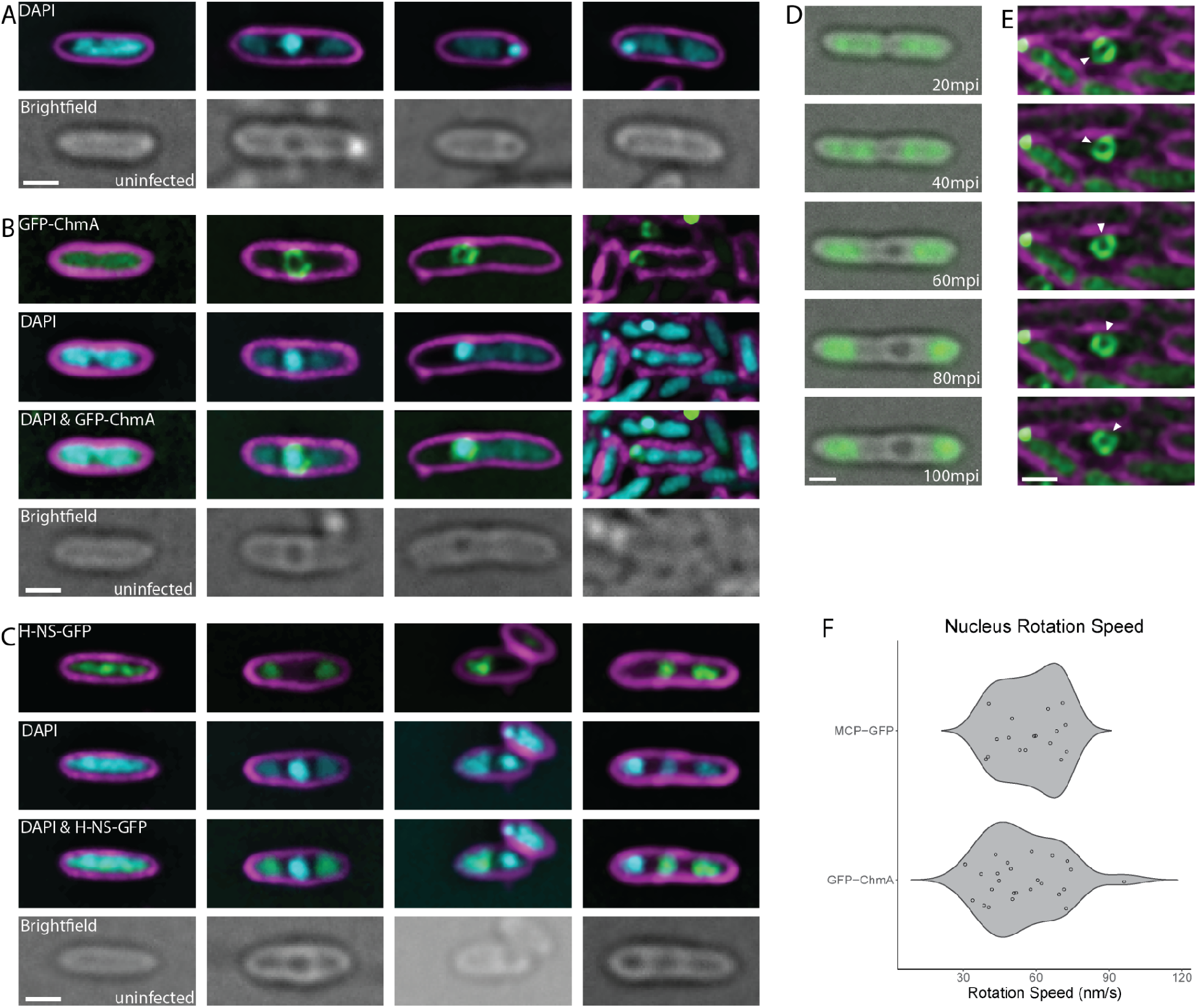
RAY phage nucleus formation. (A) DAPI localization showing the presence of a bright zone consistent with phage nucleus formation. Cells were imaged between 60-75 mpi (mid-infection) unless labeled as uninfected. (B) GFP-ChmA_RAY_ surrounds the bright DAPI zones, consistent with the morphology of a phage nucleus.Cells were imaged between 60-75 mpi (mid-infection) unless labeled as uninfected. (C) H-NS-GFP localizes to the DNA outside of the phage nucleus.Cells were imaged between 60-75 mpi (mid-infection) unless labeled as uninfected. (D) H-NS-GFP moves away from midcell during the course of infection (Movie 1). (E) The phage nucleus rotates during infection (Movie 2). Time lapse images were taken 4 seconds apart. The white arrow shows a dimmer segment of the nuclear shell that was tracked over the course of the rotation to determine the speed. (F) Violin plot showing the distribution of rotation speeds of individual measured RAY nuclei using either a chimallin (ChmA) (n=25, see dots) or major capsid protein (MCP) (n=17, see dots) GFP tag to track rotation. For all microscopy images (panels A,B,C,D,E), scale bar is 1 micron; magenta is membrane stain FM4-64, cyan is DNA stain DAPI, green is GFP, and grayscale is brightfield.

### Identification of RAY Nuclear and Cytoplasmic Proteins

To gain further insight into the subcellular organization of RAY’s lytic cycle, we used fluorescence microscopy to observe the localization of 24 RAY core and accessory proteins fused to GFP. To gain insights into both the conserved chimalliviridae replication mechanism as well as RAY-specific variations, we tagged fifteen proteins that were part of the Chimalliviridae core genome and nine that were accessory components. We were able to tentatively assign potential protein families to which most of these proteins belong based on *in silico* predictions; however, outside of the RNA polymerase subunits, few of these phage proteins have been studied in detail and their functions in the phage life cycle are unknown.

We identified ten proteins that co-localize with the phage DNA, likely inside the phage nucleus (Fig. 4A,B). Eight of these nucleus-associated RAY proteins (gp002, gp150, gp220, gp223, gp248, gp249, gp250, gp315) are part of the core genome, and two are not (gp116, gp153). Most of these proteins are likely involved in the core functions of DNA replication (gp220, gp315) (Fig. S4 & S5), transcription (gp002, gp248, gp223, gp249) (Fig. S6), or recombination (gp150, gp153) (Fig. S7 & S8) based on sequence and structural homology to previously characterized or annotated phage proteins. Two other proteins could be assigned to a protein family, but their functions are less clear (core gene gp250, SWI/SNF helicase family (Fig. S9) and accessory gene gp116, HslVU protease family (Fig. S10)).

**Figure 4.**
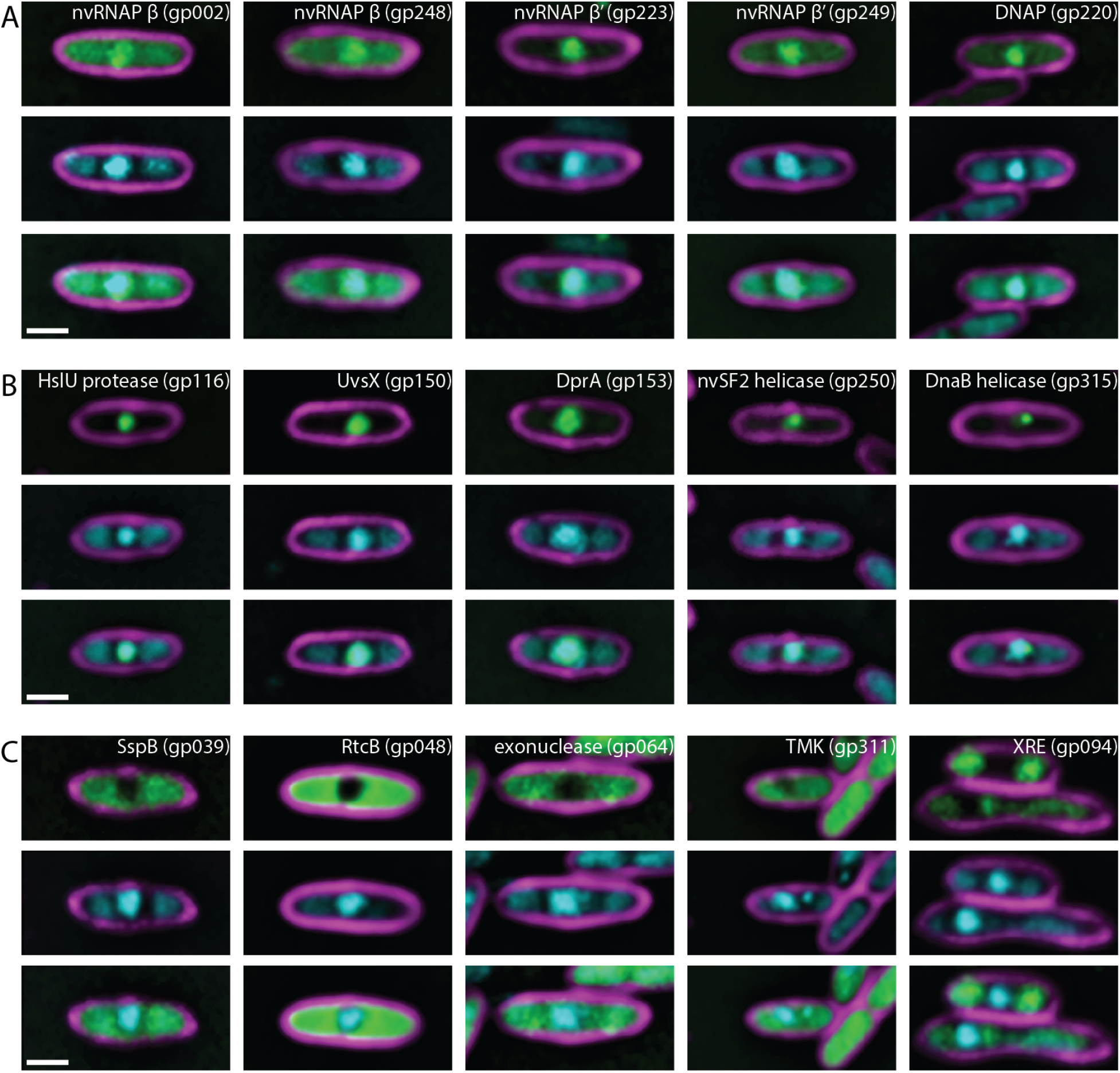
Localization of nuclear and cytoplasmic proteins encoded by RAY. (A) Putative RNA (nvRNAP **β** subunit 1 gp002, nvRNAP **β** subunit 2 gp248, nvRNAP **β’** subunit 1 gp223, and nvRNAP **β’** subunit 2 gp249) and DNA (DNA polymerase gp220) polymerases co-localize with phage DNA in the phage nucleus. (B) RAY gp116 is a homolog of HslU superfamily heat shock proteases, but other nuclear-localized phage proteins are homologs of known DNA-associated proteins such as proteins involved in recombination (UvsX gp150 and DprA gp153) and helicases (non-virion superfamily 2 helicase gp250 and DnaB-like replicative helicase gp315). (C) Cytoplasmic proteins are not predicted to be involved with DNA replication, recombination, or transcription. These include an SspB homolog gp039, RtcB homolog gp048, putative exonuclease gp064, thymidylate kinase gp311, and putative XRE superfamily transcriptional regulator gp094. For all images, scale bar is 1 micron; magenta is membrane stain FM4-64, cyan is DNA stain DAPI, and green is GFP.

We identified 4 proteins (gp039, gp048, gp064, and gp311) among our set of 24 that display diffuse localization in the cytoplasm outside of the phage nucleus and one protein (gp094) that localized on the host chromosome (Fig. 4C). Of these, only RAY gp311, a thymidylate kinase homolog (Fig. S11), is part of the core genome. Homologs of gp311 have also been shown to localize in the cytoplasm during replication of other phages (Birkholz et al., 2022; Chaikeeratisak, Nguyen, Khanna, et al., 2017), where they likely participate in nucleotide metabolism. The remaining four proteins included a putative SspB-like ClpXP adaptor protein (gp039, Fig. S12), a putative tRNA ligase (gp048, Fig. S13), a putative exonuclease (gp064, Fig. S14), and a putative XRE superfamily transcriptional regulator (gp094, Fig. S15) whose localization mimics that of bacterial H-NS by binding to host DNA (Fig. 3C, 4C). Taken together, these studies suggest that RAY forms a proteinaceous shell that encloses phage DNA and compartmentalizes proteins according to their functions.

### Virion-associated RAY Proteins

To follow the assembly process of RAY virions, we created GFP fusions to RAY’s tail sheath (gp179, Fig. S16) and major capsid (gp317, Fig. S17) proteins. Fluorescence microscopy showed that the major capsid protein localized around the nucleus (Fig. 5A), suggesting that capsids dock on the phage nucleus to initiate DNA packaging as occurs in ΦKZ-like phages and Goslar. The tail protein localizes in the cytoplasm, often forming foci near the phage nucleus (Fig. 5A); however, we did not observe bouquet-like clusters of virions. This suggests that capsids dock on the phage nucleus to be packaged with DNA, but after assembling virions, bouquet-like structures are not formed like those observed in other nucleus-forming phages (Birkholz et al., 2022; Chaikeeratisak et al., 2022).

**Figure 5.**
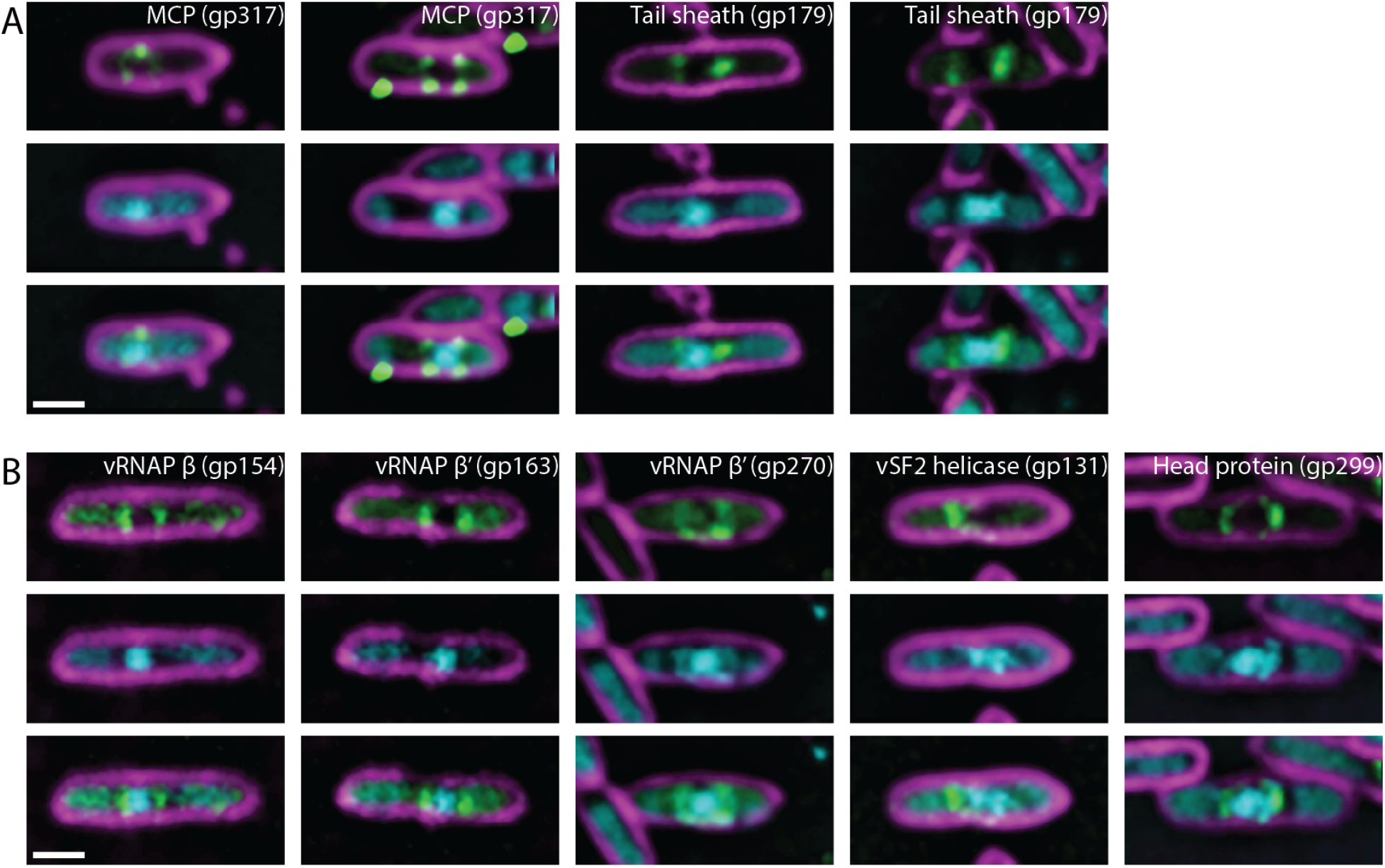
Localization of virion proteins encoded by RAY. (A) The major capsid protein gp317 and tail sheath gp179 are structural components of the RAY virion that localize near the periphery of the phage nucleus. Puncta can be seen near the membrane in the major capsid protein fusion, possibly because capsids are assembled here or alternatively as part of an aggregate formed due to overexpression of gp317-GFP. (B) Virion proteins (vRNAP **β** subunit 2 gp154, vRNAP **β**’ subunit 1 gp163, vRNAP **β**’ subunit 2 gp270, virion-associated superfamily 2 helicase gp131, and head protein of unknown function gp299) can be seen with similar localizations as the major capsid protein and sometimes (in the case of gp270 and gp299) have visible DAPI co-localized with them. For all images, scale bar is 1 micron; magenta is membrane stain FM4-64, cyan is DNA stain DAPI, and green is GFP.

We identified 5 proteins in our set of 24 that localized at the periphery of the phage nucleus during part of the lytic cycle, similar to the capsids (Fig. 5A,B). Three of these (gp154, gp163, and gp270, Fig. S18) are vRNAP subunits. The other two are a SWI/SNF family helicase (gp131, Fig. S19) and a putative head protein with unknown function (gp299, Fig. S20). Our localization of vRNAP and these other proteins during RAY infection is consistent with previous mass spectrometry results, which showed that RNA polymerase subunits are packaged within the viral capsid (Sharma et al., 2019; Thomas et al., 2012). These results also suggest that GFP fusions and fluorescence microscopy can be used as an additional approach to identify proteins that accumulate withincapsids.

### Phage Tubulin PhuZ

RAY encodes a PhuZ homolog (gp210, PhuZ_RAY_) (Fig. S21A) that contains conserved domains of the tubulin superfamily, including the tubulin T4 signature motif (Fig. 6A). In ΦKZ-like phages, PhuZ forms filaments that use dynamic instability and treadmilling to center and rotate the phage nucleus (Chaikeeratisak et al., 2019; Chaikeeratisak, Nguyen, Egan, et al., 2017; Chaikeeratisak, Nguyen, Khanna, et al., 2017; Erb et al., 2014; Kraemer et al., 2012), and in Goslar, PhuZ forms a vortex that rotates the phage nucleus without centering it (Birkholz et al., 2022). To determine whether PhuZ_RAY_ also forms filaments, we created a GFP-PhuZ_RAY_ fusion and assessed its ability to form polymers at different expression levels in both infected and uninfected cells. In uninfected cells, at low levels of arabinose (≤0.01%), GFP-PhuZ_RAY_ is uniformly distributed throughout the cell, but it spontaneously forms filaments at ≥0.05% arabinose (Fig. 6B,C). During infections, we studied GFP-PhuZ_RAY_ during infections under conditions where it did not spontaneously assemble filaments (0% arabinose) and found visible filaments in all infected cells (Fig. 6D,E), suggesting that the pre-expressed GFP-fusion in the host cell is able to co-assemble with phage-expressed native, untagged PhuZ.

**Figure 6.**
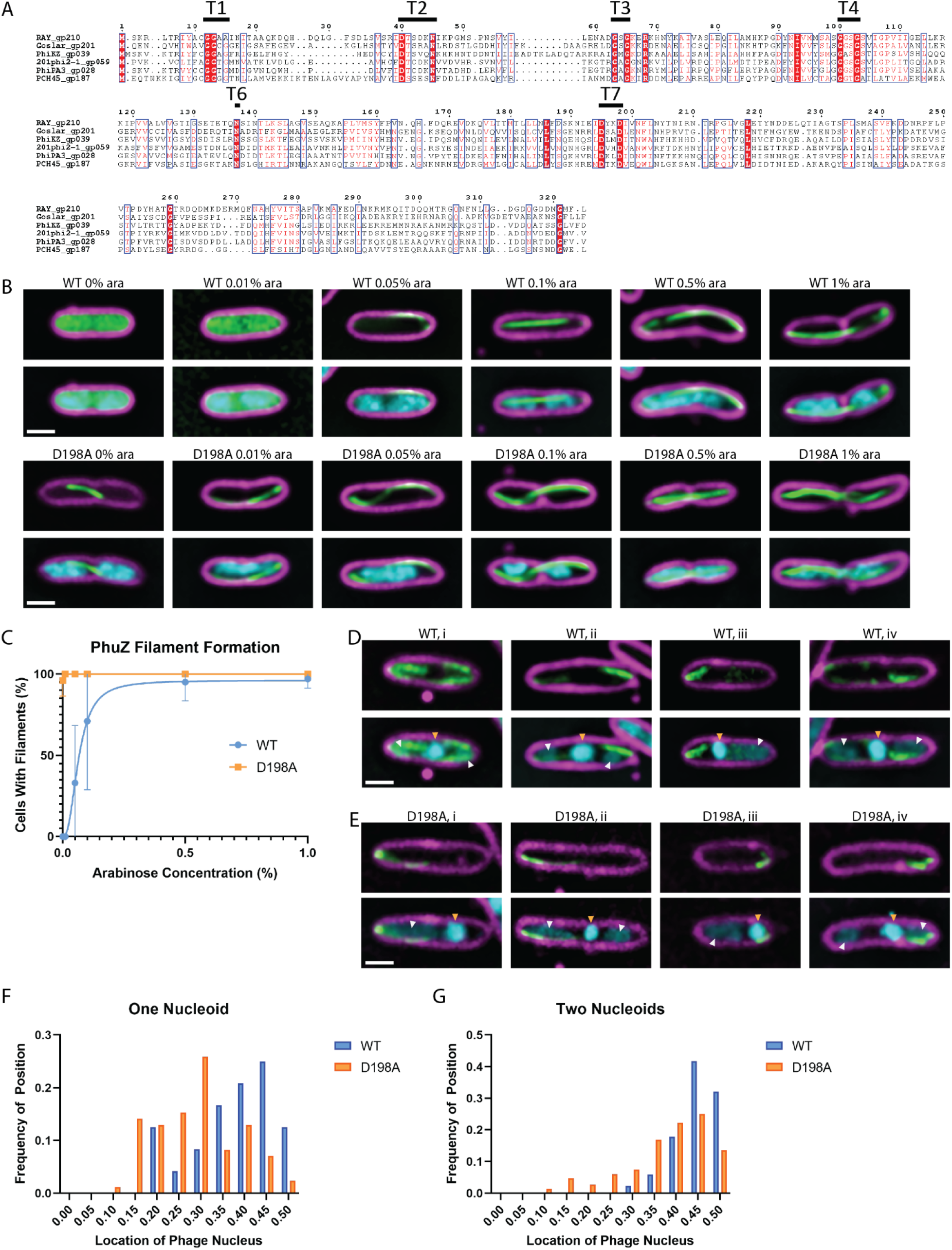
RAY PhuZ homolog. (A) PhuZ_RAY_ has conserved tubulin motifs, as marked in the sequence alignment. T5 is a structural motif with a poorly conserved sequence, so it is not labeled in this multiple sequence alignment. (B) Wild-type PhuZ_RAY_ does not polymerize spontaneously until it reaches a critical concentration, but the D198A mutant polymerizes at all levels of arabinose induction. (C) The average percentage of cells with PhuZ filaments is graphed by arabinose concentration. (D) Wild-type PhuZ filaments form with 0% arabinose during RAY infections. The nucleus is normally positioned near midcell (white arrows show bacterial DNA and gold arrows show phage nuclear DNA). (E) When the D198A mutant PhuZ is expressed, midcell localization of the phage nucleus during infection is not as common as with wild-type PhuZ (white arrows show bacterial DNA and gold arrows show phage nuclear DNA). (F) When only one bacterial nucleoid is present, the phage nucleus positioning histogram has a wide distribution for wild-type PhuZ (blue) and has no clear positioning bias for D198A mutant PhuZ (orange). (G) When two bacterial nucleoids are present, the wild-type PhuZ has a strong nucleus positioning bias toward midcell (blue), and the D198A mutant PhuZ has a weak positioning bias toward midcell (orange). For all microscopy images (panels B,D,E), scale bar is 1 micron; magenta is membrane stain FM4-64, cyan is DNA stain DAPI, and green is GFP.

To determine whether PhuZ centers the nucleus in RAY infections, we expressed a GTP-hydrolysis deficient mutant, PhuZ_RAY_D198A as a dominant negative to inhibit the dynamic properties of the endogenous filaments and test whether the lack of filament dynamics interferes with nucleus positioning (Chaikeeratisak, Nguyen, Egan, et al., 2017; Erb et al., 2014; Kraemer et al.,2012). In uninfected and infected cells, GFP-PhuZ_RAY_D198A formed filaments at all arabinose concentrations, likely due to its inability to depolymerize (Fig. 6B,C,E). We measured the distance from the phage nucleus to the cell pole when GFP-PhuZ_RAY_ and GFP-PhuZ_RAY_D198A were expressed, taking into account whether the presence and location of bacterial nucleoids contributed to phage nucleus positioning. When a single bacterial nucleoid was present, the positioning of the RAY nucleus had a broad distribution with a slight bias toward midcell (Fig. 6F, blue bars). The single nucleoid in these cells frequently occluded the center of the cell, thereby preventing the nucleus from being centered (Fig 6D, cell iii, white arrow). When two bacterial nucleoids were present, the RAY nucleus was usually (97%, n=103) positioned between them near the cell midpoint (Fig. 3A; 6D cells i, ii, and iv; 6G). When PhuZ_RAY_D198A was expressed, the positioning of the RAY nucleus was less biased towards midcell when two nucleoids were present and had no specific localization when there was one nucleoid (Fig. 6F and G). This is the first time that phage nucleus positioning was measured in the presence of the bacterial genome, and these results suggest that, while the PhuZ filaments likely play a role in positioning the nucleus in phages that do not degrade host DNA, the phage nucleus must also compete for space with the bacterial DNA when it is not degraded.

In addition to centering the phage nucleus within the host cell, PhuZ also rotates the phage nucleus in both the ΦKZ-like phages and Goslar. We therefore performed time lapse microscopy and measured nucleus rotation in RAY-infected cells expressing GFP-ChmA_RAY_ (Fig. 3E, Movie 2). We observed very few of the phage nuclei actively rotating (3.7%, n = 858), compared to Goslar (97%) (Birkholz et al., 2022) and 201ϕ2-1 (46%) (Chaikeeratisak et al., 2019). Those that did rotated for 27.7 ± 5.8 seconds with an average linear rotation speed of 54.8 ± 15.6 nm/s and angular velocity of 10.1 ± 3.2 °/s (n = 25) (Fig. 3F). We also measured nucleus rotations using GFP-tagged major capsid protein gp317 (MCP_RAY_-GFP) docked on the surface of the nucleus as a fiduciary mark. The nuclei with docked capsids displayed a similar linear rotation speed of 57.5 ± 11.9 nm/s (n = 17). This rotation speed is only marginally faster when compared to Goslar (49.7 ± 12.5 nm/s) and 201ϕ2-1 (43.6 ± 7.6 nm/s) nucleus rotation speeds (Birkholz et al., 2022; Chaikeeratisak et al., 2019), suggesting that this rotation rate is conserved, possibly to allow the even spacing of capsids docking on the phage nucleus surface for efficient DNA encapsidation.

### *In situ* structural analysis of RAY phage replication in *Erwinia amylovora*

In order to investigate the structural conservation of chimallivirus replication components, we performed cryo-focused ion beam milling and electron tomography (cryoFIB-ET) of *Erwinia amylovora* cells infected with RAY (Fig. 7). The tomograms revealed the presence of the RAY phage nuclear shell as an irregularly shaped, protein-based compartment enclosing nucleic acid and excluding host ribosomes (Fig. 7A, B). Viral capsids were observed at different stages of maturation, docked on the phage nucleus where they package DNA, or in the cytoplasm with or without tails attached. We did not observe virion particles assembled into structured bouquets, in agreement with fluorescence microscopy (Fig. 7A, B).

**Figure 7.**
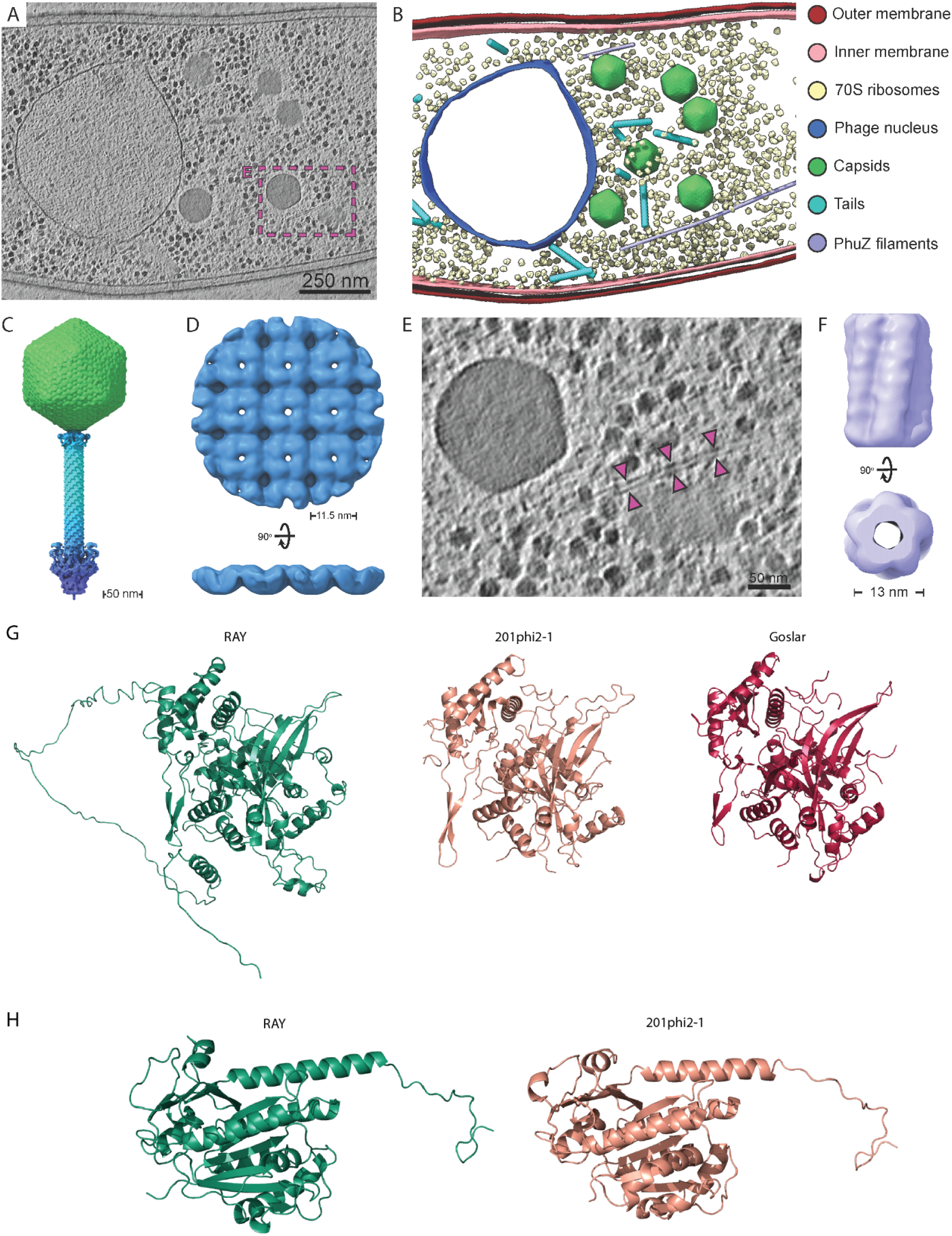
Cryo-electron tomography and structural analysis. (A) Slice through a cryo-electron tomogram of a RAY-infected *Erwinia amylovora* cell at approximately 105-110 mpi. Scale bar is 250 nm. (B) Segmentation of the tomogram in panel A. (C) Composite RAY virion reconstruction. (D) Orthogonal views of the RAY chimallin lattice reconstruction. (E) Enlarged view of region boxed in A with magenta arrows pointing to a putative RAY PhuZ filament. Scale bar is 50 nm. (F) Reconstruction of the putative RAY PhuZ filaments showing the five-stranded structure and hollow lumen. (G) AlphaFold v2.1.0 structure of RAY chimallin compared to experimentally determined 201ϕ2-1 and Goslar chimallin (Laughlin et al., 2022). (H) AlphaFold v2.1.0 structure of RAY PhuZ compared to experimentally determined 201ϕ2-1 PhuZ (Kraemer et al., 2012; Zehr et al., 2014). Accession numbers: C: EMD-28003 (additional map), D: EMD-28007, F: EMD-28008. See Figure S23 for subtomogram analysis workflows of RAY components. in the low-confidence placement of the extended N- and C-terminal segments, which mediate interactions across ChmA protomers in higher-order assemblies (Fig. S22).

To further assess the components of RAY replication, we performed subtomogram analysis of virion components, the phage nucleus shell, and PhuZ filaments. Through averaging of assembled and isolated virion components, we obtained reconstructions for the RAY capsid (~19 Å), collar (~38 Å), tail sheath (~10 Å), and baseplate (~36 Å). Dimensions of assembled virions measured from the tomograms and overlapping segments of the component maps allowed reconstruction of a composite RAY virion (Fig. 7C). The virion is largely similar to the composite structure of the isolated ΦKZ virion (Fokine et al., 2007). The RAY capsid exhibits a triangulation number of 27 (T = 27) and is approximately 142.5 nm in diameter, which is similar to the ΦKZ capsid. However, the RAY baseplate reconstruction appears in a retracted state, as opposed to the wider expanded state observed for the ΦKZ virion (Fokine et al., 2007).

Subtomogram analysis of the phage nuclear shell resulted in a ~20 Å reconstruction which revealed that the RAY nuclear shell is principally composed of a square chimallin lattice with ~11.5 nm repeat distance similar to that assembled by the 201ϕ2-1 and Goslar chimallin (Fig. 7D) (Laughlin et al., 2022). The conserved higher-order structure of the RAY chimallin lattice indicates a conserved underlying protomer structure. Indeed, the AlphaFold 2.0 predicted structure of RAY chimallin is largely consistent with the experimental chimallin structures determined for 201ϕ2-1 (RMSD: 2.2 Å) and Goslar (RMSD: 2.2 Å) (Laughlin et al., 2022). The ChmA_RAY_ protomer prediction predominantly differs from experimental structures

Finally, we performed subtomogram analysis (StA) of long, hollow, filamentous structures present in RAY-infected *Erwinia* cells assumed to be the phage-encoded tubulin, PhuZ (Fig. 7A,B,E). The resulting StA map achieved an estimated resolution of ~25 Å and revealed a hollow, five-stranded filament of approximately 13 nm in diameter (Fig. 7F), which is in stark contrast to the the three-stranded filaments made by PhuZ_201ϕ2-1_ obtained *in vitro* (Kraemer et al., 2012;Zehr et al., 2014). We did not observe other filamentous structures in our tomogram set, thus the five-stranded structures are tentatively assigned as RAY PhuZ. StA of PhuZ filaments observed in our previously published 201ϕ2-1 and ΦKZ cryoFIB-ET datasets revealed these PhuZ variants form three-stranded filaments in situ (Fig. S21B,C,D). The structure of the PhuZ_RAY_ protomer as predicted by AlphaFold is highly similar (RMSD = 1.069 Å) to the experimentally determined crystal structure of PhuZ_201ϕ2-1_ (Fig. 6H) (Kraemer et al.,2012; Zehr et al., 2014), thus indicating subtle differences are likely responsible for differences in higher-order assembly among PhuZ variants. Of note, residues R217:D305 and E225:R290 form essential salt-bridges for self-assembly of PhuZ_201ϕ2-1_ and are conserved in of PhuZ_ΦKZ_ (R230:D316 and D238:R301) but not PhuZ_RAY_ (D223:D315 and L235:D299). Further work will be necessary to unambiguously determine whether the five-stranded filaments are indeed PhuZ_RAY_ and the molecular bases of their higher-order assembly.

Taken together, the structures seen in the cryoFIB-ET corroborate the fluorescence microscopy results and confirm that RAY is a nucleus-forming phage that has unique and intriguing differences when compared with previously published Chimalliviridae Goslar, PCH45, and the ΦKZ-like phages.

## DISCUSSION

The phage nucleus is a remarkable and provocative cell biological structure, but the viral genes required for phage to replicate via this pathway are unknown. Here we bioinformatically identified the core genome that defines the nucleus-forming Chimalliviridae family. The Chimalliviridae core genome consists of a set of 72 genes encoded within seven distinct blocks. The order of the core blocks and genes within the blocks was generally conserved among divergent members of the family, as was their phylogeny. This suggests that these phages evolved from a common ancestor, and their genes have been co-evolving without significant horizontal gene transfer. We propose that the core genome identified here includes many of the key genes required for nucleus-based phage replication and that the less conserved accessory genes may be important for different subfamilies of phage or allow specialization for infecting different hosts. The core genes encode proteins that participate in many key processes such as genome replication, gene expression, and formation of virion particles. However, 74% of the core genes (53/72) are hypothetical proteins of unknown function and origin; moreover 29% (21/72) make up the core genes that are unique to the Chimalliviridae, and of these only chimallin has a known function (Chaikeeratisak, Nguyen, Khanna, et al., 2017; Laughlin et al., 2022). This analysis agrees well with previous studies that worked to identify a core geneset shared by different groups of jumbo phages (Iyer et al., 2021; Jang et al., 2013). This core geneset likely encodes numerous unknown components required for phage nucleus formation, standing in stark contrast to the core genome of the phage T4 family (Tequatroviridae), where the function of the majority of the core genes are known and genes of unknown function are primarily accessory (Comeau et al., 2007). Thus, the Chimalliviridae core genome is rich with potential for discovery of novel biological functions due to its numerous uncharacterized genes.

Phage can be classified into families (e.g. Ackermannviridae and Herelleviridae) based on genetic data (Barylski et al., 2020). Our phylogenetic analysis suggests that all phages encoding this core genome form a single family, the Chimalliviridae, that only arose once and developed a unique nucleus-based replication mechanism. Within this family are clades of phages with specific accessory genomes, suggesting that different Chimalliviridae clades have adapted the phage nucleus replication mechanism in a variety of diverse ways. The core genome of nucleus-forming phages described here can be used to study the basic requirements for nucleus-based replication, discover new members of the Chimalliviridae, and guide future studies in synthetic biology aimed at building a phage encoding only the minimal components required for phage nucleus-based replication.

We performed a detailed investigation into RAY to determine how conserved the phage nucleus replication mechanism is in a divergent Chimalliviridae family member. By examining the intracellular localization of fifteen core and nine non-core RAY proteins via GFP-tagging and fluorescence microscopy, including chimallin, we discovered that these proteins have conserved subcellular localizations across Chimalliviridae, with DNA processing proteins being found inside the phage nucleus, virion proteins being found around the periphery of the phage nucleus, and other proteins being found in the cytoplasm (Fig. 3, 4, 5, 6) (Birkholz et al., 2022; Chaikeeratisak et al., 2019; Chaikeeratisak, Nguyen, Egan, et al., 2017; Chaikeeratisak, Nguyen, Khanna, et al., 2017; Malone et al., 2020; Nguyen et al., 2021). We show that chimallin surrounds the phage DNA, and cryo-FIB-ET revealed it assembles an enclosed square lattice, forming a 6 nm thick shell that separates phage DNA from ribosomes and virion structural proteins in the cytoplasm, similarly to 201ϕ2-1, ΦKZ, ΦPA3, and Goslar (Birkholz et al.,2022; Chaikeeratisak et al., 2019; Chaikeeratisak,Nguyen, Egan, et al., 2017; Chaikeeratisak, Nguyen,Khanna, et al., 2017; Nguyen et al., 2021). The putative functions of these proteins have been updated in the NCBI annotation (NC_041973.1).

We also found several additional surprises (unconserved processes) that add to our knowledge of the diversity of Chimallivirus replication. First, we found that the host chromosome is not degraded and is excluded from the phage nucleus. This has only previously been observed with phage PCH45 (Malone et al., 2020) and suggests that the host chromosome is not a barrier to building a phage nucleus. Second, we found that the PhuZ spindle and host chromosomes both likely impact phage nucleus position. Third, we found that RAY PhuZ assembles five stranded tubes with a central lumen, which has never before been observed in any prokaryotic tubulin. This is in contrast with ΦKZ-like *Pseudomonas* phages, where PhuZ assembles triple-stranded polymers. Fourth, unlike *Pseudomonas* and *E. coli* nucleus forming phages, whose nuclei are more consistently rotated by the PhuZ spindle, we rarely observed nucleus rotation with RAY. Finally, unlike *Pseudomonas* and *E. coli* phages, RAY viral particles do not form bouquets.

We described the characterization of ten proteins that had not previously been studied in any chimallivirus. For example, we discovered one core and one accessory protein of unknown function that both contain a domain belonging to the general family of SWI/SNF helicases, but nothing is known about them, so it is difficult to predict what their functions in the phage life cycle might be. Surprisingly, we found one of these proteins localized inside the nucleus while the other one localized inside the capsids. Another example is a phage-encoded DNA-binding protein that binds to bacterial DNA. This protein is not part of the core genome and seems to be one of the accessory proteins that differentiates RAY from some of the previously studied nucleus-forming phages since this protein would not have a useful function in a phage that degrades the host genome. Bioinformatic analysis predicts this protein is an XRE family transcriptional repressor, but it is difficult to predict its biological function. Other proteins described here include a protease (gp116), a ClpXP protease adapter protein (gp039), and a tRNA ligase (gp048), all of which could potentially be involved in overcoming host defense systems.

Our results suggest a model for RAY replication that is distinct from other previously characterized nucleus-forming phages (Fig. 8). Upon infection, RAY injects its DNA and a suite of proteins for viral replication, including four vRNAP proteins, as well as proteins of unknown function (such as putative SF2 helicase gp131 and internal head protein gp299). The injected RNA polymerase expresses chimallin (Ceyssens et al., 2014), which forms a proteinaceous shell surrounding the phage DNA (Fig. 3B). Unlike with 201ϕ2-1, ΦKZ, ΦPA3, and Goslar, the host chromosome is not degraded, similar to PCH45 infection (Malone et al., 2020), and the cell does not undergo substantial swelling. The growing phage nucleus therefore must compete for limited space with the chromosomal DNA. If the cell contains two well separated nucleoids, the nucleus usually localizes between them (Fig. 3B; 6D,E; Movie 1). Whether RAY does not cause significant cell bulging or degrade the chromosome because it does not encode genes that could cause this to occur or because the host encodes defense mechanisms that counter these processes is unknown. Capsids dock on the phage nucleus (Fig. 5A,B; 7A,B), package DNA, and eventually mature viral particles form (Fig. 8). PhuZ_RAY_ forms filaments that appear to contribute to RAY nucleus positioning (Fig. 6F,G), but whether they contribute to capsid trafficking as speculated in Figure 8 is unknown. It is also unclear if they contribute to nuclear rotation: Since the RAY nucleus was only observed to rotate in ~3.7% of infected cells, it was not possible to measure a significant difference between expressing live versus catalytically dead PhuZ. After assembly of mature phage particles, RAY does not form bouquets (Fig. 7A,B). Given RAY’s inability to degrade the chromosome or cause the cell to bulge, there may be no room for such large and elaborate structures to assemble.

**Figure 8.**
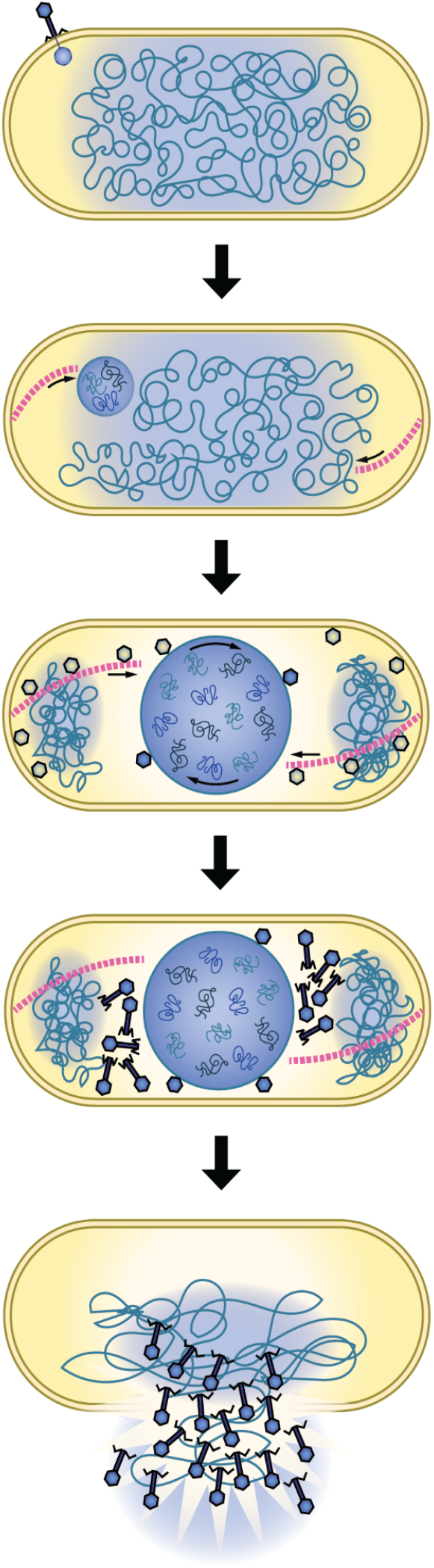
RAY infection model. RAY injects its DNA into the host cell. A nascent phage nucleus encloses it and is pushed toward midcell by the PhuZ filaments. The bacterial DNA is pushed out of the way as the phage nucleus expands, the phage DNA is replicated, and capsids are trafficked along PhuZ filaments. Capsids dock on the phage nucleus to fill with DNA before detaching and acquiring tails. At the end of RAY’s infection cycle, the cell lyses and progeny phage are released and may infect new hosts.

Fascinatingly, the phage-encoded tubulin homolog PhuZ is absent from one third of known Chimalliviridae yet displays conserved function in phage nucleus rotation in 201ϕ2-1, ΦKZ, ΦPA3, and Goslar (Birkholz et al., 2022; Chaikeeratisak et al., 2019), and in nuclear positioning in 201ϕ2-1, ΦKZ, ΦPA3, and RAY (Chaikeeratisak, Nguyen,Egan, et al., 2017; Chaikeeratisak, Nguyen, Khanna,et al., 2017; Erb et al., 2014; Kraemer et al., 2012) (Fig. 6F,G). However, unlike previously studied PhuZ proteins for 201ϕ2-1, ΦKZ, and ΦPA3 which form three-stranded filaments (Zehr et al., 2014), PhuZ_RAY_ appears to form five-stranded filaments which make PhuZ_RAY_ the first phage-encoded tubulin-based structure that possesses a lumen (Fig. 7F, Fig. S21). This is not only distinct from other PhuZ homologs, but also from most bacterial tubulins, which assemble either single filaments (FtsZ) (Du & Lutkenhaus, 2019; Erickson et al.,2010; Haeusser & Margolin, 2016) or two or four stranded filaments (TubZ) (Aylett et al., 2010;Larsen et al., 2007; Montabana & Agard, 2014). However, it is similar to *Prosthecobacter* tubulins BtubA/BtubB, which assemble five-stranded tubules *in vivo*, and a variety of single, double and polymeric structures *in vitro* and are more similar to eukaryotic tubulins than bacterial tubulins, likely arriving in bacteria through horizontal gene transfer (Díaz-Celis et al., 2017; Schlieper et al., 2005; Yee et al., 2007). PhuZ is an example of an accessory protein that has likely adapted to make different types of cytoskeletal structures for different phages. Among the subset of Chimalliviridae we have examined thus far, we have observed PhuZ to form a bipolar spindle or a vortex, and either three-stranded filaments or five-stranded tubes. Additionally, prior studies have reported only a 50% decrease in phage titers when expressing dominant negative PhuZ mutants despite severe phage nucleus positioning defects (Chaikeeratisak,Nguyen, Egan, et al., 2017; Erb et al., 2014;Kraemer et al., 2012). In agreement with these studies, the PhuZ gene can be completely deleted from ΦKZ, and while phage nucleus position is again strongly affected, the phage is still viable (Guan et al., 2022). Taken together, the lack of universal conservation among the Chimalliviridae, the small effect of dominant negative mutants on phage titer, and the knockout experiments showing PhuZ is non-essential suggest that PhuZ is an accessory protein that may confer improved infection efficiency in some nucleus-forming phages, but it is not essential for this viral replication pathway.

Many phages package proteins within their capsids that they are thought to inject into the cell along with their DNA (Falco et al., 1980; Jin et al.,2015; Molineux & Panja, 2013; Sokolova et al.,2020; Thomas et al., 2012). These proteins have previously been discovered by performing mass spectrometry on purified viral particles (Thomas et al., 2012). Using GFP fusions, we have shown localizations consistent with virion packaging of these proteins, corroborating previous mass spectroscopy data and showing that visualization of internal capsid proteins is possible using fluorescence microscopy. For example, the virion RNA polymerase subunits (vRNAP) localized around the phage nucleus where the capsids accumulated, in agreement with the previous mass spectrometry data that these vRNAPs are encapsidated (Thomas et al., 2012). So did gp131, which was also previously found to be virion associated and is predicted to be an SF2 superfamily helicase. We speculate that gp131 may be responsible for organizing the phage DNA inside the capsid, similar to ΦKZ gp203 (Thomas et al., 2012), or that it could be injected along with DNA and function as a helicase involved in early gene transcription along with the vRNAP.

This work highlights how, despite the unique features of each nucleus-forming phage studied to date, many of the basic components are conserved across the entire diverse family of Chimilliviridae. Here we document differences in phage tubulin assembly, nucleus positioning, nucleus rotation, chromosome degradation, and cell bulging that likely represent just a few of the many phenotypic consequences that result from continual phage-host evolutionary conflicts. Many of these differences are likely due to the continually evolving accessory genome. Now that the core and accessory genomes of this unique phage family have been identified, future studies can be better directed toward understanding their functions in order to elucidate the intricacies of the nucleus-based phage replication mechanism

## MATERIALS AND METHODS

### Genome Alignments and Phylogenetic Analysis

To find close and distant relatives of RAY, we used ViPTree (Nishimura et al., 2017) to find phage genomes related to RAY (Fig. S24) and PSI-BLAST to find phage encoding chimallin homologs (Table 1) and msRNAP homologs since all chimallin-encoding phages also encode msRNAPs (Iyer et al., 2021; Jang et al., 2013; Sokolova et al., 2020) PSI-BLAST iterations were performed until no new sequences were found above the 0.005 E-value threshold (for chimallin, the highest E-value in the results was 1e-08). These results were checked for validity by multiple sequence alignments with previously identified homologs to ensure they were not false positives. *Bacillus* phage PBS1, *Staphylococcus* phage PALS_2, and *Yersinia* phage ΦR1-37 were used as the representative phages that encode msRNAPs but not chimallin homologs, and well-studied *Escherichia* phage T4, giant *Bacillus* phage G, and *Erwinia* phage vB_EamM_Alexandra were also included in our analysis for comparison. VICTOR (Meier-Kolthoff & Göker, 2017) predicted the species assignments, which were used to determine which phage could be discluded to keep the trees readable; genus assignments, which were used to color-code the trees; and subfamily and family assignments, which were used along with core genome analysis to define the family Chimalliviridae (Table 2). Phylogenetic trees were made using VICTOR (Fig. 1A) and colored using Photoshop (Adobe) or Clustal Omega (Fig. 1B-D, S1, S2, S21) and colored using iTOL (Letunic & Bork, 2021).

### Core Genome Determination

Bulk PSI-BLAST in the standalone BLAST+ Suite (Altschul et al., 1997) was performed on the tailed phages taxon (taxid: 28883) with RAY proteins as the query, a max iteration of 4, and an e-value cutoff of 0.5. Then two scripts were run on the result to search for proteins from the phages of interest as well as output the ones present in more than 90% of the phages through a text mining approach. Scripts are available on GitHub (https://github.com/jina-leemon/core-genome-proj). Blocks were defined as 3 or more conserved genes, tolerating a maximum of 3 non-conserved genes between. The core genome numbers (Table 3) were assigned depending on the homologs that showed up through PSI-BLAST in ΦKZ, Goslar, and RAY.

### Bacterial Growth and Phage Infection Conditions

*Erwinia amylovora* ATCC 29780 was grown on LB plates at room temperature or in LB liquid cultures at 30°C. To collect RAY lysates, 0.5 mL of dense *E. amylovora* culture grown in liquid LB media at 30°C overnight was incubated with 10 μL of RAY serial dilutions, incubated for 15 minutes at room temperature, and mixed with 4.5 mL molten (approximately 55°C) LB 0.35% top agar. This mixture was quickly poured over standard LB plates and incubated overnight at room temperature. The following day, 5 mL phage buffer was poured over plates showing web lysis and incubated at room temperature for 5 hours. The phage lysate was collected by aspiration and centrifuged for 10 min at 3220 rcf to pellet cell debris. The resulting clarified phage lysate was filtered through a 0.45 μm filter to remove any residual bacterial contamination and stored at 4°C.

### Expression of Phage Proteins

To visualize phage proteins in *Erwinia amylovora,* the proteins were fused to GFPmut1 and expressed from the pHERD-30T plasmid under the inducible control of the AraBAD promoter. GFP fusions were designed using Benchling with the GFP tag attached at either the N- or C-terminus depending on the protein. The plasmids were then synthesized by GenScript. Plasmids were transformed into electrocompetent *Erwinia amylovora* cells via electroporation. The cells were plated on LB plates with 15 μg/mL gentamicin sulfate as a selectable marker for transformants.

### Phage Protein Characterization

PSI-BLAST was used in order to identify potential homologs of each unknown RAY protein (Altschul et al., 1997). The PSI-BLASTs were performed with the non-redundant protein database, limited to tailed phages (taxid: 28883), excluding uncultured and environmental sample sequences, and with a maximum of 5000 sequences. Iterations were run excluding results with e-values lower than the default PSI-BLAST threshold, stopping when results converged. For each protein studied, Phyre^2^ was used in order to predict secondary structure and to identify known proteins with similar structures (Kelley et al., 2015). The amino acid sequences of each protein were uploaded to Phyre^2^ with the normal modeling mode. Phyre^2^ results with confidence higher than 70% were compared to PSI-BLAST results as an independent method for predicting the putative functions of unannotated RAY proteins. Using the potential homologs of each unknown RAY protein identified by PSI-BLAST, multiple sequence alignments were created using Clustal Omega (Madeira et al., 2022) to align homologs from previously studied nucleus forming phages 201ϕ2-1, ΦKZ, ΦPA3, PCH45, and Goslar. If one or more of these phages were missing a homolog of the query protein (as was the case with some of the non-core genome proteins), a close relative of RAY *(Erwinia* phage AH06) and/or a distant relative of RAY *(Klebsiella* phage Miami) were supplemented to fill out the alignment when possible. The resulting multiple sequence alignments were uploaded to ESPript for visualization using the default settings and downloaded as PDF files (Robert & Gouet, 2014).

### Fluorescence Microscopy

*E. amylovora* cells were inoculated onto imaging pads in welled microscopy slides. Pads were made up of 1% agarose, 25% LB, 2 μg/mL FM4-64, and 0.1 μg/mL DAPI. Between 0-1% arabinose was used to induce expression from the pHERD-30T plasmid, depending on the construct. The slides were incubated at 30°C for 3 hours, then moved to room temperature for infection with 10 μL undiluted RAY lysate. Slides were imaged using the DeltaVision Elite deconvolution microscope (Applied Precision) and deconvolved using the aggressive algorithm in the DeltaVision softWoRx program (Applied Precision). Image analysis was performed on images prior to deconvolution.

### Nucleus Rotation Analysis

Time lapses (2 minutes long with 4-second intervals) were obtained, generally between 60-90 mpi. The time lapses that included actively rotating nuclei were analyzed using FIJI. The segmented line tool was used to measure all distances. To determine the nucleus rotation speed, a distinct point on the surface of the nucleus was tracked frame by frame to measure the total distance traveled. The number of frames used and the time lapse interval size were used to determine the rotation time. The rotation speed was then calculated by dividing the total distance traveled by the rotation time. The total number of cells with nuclei and the number of nuclei that were actively rotating were manually quantified to determine the percentage of rotating nuclei. The angular velocity was determined by dividing the total distance traveled by the length of the nucleus’ radius (half of the nucleus’ diameter).

### PhuZ Filamentation and Nucleus Positioning Analysis

For the filamentation analysis, the number of cells with filaments versus without filaments was manually quantified from microscopy images, and the percentage of cells with filaments at different arabinose concentrations was recorded. A filament was defined as a visible line that was at least double the level of background GFP fluorescence. The data were graphed using Prism (GraphPad), and a nonlinear [Agonist] vs. response curve was fit to represent the [Arabinose] vs. filamentation data.

For the nucleus positioning analysis, the length of the cell and the distance between the center of the phage nucleus and the cell pole was measured using FIJI. The nucleus position was calculated as the ratio of the distance from the phage nucleus to the pole over the entire cell length. The data were graphed as a relative frequency histogram using Prism (GraphPad).

### Cryo-electron Microscopy of RAY-infected Cells

*E. amylovora* cells were infected on agarose pads as previously described (Chaikeeratisak, Nguyen, Khanna, et al., 2017) with a few changes. Briefly, cells were grown on agarose pads as described above for fluorescence microscopy and infected at room temperature with 10 μL undiluted RAY lysate. 60 minutes post infections, 25 μL 25% LB was added to each pad and cells were gently scraped off with the bottom of a 1.7 mL eppendorf tube. A droplet containing cells was collected from each pad and an aliquot was saved as an unconcentrated sample. The remainder was centrifuged at 8000 rcf for 30 sec, and resuspended in a portion of the supernatant to concentrate the cells.

Infected cell suspension was mixed 9:1 with 50% (w/v) trehalose solution to mitigate crystalline ice formation and 4 μL of this mixture immediately applied to a R2/1 Cu 200 grid (Quantifoil), which had been glow-discharged for 1 min at 0.19 mbar and 20 mA in a PELCO easiGlow device shortly before use. The grid was then mounted in a custom-built manual plugning device (MPI-Biochemistry, Martinsreid, Germany) and excess media wicked away with filter paper (Whatman #1) from the backside of the grid for 4-7 seconds prior to plunging the grid into a 50:50 ethane:propane mixture (Airgas) cooled by liquid nitrogen. The data presented in this manuscript were collected from two independent preparations of RAY-infected *E. amylovora* at approximately 100-110 mpi.

Frozen grids of infected cells were mounted into notched Autogrids (TFS) compatible with cryo-focused ion beam milling. Samples were loaded into an Aquilos 2 cryo-focused ion beam/scanning electron microscope (TFS) with a Gallium ion source and milled to generate lamellae using progressively lower milling currents from 0.5 nA to 10 pA as previously described (Lam & Villa,2021).

For tilt-series collection, samples were transferred to a Titan Krios G3 transmission electron microscope (TFS) operated at 300 kV, configured for fringe-free illumination, and equipped with a K2 direct electron detector (Gatan) mounted post a Quantum 968 LS imaging filter (Gatan). The microscope was operated in EFTEM mode with a slit-width of 20 eV and using a 70 μm objective aperture. Automated data acquisition was performed using SerialEM-v3.8.7 (Mastronarde, 2005) and all images were acquired using the K2 in counting mode. Tilt-series were acquired at 4.27 Å/pixel following a dose-symmetric scheme to span a nominal range of either +/- 57° (dataset-1) or +/- 64° (dataset-2) in 2° steps. The exposure was uniformly distributed across tilt-images, achieving totals of approximately 150 e/Å^-2^ (dataset-1) or 170 e/Å^-2^ (dataset-2). Target defoci for tilt-series were between 4-6 μm. A total of 32 tilt-series were acquired, of which 26 (13 from each session) were deemed suitable for subsequent processing.

### Subtomogram Analysis

All pre-processing steps were performed using Warp-v1.09 (Tegunov & Cramer, 2019) unless otherwise specified. Tilt-movies were corrected for whole-frame motion, their defocus values estimated, and stacked into tilt-series. Tilt-series were aligned via patch-tracking with Etomo (IMOD-v4.10.28) (Kremer et al., 1996). Tomograms were reconstructed with the default deconvolution filter settings for visualization and without for template-matching and subsequent processing, when necessary.

First, an *ab initio* host 70S ribosome reference was generated from the data by manually picking particles and aligning them in RELION-v3.1.3 (Bharat et al., 2015; Scheres, 2012). A low-pass filtered (50 Å) reference was used for template-matching across the entire dataset and the hits curated by Figure-Of-Merit score to remove false-positives. The 51,512 template-matching hits were classified in RELION-v3.1.3 without a starting reference to further remove false positives. Alignment of the 28,032 selected particles resulted in a 18 Å map of the 70S ribosome. Further refinement of particle poses and tilt-series parameters in M-v1.09 (Tegunov et al., 2021) improved the map to 10.3 Å. This map was low-pass filtered and used for a second round of template-matching against the tomograms, which resulted in 74,713 hits after curation. The new particle set was coarsely aligned with RELION-v3.1.3 and followed by M-v1.0.9, which resulted in a 9.5 Å map. Further alignment with RELION-v3.1.3 improved the map to 9.2 Å. Reference-free classification without alignment in identified 45,708 70S particles which refined to 8.9 Å and 2,551 50S particles which refined to 13.4 Å.

For virion capsids, 156 particle positions were manually picked and aligned while enforcing icosahedral symmetry in RELION-v3.1.3. The vertices were subsequently extracted to create 1,872 sub-particles, which aligned with C5 symmetry and then classified in C1 without a starting reference nor additional alignment. The 1,056 selected particles were further classified without a starting reference using an ellipsoid mask, C6 symmetry with relaxed C5 symmetry, and local-searches to separate pentameric vertices from the portal vertices. The 1,023 pentameric vertices were further aligned with C5 symmetry to yield a 19 Å map. The 22 portal vertices were aligned with C6 symmetry to yield a 38 Å map.

For virion tails, the start and end points of nine tails exhibiting clear polarity in the tomograms were picked and filament cropping models generated using Dynamo-v1.1514 (Navarro et al., 2020). Subtomograms were extracted every 2 nm to yield 900 segments and the azimuth angles randomized. An initial reference was generated by reconstruction of the segments and smoothing of the resulting map using a Gaussian filter of 3 pixel width. The smoothed map was used to align all segments without enforced symmetry and restricted angular search range. The resulting map exhibited clear polar and C6 symmetry. Next, the start and end points of all tails were picked, filament models generated, and subtomograms extracted every 2 nm with randomized azimuth angles. A copy of the metadata table was generated in which the polarity of each filament was flipped. Both tables were aligned for a single iteration against the previously generated reference with C6 symmetry enforced while limiting the alignment to 40 Å. Duplicate particles which had converged on the same position due to the initial oversampling and subsequent alignment step were removed. In order to determine the polarity of segments, the cross-correlation (CC) values for analogous tails between the two tables were compared. For each tail, the orientation leading to the higher median CC across all its segments was chosen for subsequent steps. For tails exhibiting similar, low median CC values in both orientations were discarded. A CC threshold was selected for the entire dataset to remove these ambiguous tails and other low CC segments. This was typical of tails only partially contained within the lamellae, for which a majority was either FIB-ablated or extended outside of the field-of-view. The resulting 3,291 segments were converted for processing in RELION-v3.1.3 using dynamo2m-v0.2.2 (Burt et al., 2021) and halfsets were assigned on a per-tail basis. Refinement in RELION-v3.1.3 while enforcing C6 symmetry resulted in a 10.4 Å map. Reference-free classification with C6 symmetry and without alignment separated tail segments from baseplates. Alignment of the selected 3,257 tails segments with C6 symmetry resulted in a 10 Å map which exhibits a rise of 37.4 Å and twist of 22.3°. The 34 baseplates were aligned with C6 symmetry to yield 36 Å map.

For ChmA, the surfaces of the twenty phage nuclei were coarsely contoured to generate surface cropping models with Dynamo-v1.1514. Subtomograms were extracted every 4 nm and oriented normal to the surface models. An initial reference was generated by reconstruction of the segments and smoothing of the resulting map using a Gaussian filter of 3 pixel width. A subset of 3,910 segments from two tomograms were aligned against the smoothed map with a restricted angular search range to prevent flipping of sidedness and without enforced symmetry. The resulting map exhibited apparent p442 lattice symmetry with an approximate 11.5 nm spacing, which was corroborated by inspection of ‘neighbor plots’ (Kovtun et al., 2018; Pyle et al., 2022). This new reference was C4 symmeterized and used to align the entire dataset of 123,416 oversampled segments for a single iteration while limiting alignment to 40 Å. Duplicate particles which had converged on the same position due to the initial oversampling and subsequent alignment step were removed to leave 52,646 segments. Misaligned and edge segments were then removed by selecting only segments with at least three neighboring segments with 10 to 13 nm, which retained 23,509 segments. The metadata was converted for processing in RELION-v3.1.3 using dynamo2m-v0.2.2. Further alignment in RELION-v3.1.3 enforcing C4 symmetry yielded a 20 Å map.

For PhuZ, the start and end points of 14 filaments were manually picked from the tomograms and cropping models generated with Dynamo-v1.1514. The polarity of the filaments was uniformly assigned from the cell pole to the mid-cell. Segments were extracted every 1 nm along the filaments and the azimuth angles were randomized, which resulted in 5,000 subtomograms. Sub-tomograms were aligned to the unaligned average reconstructed from the initial cropping points with Dynamo-v1.1514 while limiting the alignment resolution to 40 Å and preventing subtomograms from flipping polarity. Duplicate particles which had converged on the same position due to the initial oversampling and subsequent alignment step were removed, which resulted in 3,080 remaining subtomograms. The metadata was converted for processing in RELION-v3.1.3 using dynamo2m-v0.2.2 and halfsets were assigned on a per-filament basis. Alignment in RELION-v3.1.3 resulted in a 25 Å map. Individual protomers are not resolved in this map and attempts at estimating reliable helical parameters were unsuccessful.

All resolution estimates are based on the 0.143-threshold of the Fourier shell correlation between masked half-maps using high-resolution noise-substitution to mitigate masking artifacts (Chen et al., 2013). Local-resolution estimates were commuted using RELION. The virion ‘frakenmap’ was generated by the *vop maximum* command in UCSF-Chimera-v1.15 (Goddard et al., 2007) to combine the various components for display purposes. Maps were rendered using ChimeraX-v1.3 or v1.4 (Goddard et al., 2018).

### Tomogram Segmentation

To aid in segmentation and for display purposes, tomograms were missing-wedge corrected using IsoNet-v0.1 (Liu et al.,n.d.) on tomograms reconstructed at 20 Å/px and devolved with Warp-v1.09. Membranes were initially segmented using TomoSegMemTV (Martinez-Sanchez et al., 2014) and patched manually using Amira3D-v2021.2 (TFS). Subtomogram averages were placed at refined particle positions using Dynamo-v1.1514. The segmentation was rendered using ChimeraX-v1.3 (Goddard et al.,2018).

### Structural Predictions

AlphaFold v2.1.0 (Jumper et al., 2021) was used in order to predict the tertiary structure of unsolved RAY proteins. We ran ChmA_RAY_ (gp222) and PhuZ_RAY_ (gp210) through AlphaFold v2.1.0 to obtain the predicted structure of the proteins. These predictions were then imported into PyMOL Version 2.5.2 (Schrödinger, LLC, 2021) in order to visualize the structures. Structural alignments between previously published protein structures and RAY protein structural predictions were made using PyMOL’s align function, and the RMSD values of these alignments were found using the same method.

## DATA AVAILABILITY

Scripts used for core genome determination PSI-BLAST are available on GitHub (https://github.com/jina-leemon/core-genome-proj).

Tilt-series frames and alignment metadata for the RAY-infected *E. amylovora* cells are deposited with the Electron Microscopy Public Image Archive with accession number EMPIAR-11198. All subtomogram averaging maps from this study are deposited with the Electron Microscopy Data Bank with the following accession numbers: *E. amylovora* 70S (EMD-27973), *E. amylovora* 50S (EMD-27993), RAY capsid vertex (EMD-28003), RAY collar (EMD-28004), RAY tail sheath (EMD-28005), RAY baseplate (EMD-28006), RAY chimallin (EMD-28007), putative RAY PhuZ (EMD-28008), 201ϕ2-1 PhuZ (28009), and ΦKZ PhuZ (28010). The composite RAY virion map is deposited as an additional map with the RAY capsid vertex (EMD-28003).

## ACKNOWLEDGMENTS

We thank members of the Pogliano, Villa, Grose, Meyer, Dutton, and Corbett laboratories for helpful discussions and feedback. We thank the Grose laboratory for graciously providing the original *Erwinia* phage RAY isolate this work was based on. We thank B. Dennis and K. Smith of UCSD Physics Computing for computational support. Electron microscopy data were collected at the UCSD Cryo-Electron Microscopy Facility, which was built and equipped with funds from UCSD and an initial gift from the Agouron Institute. The authors acknowledge funding from the National Institutes of Health grant R01 GM129245 (to J.P. and E.V.) as well as from the National Science Foundation grant DBI 1920374 (to E.V.). T.L. is a Simons Foundation Awardee of the Life Sciences Research Foundation. E.V. is a Howard Hughes Medical Institute Investigator.

The authors declare that they have no conflict of interest.

## REFERENCES

Altschul, S. F., Madden, T. L., Schäffer, A. A., Zhang, J., Zhang, Z., Miller, W., & Lipman, D. J. (1997). Gapped BLAST and PSI-BLAST: a new generation of protein database search programs. Nucleic Acids Research, 25(17), 3389–3402.

Arens, D. K., Brady, T. S., Carter, J. L., Pape, J. A., Robinson, D. M., Russell, K. A., Staley, L. A., Stettler, J. M., Tateoka, O. B., Townsend, M. H., Whitley, K. V., Wienclaw, T. M., Williamson, T. L., Johnson, S. M., & Grose, J. H. (2018). Characterization of two related Erwinia myoviruses that are distant relatives of the PhiKZ-like Jumbo phages. PloS One, 13(7), e0200202.

Aylett, C. H. S., Izoré, T., Amos, L. A., & Löwe, J. (2013). Structure of the tubulin/FtsZ-like protein TubZ from Pseudomonas bacteriophage ΦKZ. Journal of Molecular Biology, 425(12), 2164–2173.

Aylett, C. H. S., Wang, Q., Michie, K. A., Amos, L. A., & Löwe, J. (2010). Filament structure of bacterial tubulin homologue TubZ. Proceedings of the National Academy of Sciences of the United States of America, 107(46), 19766–19771.

Barylski, J., Enault, F., Dutilh, B. E., Schuller, M. B., Edwards, R. A., Gillis, A., Klumpp, J., Knezevic, P., Krupovic, M., Kuhn, J. H., Lavigne, R., Oksanen, H. M., Sullivan, M. B., Jang, H. B., Simmonds, P., Aiewsakun, P., Wittmann, J., Tolstoy, I., Brister, J. R., … Adriaenssens, E. M. (2020). Analysis of Spounaviruses as a Case Study for the Overdue Reclassification of Tailed Phages. Systematic Biology, 69(1), 110–123.

Bharat, T. A. M., Russo, C. J., Löwe, J., Passmore, L. A., & Scheres, S. H. W. (2015). Advances in Single-Particle Electron Cryomicroscopy Structure Determination applied to Sub-tomogram Averaging. In Structure (Vol. 23, Issue 9, pp. 1743–1753). https://doi.org/10.1016/j.str.2015.06.026

Birkholz, E. A., Laughlin, T. G., Armbruster, E., Suslov, S., Lee, J., Wittmann, J., Corbett, K. D., Villa, E., & Pogliano, J. (2022). A cytoskeletal vortex drives phage nucleus rotation during jumbo phage replication in E. coli. Cell Reports, 40(7), 111179.

Burt, A., Gaifas, L., Dendooven, T., & Gutsche, I. (2021). A flexible framework for multi-particle refinement in cryo-electron tomography. PLoS Biology, 19(8), e3001319.

Cazares, A., Mendoza-Hernández, G., & Guarneros, G. (2014). Core and accessory genome architecture in a group of Pseudomonas aeruginosa Mu-like phages. BMC Genomics, 15, 1146.

Ceyssens, P.-J., Minakhin, L., Van den Bossche, A., Yakunina, M., Klimuk, E., Blasdel, B., De Smet, J., Noben, J.-P., Bläsi, U., Severinov, K., & Lavigne, R. (2014). Development of giant bacteriophage ϕKZ is independent of the host transcription apparatus. Journal of Virology, 88(18), 10501–10510.

Chaikeeratisak, V., Birkholz, E. A., Prichard, A. M., Egan, M. E., Mylvara, A., Nonejuie, P., Nguyen, K. T., Sugie, J., Meyer, J. R., & Pogliano, J. (2021). Viral speciation through subcellular genetic isolation and virogenesis incompatibility. Nature Communications, 12(1), 342.

Chaikeeratisak, V., Khanna, K., Nguyen, K. T., Egan, M. E., Enustun, E., Armbruster, E., Lee, J., Pogliano, K., Villa, E., & Pogliano, J. (2022). Subcellular organization of viral particles during maturation of nucleus-forming jumbo phage. Science Advances, 8(18), eabj9670.

Chaikeeratisak, V., Khanna, K., Nguyen, K. T., Sugie, J., Egan, M. E., Erb, M. L., Vavilina, A., Nonejuie, P., Nieweglowska, E., Pogliano, K., Agard, D. A., Villa, E., & Pogliano, J. (2019). Viral Capsid Trafficking along Treadmilling Tubulin Filaments in Bacteria. Cell, 177(7), 1771–1780.e12.

Chaikeeratisak, V., Nguyen, K., Egan, M. E., Erb, M. L., Vavilina, A., & Pogliano, J. (2017). The Phage Nucleus and Tubulin Spindle Are Conserved among Large Pseudomonas Phages. Cell Reports, 20(7), 1563–1571.

Chaikeeratisak, V., Nguyen, K., Khanna, K., Brilot, A. F., Erb, M. L., Coker, J. K. C., Vavilina, A., Newton, G. L., Buschauer, R., Pogliano, K., Villa, E., Agard, D. A., & Pogliano, J. (2017). Assembly of a nucleus-like structure during viral replication in bacteria. Science, 355(6321), 194–197.

Chen, S., McMullan, G., Faruqi, A. R., Murshudov, G. N., Short, J. M., Scheres, S. H. W., & Henderson, R. (2013). High-resolution noise substitution to measure overfitting and validate resolution in 3D structure determination by single particle electron cryomicroscopy. In Ultramicroscopy (Vol. 135, pp. 24–35). https://doi.org/10.1016/j.ultramic.2013.06.004

Comeau, A. M., Bertrand, C., Letarov, A., Tétart, F., & Krisch, H. M. (2007). Modular architecture of the T4 phage superfamily: a conserved core genome and a plastic periphery. Virology, 362(2), 384–396.

de Martín Garrido, N., Orekhova, M., Lai Wan Loong, Y. T. E., Litvinova, A., Ramlaul, K., Artamonova, T., Melnikov, A. S., Serdobintsev, P., Aylett, C. H. S., & Yakunina, M. (2021). Structure of the bacteriophage PhiKZ non-virion RNA polymerase. Nucleic Acids Research, 49(13), 7732–7739.

Díaz-Celis, C., Risca, V. I., Hurtado, F., Polka, J. K., Hansen, S. D., Maturana, D., Lagos, R., Mullins, R. D., & Monasterio, O. (2017). Bacterial Tubulins A and B Exhibit Polarized Growth, Mixed-Polarity Bundling, and Destabilization by GTP Hydrolysis. Journal of Bacteriology, 199(19). https://doi.org/10.1128/JB.00211-17

Du, S., & Lutkenhaus, J. (2019). At the Heart of Bacterial Cytokinesis: The Z Ring. Trends in Microbiology, 27(9), 781–791.

Erb, M. L., Kraemer, J. A., Coker, J. K. C., Chaikeeratisak, V., Nonejuie, P., Agard, D. A., & Pogliano, J. (2014). A bacteriophage tubulin harnesses dynamic instability to center DNA in infected cells. eLife, 3. https://doi.org/10.7554/eLife.03197

Erickson, H. P., Anderson, D. E., & Osawa, M. (2010). FtsZ in bacterial cytokinesis: cytoskeleton and force generator all in one. Microbiology and Molecular Biology Reviews: MMBR, 74(4), 504–528.

Esplin, I. N. D., Berg, J. A., Sharma, R., Allen, R. C., Arens, D. K., Ashcroft, C. R., Bairett, S. R., Beatty, N. J., Bickmore, M., Bloomfield, T. J., Brady, T. S., Bybee, R. N., Carter, J. L., Choi, M. C., Duncan, S., Fajardo, C. P., Foy, B. B., Fuhriman, D. A., Gibby, P. D., … Grose, J. H. (2017). Genome Sequences of 19 Novel Erwinia amylovora Bacteriophages. Genome Announcements, 5(46). https://doi.org/10.1128/genomeA.00931-17

Falco, S. C., Zehring, W., & Rothman-Denes, L. B. (1980). DNA-dependent RNA polymerase from bacteriophage N4 virions. Purification and characterization. The Journal of Biological Chemistry, 255(9), 4339–4347.

Fokine, A., Battisti, A. J., Bowman, V. D., Efimov, A. V., Kurochkina, L. P., Chipman, P. R., Mesyanzhinov, V. V., & Rossmann, M. G. (2007). Cryo-EM study of the Pseudomonas bacteriophage phiKZ. Structure, 15(9), 1099–1104.

Goddard, T. D., Huang, C. C., & Ferrin, T. E. (2007). Visualizing density maps with UCSF Chimera. Journal of Structural Biology, 157(1), 281–287.

Goddard, T. D., Huang, C. C., Meng, E. C., Pettersen, E. F., Couch, G. S., Morris, J. H., & Ferrin, T. E. (2018). UCSF ChimeraX: Meeting modern challenges in visualization and analysis. Protein Science: A Publication of the Protein Society, 27(1), 14–25.

Guan, J., Oromí-Bosch, A., Mendoza, S. D., Karambelkar, S., Berry, J., & Bondy-Denomy, J. (2022). RNA targeting with CRISPR-Cas13a facilitates bacteriophage genome engineering. In bioRxiv (p. 2022.02.14.480438). https://doi.org/10.1101/2022.02.14.480438

Haeusser, D. P., & Margolin, W. (2016). Splitsville: structural and functional insights into the dynamic bacterial Z ring. Nature Reviews. Microbiology, 14(5), 305–319.

Iyer, L. M., Anantharaman, V., Krishnan, A., Burroughs, A. M., & Aravind, L. (2021). Jumbo Phages: A Comparative Genomic Overview of Core Functions and Adaptions for Biological Conflicts. Viruses, 13(1). https://doi.org/10.3390/v13010063

Jang, H. B., Fagutao, F. F., Nho, S. W., Park, S. B., Cha, I. S., Yu, J. E., Lee, J. S., Im, S. P., Aoki, T., & Jung, T. S. (2013). Phylogenomic network and comparative genomics reveal a diverged member of the ΦKZ-related group, marine vibrio phage ΦJM-2012. Journal of Virology, 87(23), 12866–12878.

Jin, Y., Sdao, S. M., Dover, J. A., Porcek, N. B., Knobler, C. M., Gelbart, W. M., & Parent, K. N. (2015). Bacteriophage P22 ejects all of its internal proteins before its genome. Virology, 485, 128–134.

Jumper, J., Evans, R., Pritzel, A., Green, T., Figurnov, M., Ronneberger, O., Tunyasuvunakool, K., Bates, R., Žídek, A., Potapenko, A., Bridgland, A., Meyer, C., Kohl, S. A. A., Ballard, A. J., Cowie, A., Romera-Paredes, B., Nikolov, S., Jain, R., Adler, J., … Hassabis, D. (2021). Highly accurate protein structure prediction with AlphaFold. Nature, 596(7873), 583–589.

Kelley, L. A., Mezulis, S., Yates, C. M., Wass, M. N., & Sternberg, M. J. E. (2015). The Phyre2 web portal for protein modeling, prediction and analysis. Nature Protocols, 10(6), 845–858.

Kieser, Q., Noyce, R. S., Shenouda, M., Lin, Y.-C. J., & Evans, D. H. (2020). Cytoplasmic factories, virus assembly, and DNA replication kinetics collectively constrain the formation of poxvirus recombinants. PloS One, 15(1), e0228028.

Knipe, D. M., Prichard, A., Sharma, S., & Pogliano, J. (2022). Replication Compartments of Eukaryotic and Bacterial DNA Viruses: Common Themes Between Different Domains of Host Cells. Annual Review of Virology, 9(1), 307–327.

Konstantinidis, K. T., & Tiedje, J. M. (2005). Genomic insights that advance the species definition for prokaryotes. Proceedings of the National Academy of Sciences of the United States of America, 102(7), 2567–2572.

Kovtun, O., Leneva, N., Bykov, Y. S., Ariotti, N., Teasdale, R. D., Schaffer, M., Engel, B. D., Owen, D. J., Briggs, J. A. G., & Collins, B. M. (2018). Structure of the membrane-assembled retromer coat determined by cryo-electron tomography. In Nature (Vol. 561, Issue 7724, pp. 561–564). https://doi.org/10.1038/s41586-018-0526-z

Kraemer, J. A., Erb, M. L., Waddling, C. A., Montabana, E. A., Zehr, E. A., Wang, H., Nguyen, K., Pham, D. S. L., Agard, D. A., & Pogliano, J. (2012). A phage tubulin assembles dynamic filaments by an atypical mechanism to center viral DNA within the host cell. Cell, 149(7), 1488–1499.

Kremer, J. R., Mastronarde, D. N., & McIntosh, J. R. (1996). Computer visualization of three-dimensional image data using IMOD. Journal of Structural Biology, 116(1), 71–76.

Krylov, V., Bourkaltseva, M., Pleteneva, E., Shaburova, O., Krylov, S., Karaulov, A., Zhavoronok, S., Svitich, O., & Zverev, V. (2021). Phage phiKZ-The First of Giants. Viruses, 13(2). https://doi.org/10.3390/v13020149

Labarde, A., Jakutyte, L., Billaudeau, C., Fauler, B., López-Sanz, M., Ponien, P., Jacquet, E., Mielke, T., Ayora, S., Carballido-López, R., & Tavares, P. (2021). Temporal compartmentalization of viral infection in bacterial cells. Proceedings of the National Academy of Sciences of the United States of America, 118(28). https://doi.org/10.1073/pnas.2018297118

Lam, V., & Villa, E. (2021). Practical Approaches for Cryo-FIB Milling and Applications for Cellular Cryo-Electron Tomography. Methods in Molecular Biology, 2215, 49–82.

Larsen, R. A., Cusumano, C., Fujioka, A., Lim-Fong, G., Patterson, P., & Pogliano, J. (2007). Treadmilling of a prokaryotic tubulin-like protein, TubZ, required for plasmid stability in Bacillus thuringiensis. Genes & Development, 21(11), 1340–1352.

Laughlin, T. G., Deep, A., Prichard, A. M., Seitz, C., Gu, Y., Enustun, E., Suslov, S., Khanna, K., Birkholz, E. A., Armbruster, E., McCammon, J. A., Amaro, R. E., Pogliano, J., Corbett, K. D., & Villa, E. (2022). Architecture and self-assembly of the jumbo bacteriophage nuclear shell. Nature, 608(7922), 429–435.

Letunic, I., & Bork, P. (2021). Interactive Tree Of Life (iTOL) v5: an online tool for phylogenetic tree display and annotation. Nucleic Acids Research, 49(W1), W293–W296.

Liu, Y.-T., Zhang, H., Wang, H., Tao, C.-L., Bi, G.-Q., & Hong Zhou, Z. (n.d.). Isotropic Reconstruction of Electron Tomograms with Deep Learning. https://doi.org/10.1101/2021.07.17.452128

Madeira, F., Pearce, M., Tivey, A. R. N., Basutkar, P., Lee, J., Edbali, O., Madhusoodanan, N., Kolesnikov, A., & Lopez, R. (2022). Search and sequence analysis tools services from EMBL-EBI in 2022. Nucleic Acids Research. https://doi.org/10.1093/nar/gkac240

Malone, L. M., Warring, S. L., Jackson, S. A., Warnecke, C., Gardner, P. P., Gumy, L. F., & Fineran, P. C. (2020). A jumbo phage that *forms a nucleus-like structure evades CRISPR-Cas DNA targeting but is vulnerable to type III RNA-based immunity*. Nature Microbiology, 5(1), 48–55.

Martinez-Sanchez, A., Garcia, I., Asano, S., Lucic, V., & Fernandez, J.-J. (2014). Robust membrane detection based on tensor voting for electron tomography. In Journal of Structural Biology (Vol. 186, Issue 1, pp. 49–61). https://doi.org/10.1016/j.jsb.2014.02.015

Mastronarde, D. N. (2005). Automated electron microscope tomography using robust prediction of specimen movements. Journal of Structural Biology, 152(1), 36–51.

Mathee, K., Narasimhan, G., Valdes, C., Qiu, X., Matewish, J. M., Koehrsen, M., Rokas, A., Yandava, C. N., Engels, R., Zeng, E., Olavarietta, R., Doud, M., Smith, R. S., Montgomery, P., White, J. R., Godfrey, P. A., Kodira, C., Birren, B., Galagan, J. E., & Lory, S. (2008). Dynamics of Pseudomonas aeruginosa genome evolution. Proceedings of the National Academy of Sciences of the United States of America, 105(8), 3100–3105.

Matsui, T., Yoshikawa, G., Mihara, T., Chatchawankanphanich, O., Kawasaki, T., Nakano, M., Fujie, M., Ogata, H., & Yamada, T. (2017). Replications of Two Closely Related Groups of Jumbo Phages Show Different Level of Dependence on Host-encoded RNA Polymerase. Frontiers in Microbiology, 8, 1010.

Meier-Kolthoff, J. P., & Göker, M. (2017). VICTOR: genome-based phylogeny and classification of prokaryotic viruses. Bioinformatics, 33(21), 3396–3404.

Mendoza, S. D., Nieweglowska, E. S., Govindarajan, S., Leon, L. M., Berry, J. D., Tiwari, A., Chaikeeratisak, V., Pogliano, J., Agard, D. A., & Bondy-Denomy, J. (2020). A bacteriophage nucleus-like compartment shields DNA from CRISPR nucleases. Nature, 577(7789), 244–248.

Molineux, I. J., & Panja, D. (2013). Popping the cork: mechanisms of phage genome ejection. Nature Reviews. Microbiology, 11(3), 194–204.

Montabana, E. A., & Agard, D. A. (2014). Bacterial tubulin TubZ-Bt transitions between a two-stranded intermediate and a *four-stranded filament upon GTP hydrolysis*. Proceedings of the National Academy of Sciences of the United States of America, 111(9), 3407–3412.

Navarro, P., Scaramuzza, S., Stahlberg, H., & Castaño-Díez, D. (2020). The Dynamo Software Package for Cryo-electron Tomography and Subtomogram Averaging. In Microscopy and Microanalysis (Vol. 26, Issue S2, pp. 3142–3145). https://doi.org/10.1017/s1431927620023958

Nguyen, K. T., Sugie, J., Khanna, K., Egan, M. E., Birkholz, E. A., Lee, J., Beierschmitt, C., Villa, E., & Pogliano, J. (2021). Selective transport of fluorescent proteins into the phage nucleus. PloS One, 16(6), e0251429.

Nishimura, Y., Yoshida, T., Kuronishi, M., Uehara, H., Ogata, H., & Goto, S. (2017). ViPTree: the viral proteomic tree server. Bioinformatics, 33(15), 2379–2380.

Orekhova, M., Koreshova, A., Artamonova, T., Khodorkovskii, M., & Yakunina, M. (2019). The study of the phiKZ phage non-canonical non-virion RNA polymerase. Biochemical and Biophysical Research Communications, 511(4), 759–764.

Park, S.-C., Lee, K., Kim, Y. O., Won, S., & Chun, J. (2019). Large-Scale Genomics Reveals the Genetic Characteristics of Seven Species and Importance of Phylogenetic Distance for Estimating Pan-Genome Size. Frontiers in Microbiology, 10, 834.

Paszkowski, P., Noyce, R. S., & Evans, D. H. (2016). Live-Cell Imaging of Vaccinia Virus Recombination. PLoS Pathogens, 12(8), e1005824.

Pyle, E., Hutchings, J., & Zanetti, G. (2022). Strategies for Picking Membrane-Associated Particles within Subtomogram Averaging Workflows. In Faraday Discussions. https://doi.org/10.1039/d2fd00022a

Robert, X., & Gouet, P. (2014). Deciphering key features in protein structures with the new ENDscript server. Nucleic Acids Research, 42(Web Server issue), W320–W324.

Scheres, S. H. W. (2012). RELION: implementation of a Bayesian approach to cryo-EM structure determination. Journal of Structural Biology, 180(3), 519–530.

Schlieper, D., Oliva, M. A., Andreu, J. M., & Löwe, J. (2005). Structure of bacterial tubulin BtubA/B: evidence for horizontal gene transfer. Proceedings of the National Academy of Sciences of the United States of America, 102(26), 9170–9175.

Schrödinger, LLC. (2021). The PyMOL Molecular Graphics System, Version 2.5.2.

Sharma, R., Pielstick, B. A., Bell, K. A., Nieman, T. B., Stubbs, O. A., Yeates, E. L., Baltrus, D. A., & Grose, J. H. (2019). A Novel, Highly Related Jumbo Family of Bacteriophages That Were Isolated Against Erwinia. Frontiers in Microbiology, 10, 1533.

Sokolova, M. L., Misovetc, I., & Severinov, K., V. (2020). Multisubunit RNA Polymerases of Jumbo Bacteriophages. Viruses, 12(10). https://doi.org/10.3390/v12101064

Tegunov, D., & Cramer, P. (2019). Real-time cryo-electron microscopy data preprocessing with Warp. Nature Methods, 16(11), 1146–1152.

Tegunov, D., Xue, L., Dienemann, C., Cramer, P., & Mahamid, J. (2021). Multi-particle cryo-EM refinement with M visualizes ribosome-antibiotic complex at 3.5 Å in cells. Nature Methods, 18(2), 186–193.

Thomas, J. A., Rolando, M. R., Carroll, C. A., Shen, P. S., Belnap, D. M., Weintraub, S. T., Serwer, P., & Hardies, S. C. (2008). Characterization of Pseudomonas chlororaphis myovirus 201varphi2-1 via genomic sequencing, mass spectrometry, and electron microscopy. Virology, 376(2), 330–338.

Thomas, J. A., Weintraub, S. T., Wu, W., Winkler, D. C., Cheng, N., Steven, A. C., & Black, L. W. (2012). Extensive proteolysis of head *and inner body proteins by a morphogenetic protease in the giant Pseudomonas aeruginosa phage ϕKZ*. Molecular Microbiology, 84(2), 324–339.

Tomer, E., Cohen, E. M., Drayman, N., Afriat, A., Weitzman, M. D., Zaritsky, A., & Kobiler, O. (2019). Coalescing replication *compartments provide the opportunity for recombination between coinfecting herpesviruses*. FASEB Journal: Official Publication of the Federation of American Societies for Experimental Biology, 33(8), 9388–9403.

Trinh, J. T., Shao, Q., Guan, J., & Zeng, L. (2020). Emerging heterogeneous compartments by viruses in single bacterial cells. Nature Communications, 11(1), 3813.

Vanneste, J. L. (2000). Fire Blight: The Disease and Its Causative Agent, Erwinia Amylovora. CABI.

van Tonder, A. J., Mistry, S., Bray, J. E., Hill, D. M. C., Cody, A. J., Farmer, C. L., Klugman, K. P., von Gottberg, A., Bentley, S. D., Parkhill, J., Jolley, K. A., Maiden, M. C. J., & Brueggemann, A. B. (2014). Defining the estimated core genome of bacterial populations using a Bayesian decision model. PLoS Computational Biology, 10(8), e1003788.

Yakunina, M., Artamonova, T., Borukhov, S., Makarova, K. S., Severinov, K., & Minakhin, L. (2015). A non-canonical multisubunit RNA polymerase encoded by a giant bacteriophage. Nucleic Acids Research, 43(21), 10411–10420.

Yee, B., Lafi, F. F., Oakley, B., Staley, J. T., & Fuerst, J. A. (2007). A canonical FtsZ protein in Verrucomicrobium spinosum, a member of *the Bacterial phylum Verrucomicrobia that also includes tubulin-producing Prosthecobacter species*. BMC Evolutionary Biology, 7, 37.

Zehr, E. A., Kraemer, J. A., Erb, M. L., Coker, J. K. C., Montabana, E. A., Pogliano, J., & Agard, D. A. (2014). The structure and assembly mechanism of a novel three-stranded tubulin filament that centers phage DNA. Structure, 22(4), 539–548.

