## Supplementary Figures for "Identifying the core genome of the nucleus-forming bacteriophage family and characterization of *Erwinia* phage RAY"

### SUPPLEMENTARY INFORMATION

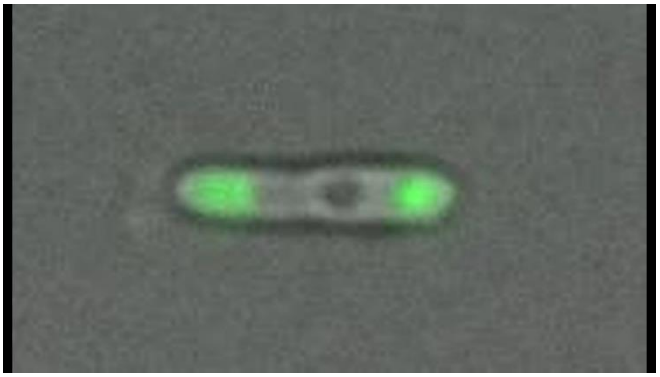

**Movie 1. H-NS Timelapse.** Images were taken every 10 minutes, from 20 mpi to 100 mpi. The phage nucleus can be seen forming in brightfield (grayscale) and the H-NS-GFP (green) can be seen migrating toward the cell poles as the infection progresses.

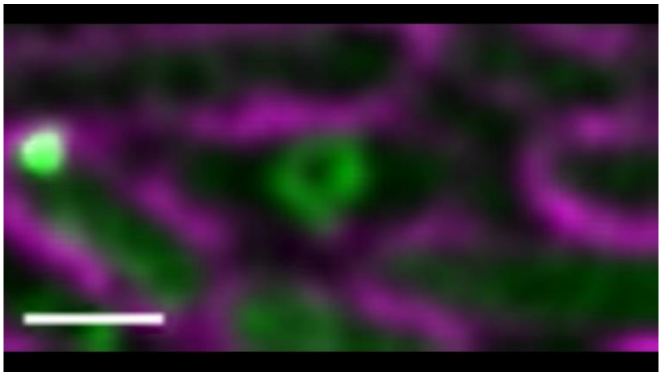

**Movie 2. Nucleus rotation.** Images were taken every 4 seconds for 2 minutes at 60 mpi. The duration of clearest rotation for this representative cell lasted for 32 seconds, which are shown in this time lapse. Scale bar is 1 micron; magenta is membrane stain FM4-64, cyan is DNA stain DAPI, and green is GFP.

**Table 1. List of chimallin-encoding phage used in our analyses.** These phage were used for our determination of the Chimalliviridae core genome.

| Host | Phage Name | Host | Phage Name |
|---|---|---|---|
| Aeromonas | PS1 | Erwinia | pEa_SNUABM_37 |
| Aeromonas | pAEv1810 | Erwinia | AH06 |
| Aeromonas | CF8 | Erwinia | vB_EamM_RAY |
| Aeromonas | LAh10 | Erwinia | vB_EamM_Joad |
| Bacillus | vB_BspM_AgentSmith | Erwinia | AH04 |
| Burkholderia | FLC6 | Escherichia | vB_EcoM_Goslar |
| Burkholderia | FLC9 | Halocynthia | JM-2012 |
| Cronobacter | CR5 | Klebsiella | vB_KvM-Eowyn |
| Edwardsiella | pEt-SU | Klebsiella | vB_KpM_FBKp24 |
| Erwinia | pEa_SNUABM_29 | Klebsiella | KpLz-2_45 |
| Erwinia | pEa_SNUABM_11 | Klebsiella | Miami |
| Erwinia | vB_EamM_Asesino | Klebsiella | N1M2 |
| Erwinia | vB_EamM_Huxley | Kosakonia | Kc263 |
| Erwinia | pEa_SNUABM_8 | Photobacterium | PDCC-1 |
| Erwinia | vB_EamM_ChrisDB | Proteus | 10 |
| Erwinia | vB_EamM_Caitlin | Pseudomonas | pPa_SNUABM_DT01 |
| Erwinia | phiEaH2 | Pseudomonas | 201phi2-1 |
| Erwinia | Wellington | Pseudomonas | PhiPA3 |
| Erwinia | Derbicus | Pseudomonas | phiKZ |

| | | | |
|---|---|---|---|
| Erwinia | vB_EamM_Phobos | Pseudomonas | PA1C |
| Erwinia | pEa_SNUABM_54 | Pseudomonas | Psa21 |
| Erwinia | PhiEaH1 | Pseudomonas | Phabio |
| Pseudomonas | Noxifer | Serratia | Moabite |
| Pseudomonas | EL | Vibrio | vB_VmeM-Yong MS32 |
| Pseudomonas | OBP | Vibrio | BONAISHI |
| Ralstonia | RSL2 | Vibrio | vB_pir03 |
| Ralstonia | RSF1 | Vibrio | vB_VpaM_sm033 |
| Ralstonia | RP31 | Vibrio | pVa-21 |
| Salmonella | STsAS | Vibrio | Aphrodite1 |
| Salmonella | SPAsTU | Vibrio | VP4B |
| Salmonella | SPN3US | Vibrio | pTD1 |
| Salmonella | vB_SalM_SA002 | Xanthomonas | Xoo-sp14 |
| Serratia | PCH45 | Xanthomonas | vB_XciM_LucasX |

**Table 2. VICTOR predictions of taxonomic groups.** Predicted family, subfamily, genus, and species categories from VICTOR were used to guide our inclusion of phages in our analyses.

| Genomes | species | genus | subfamily | family |
|---|---|---|---|---|
| Mycoplasma phage P1 (NC_002515) | 41 | 11 | 3 | 1 |
| Clostridium phage phiCP7R (NC_017980) | 46 | 14 | 4 | 2 |
| Clostridium phage phiCPV4 (NC_018083) | 46 | 14 | 4 | 2 |
| Escherichia phage T4 (NC_000866) | 40 | 10 | 2 | 3 |
| Bacillus phage G (NC_023719) | 51 | 16 | 6 | 3 |
| Erwinia phage vB_EamM_Alexandra (NC_047995) | 69 | 23 | 10 | 3 |
| Cronobacter phage vB_CsaP_Ss1 (KM058087) | 5 | 1 | 1 | 4 |
| Pectobacterium phage ZF40 (NC_019522) | 47 | 15 | 5 | 4 |
| Erwinia phage phiEa21-4 (NC_011811) | 63 | 19 | 7 | 4 |
| Erwinia phage phiEa104 (NC_015292) | 63 | 19 | 7 | 4 |
| Erwinia phage vB_Eam-MM7 (NC_041978) | 63 | 19 | 7 | 4 |
| Yersinia phage vB_YenP_ISAO8 (NC_028850) | 53 | 22 | 9 | 4 |
| Yersinia phage phiR8-01 (NC_047951) | 68 | 22 | 9 | 4 |
| Salmonella phage ZCSE2 (NC_048179) | 74 | 26 | 11 | 4 |
| Salmonella phage SE4 (NC_048764) | 74 | 26 | 11 | 4 |
| Salmonella phage BP63 (NC_031250) | 77 | 26 | 11 | 4 |
| Salmonella phage UPF_BP2 (NC_048649) | 77 | 26 | 11 | 4 |
| Salmonella phage SE13 (NC_048763) | 77 | 26 | 11 | 4 |
| Salmonella phage yarpen (NC_048863) | 77 | 26 | 11 | 4 |
| Salmonella phage birk (NC_048864) | 77 | 26 | 11 | 4 |
| Staphylococcus phage PALS_2 (MN091626) | 21 | 5 | 8 | 5 |
| Yersinia phage phiR1-37 (NC_016163) | 44 | 12 | 8 | 5 |

| | | | | |
|---|---|---|---|---|
| Bacillus phage PBS1 (NC_043027) | 67 | 21 | 8 | 5 |
| Burkholderia phage FLC9 (LC667451) | 8 | 2 | 12 | 5 |
| Klebsiella phage vB_KvM-Eowyn (LR881104) | 9 | 3 | 12 | 5 |
| Vibrio phage vB_VmeM-Yong XC31 (MK308674) | 15 | 4 | 12 | 5 |
| Pseudomonas phage pPa_SNUABM_DT01 (MW735835) | 30 | 6 | 12 | 5 |
| Bacillus phage vB_BspM_AgentSmith (MW749006) | 31 | 7 | 12 | 5 |
| Xanthomonas phage Xoo-sp14 (MT939492) | 28 | 8 | 12 | 5 |
| Xanthomonas phage vB_XciM_LucasX (MW825358) | 33 | 8 | 12 | 5 |
| Erwinia phage pEa_SNUABM_54 (MW879341) | 36 | 9 | 12 | 5 |
| Vibrio phage JM-2012 (NC_017975) | 45 | 13 | 12 | 5 |
| Ralstonia phage RP31 (AP017925) | 1 | 17 | 12 | 5 |
| Burkholderia phage FLC6 (LC592711) | 7 | 17 | 12 | 5 |
| Ralstonia phage RSF1 (NC_028899) | 54 | 17 | 12 | 5 |
| Ralstonia phage RSL2 (NC_028950) | 55 | 17 | 12 | 5 |
| Salmonella phage vB_SalM_SA002 (MN445183) | 23 | 18 | 12 | 5 |
| Proteus phage 10 (MT661596) | 25 | 18 | 12 | 5 |
| Erwinia phage pEa_SNUABM_37 (MW845760) | 35 | 18 | 12 | 5 |
| Erwinia phage AH06 (MZ501268) | 39 | 18 | 12 | 5 |
| Erwinia phage vB_EamM_RAY (NC_041973) | 62 | 18 | 12 | 5 |
| Pseudomonas phage 201phi2-1 (EU197055) | 2 | 20 | 12 | 5 |
| Pseudomonas phage PhiPA3 (HQ630627) | 3 | 20 | 12 | 5 |
| Pseudomonas phage Phabio (MF042360) | 10 | 20 | 12 | 5 |
| Pseudomonas phage Psa21 (MK552327) | 16 | 20 | 12 | 5 |
| Pseudomonas phage PA1C (MK599315) | 17 | 20 | 12 | 5 |

| | | | | |
|---|---|---|---|---|
| Pseudomonas phage phiKZ (NC_004629) | 42 | 20 | 12 | 5 |
| Pseudomonas phage Noxifer (NC_041994) | 64 | 20 | 12 | 5 |
| Escherichia phage vB_EcoM_Goslar (NC_048170) | 71 | 24 | 12 | 5 |
| Vibrio phage pVa-21 (KY499642) | 6 | 25 | 12 | 5 |
| Salmonella phage STsAS (MH221128) | 12 | 25 | 12 | 5 |
| Salmonella phage SPAsTU (MH221129) | 13 | 25 | 12 | 5 |
| Erwinia phage pEa_SNUABM_29 (MW812339) | 32 | 25 | 12 | 5 |
| Erwinia phage pEa_SNUABM_11 (MW845758) | 34 | 25 | 12 | 5 |
| Erwinia phage phiEaH2 (NC_019929) | 48 | 25 | 12 | 5 |
| Cronobacter phage CR5 (NC_021531) | 49 | 25 | 12 | 5 |
| Salmonella phage SPN3US (NC_027402) | 52 | 25 | 12 | 5 |
| Erwinia phage vB_EamM_Phobos (NC_031043) | 56 | 25 | 12 | 5 |
| Erwinia phage vB_EamM_Asesino (NC_031107) | 57 | 25 | 12 | 5 |
| Erwinia phage vB_EamM_Caitlin (NC_031120) | 58 | 25 | 12 | 5 |
| Erwinia phage pEa_SNUABM_8 (MW760841) | 59 | 25 | 12 | 5 |
| Erwinia phage vB_EamM_ChrisDB (NC_031126) | 59 | 25 | 12 | 5 |
| Erwinia phage vB_EamM_Huxley (NC_031127) | 60 | 25 | 12 | 5 |
| Erwinia phage Wellington (NC_048016) | 70 | 25 | 12 | 5 |
| Erwinia phage Derbicus (NC_048173) | 72 | 25 | 12 | 5 |
| Serratia phage PCH45 (MN334766) | 22 | 27 | 12 | 5 |
| Klebsiella phage vB_KpM_FBKp24 (MW394391) | 29 | 27 | 12 | 5 |
| Erwinia phage PhiEaH1 (NC_023610) | 50 | 27 | 12 | 5 |
| Serratia phage Moabite (NC_048792) | 75 | 27 | 12 | 5 |
| Klebsiella phage KpLz-2_45 (NC_061418) | 78 | 27 | 12 | 5 |

| | | | | |
|---|---|---|---|---|
| Pseudomonas phage OBP (JN627160) | 4 | 28 | 12 | 5 |
| Erwinia phage vB_EamM_Joad (MF459647) | 11 | 28 | 12 | 5 |
| Aeromonas phage CF8 (MK774614) | 18 | 28 | 12 | 5 |
| Aeromonas phage LAh10 (MK838116) | 19 | 28 | 12 | 5 |
| Aeromonas phage PS1 (MN032614) | 20 | 28 | 12 | 5 |
| Klebsiella phage N1M2 (MN642089) | 24 | 28 | 12 | 5 |
| Klebsiella phage Miami (MT701590) | 26 | 28 | 12 | 5 |
| Kosakonia phage Kc263 (MZ348422) | 37 | 28 | 12 | 5 |
| Erwinia phage AH04 (MZ501267) | 38 | 28 | 12 | 5 |
| Pseudomonas phage EL (NC_007623) | 43 | 28 | 12 | 5 |
| Vibrio phage pTD1 (NC_041916) | 61 | 28 | 12 | 5 |
| Vibrio phage Aphrodite1 (NC_042100) | 65 | 28 | 12 | 5 |
| Vibrio phage VP4B (NC_042136) | 66 | 28 | 12 | 5 |
| Edwardsiella phage pEtSU (NC_048182) | 73 | 28 | 12 | 5 |
| Photobacterium phage PDCC-1 (NC_048821) | 76 | 28 | 12 | 5 |
| Aeromonas phage pAEv1810 (OL964756) | 79 | 28 | 12 | 5 |
| Vibrio phage BONAISHI (MH595538) | 14 | 29 | 12 | 5 |
| Vibrio phage vB_pir03 (MT811961) | 27 | 29 | 12 | 5 |
| Vibrio phage vB_VpaM_sm033 (OV032902) | 80 | 29 | 12 | 5 |

**Table 3. Core genome blocks by function.** The core genome numbers were determined with homologies within and across the genomes of ΦKZ, Goslar, and RAY. Each block is color coded and marked by number, and the last column indicates putative functions from PSI-BLAST hits, many of which are hypothetical. If a core gene has multiple homologs in one phage (for instance, cg42), they will be lettered by numerical order (for instance, RAY gp017 = cg42A, RAY gp018 = cg42B, and RAY gp019 = cg42C).

| Block Number | Core Genome Number | Goslar | ΦKZ | RAY | Putative Function |
|---|---|---|---|---|---|
| 1 | cg1 | gp192 | gp049 | gp219 | hypothetical protein |
| | cg2 | gp191 | gp050 | gp220 | virion DNAP |
| | cg3 | gp190 | gp052 | gp221 | hypothetical protein |
| | cg4 | gp189 | gp054 | gp222 | Nuclear Shell Protein |
| | cg5 | gp188 | gp055 | gp070, gp223 | putative DNA-directed RNA polymerase beta subunit |
| | cg6 | gp184 | gp059 | gp229 | hypothetical protein |
| 2 | cg7 | gp180 | gp062 | gp236 | hypothetical protein |
| | cg8 | gp178 | gp065 | gp238 | putative nuclease |
| | cg9 | gp177 | gp066 | gp239 | hypothetical protein |
| | cg10 | gp176 | gp067 | gp240 | hypothetical protein |
| | cg11 | gp175 | gp068 | gp243 | putative nvRNAP (non-virion RNAP) sigma factor |
| | cg12 | gp174 | gp069 | gp244 | hypothetical protein |
| | cg13 | gp173 | gp070 | gp245 | hypothetical protein |
| | cg14 | gp172 | n/a | gp246 | hypothetical protein |
| | cg15 | gp171 | gp071, gp073 | gp248 | putative DNA directed RNA polymerase beta subunit |
| | cg16 | gp165 | gp074 | gp249 | putative DNA directed RNA polymerase beta subunit |
| | cg17 | gp164 | gp075 | gp250 | putative RAD2/SF2 helicase |
| 3 | cg18 | gp081 | gp077 | gp267 | hypothetical protein |
| | cg19 | gp079 | gp079 | gp269 | hypothetical protein |
| | cg20 | gp078 | gp080 | gp270 | putative DNA-directed RNA polymerase beta prime subunit |
| | cg21 | gp068 | gp082 | gp285 | putative DNA polymerase |
| | cg22 | gp067 | gp084 | gp286 | putative virion structural protein |
| 4 | cg23 | gp063 | gp087 | gp290 | putative virion structural protein |
| | cg24 | gp062 | gp088 | gp291 | putative virion structural protein |
| | cg25 | gp061 | gp089 | gp292 | hypothetical protein |
| | cg26 | gp060 | gp090 | gp293 | putative virion structural protein |
| | cg27 | gp058 | gp093, gp162, gp163 | gp295, gp298 | virion structural protein/ internal head |

| | | | | | |
|---|---|---|---|---|---|
| 5 | cg28 | gp051 | gp098 | gp304 | hypothetical protein |
| | cg29 | gp050 | gp099 | gp305 | putative virion structural protein |
| | cg30 | gp049 | gp100 | gp306 | hypothetical protein |
| | cg31 | gp048 | gp101 | gp307 | putative virion structural protein |
| | cg32 | gp008 | gp188 | gp311 | putative thymidylate kinase |
| | cg33 | gp043 | gp118 | gp315 | putative DnaB helicase |
| | cg34 | gp041 | gp120 | gp317 | major capsid protein |
| | cg35 | gp040 | gp122 | gp001 | hypothetical |
| | cg36 | gp039 | gp123 | gp002 | putative RNA polymerase beta subunit |
| | cg37 | gp036 | gp129 | gp006 | putative virion structural protein |
| | cg38 | gp035 | gp128 | gp007 | putative virion structural protein |
| | cg39 | gp032 | gp139 | gp010 | virion structural protein |
| | cg40 | gp010 | gp140 | gp012 | hypothetical |
| | cg41 | gp030 | gp130 | gp016 | virion structural protein |
| | cg42 | gp025, gp028, gp029 | gp131, gp132, gp134, gp135 | gp017, gp018, gp019 | virion structural protein/ tail |
| | cg43 | gp013 | gp164 | gp021 | structural protein |
| | cg44 | gp012 | gp165 | gp023 | putative SbcC-like protein |
| 6 | cg45 | gp249 | gp161 | gp144 | hypothetical |
| | cg46 | gp248 | gp157 | gp145 | virion structural protein |
| | cg47 | gp246 | gp155 | gp147 | putative ribonuclease HI |
| | cg48 | gp244 | gp153 | gp149 | hypothetical |
| | cg49 | gp243 | gp152 | gp150 | putative UvsX protein |
| | cg50 | gp240 | gp149 | gp154 | virion structural protein |
| | cg51 | gp238 | gp147 | gp156 | hypothetical |
| | cg52 | gp235 | gp171 | gp159 | hypothetical |
| | cg53 | gp234 | gp174 | gp160 | hypothetical |
| | cg54 | gp233 | gp182 | gp161 | virion structural protein |
| | cg55 | gp232 | gp181 | gp162 | putative lysozyme domain protein |
| | cg56 | gp231 | gp180 | gp163 | putative DNA-direct RNA polymerase beta subunit 2 |
| | cg57 | gp228 | gp178 | gp164 | putative RNA polymerase beta subunit |
| | cg58 | gp226 | gp177 | gp167 | hypothetical protein |
| | cg59 | gp225 | gp176 | gp168 | hypothetical protein |

| | | | | | |
|---|---|---|---|---|---|
| | cg60 | gp223 | gp175 | gp170 | putative virion structural protein |
| 7 | cg61 | gp218 | gp030 | gp178 | putative major virion structural protein |
| | cg62 | gp217 | gp029 | gp179 | putative tail sheath protein |
| | cg63 | gp216 | gp028 | gp180 | putative virion structural protein |
| | cg64 | gp215 | gp027 | gp181 | putative virion structural protein |
| | cg65 | gp214 | gp026 | gp182 | putative structural protein |
| | cg66 | gp213 | gp025 | gp183 | putative terminase large subunit function |
| | cg67 | gp211 | gp032 | gp187 | hypothetical protein |
| | cg68 | gp202 | gp042 | gp209 | hypothetical protein |

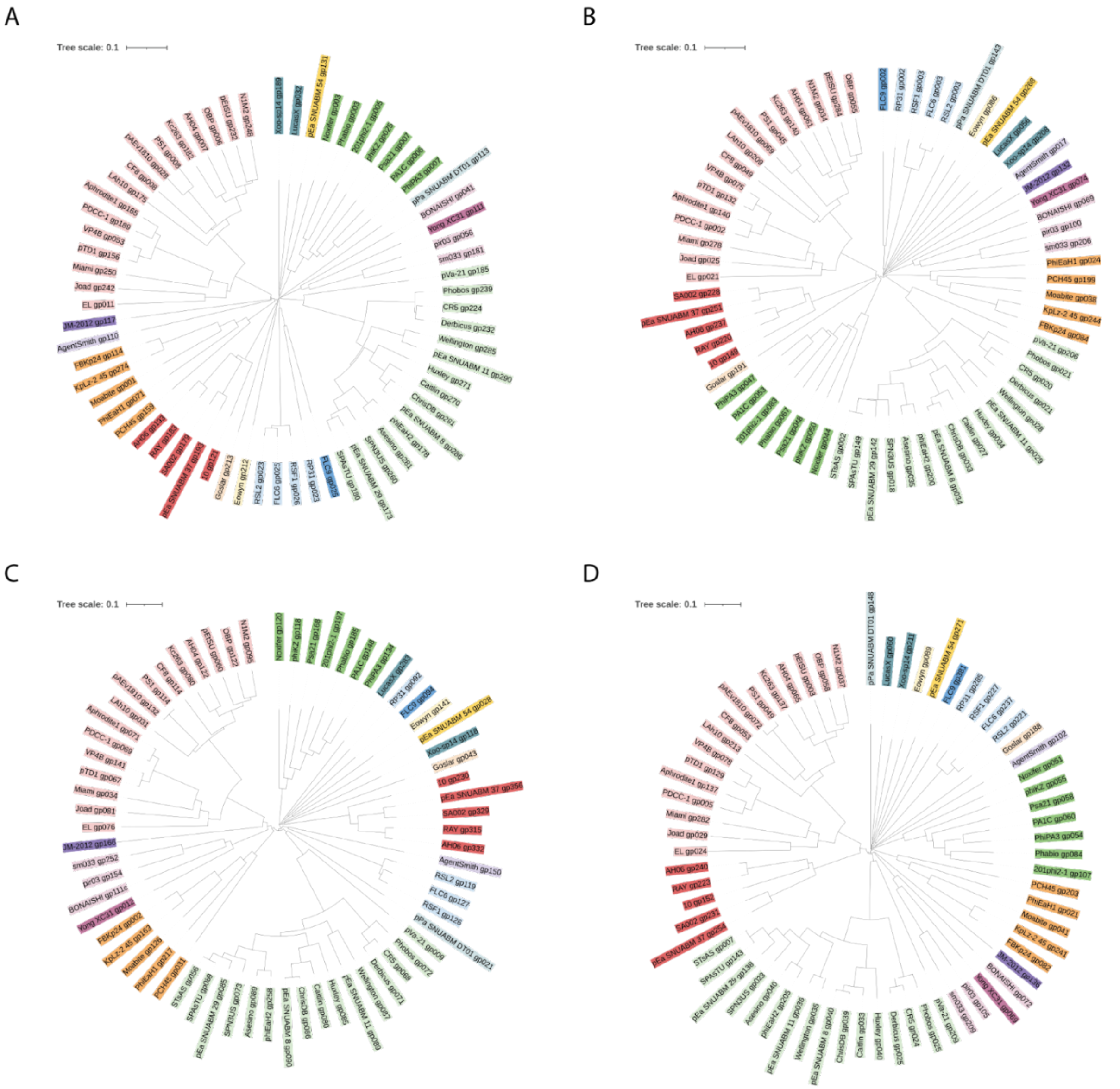


**Figure S1. Phylogenetic trees including Chimalliviridae members based on well-conserved phage proteins.** (A) A protein tree of terminase large subunit homologs from Chimalliviridae. (B) A protein tree of DNA polymerase homologs from Chimalliviridae. (C) A protein tree of replicative (DnaB-like) helicases from Chimalliviridae. (D) A protein tree of non-virion RNA polymerase β' subunit 1 homologs from Chimalliviridae. All homologs were found by the top PSI-BLAST hit. Chimalliviridae are color-coded by predicted genus as in Figure 1.

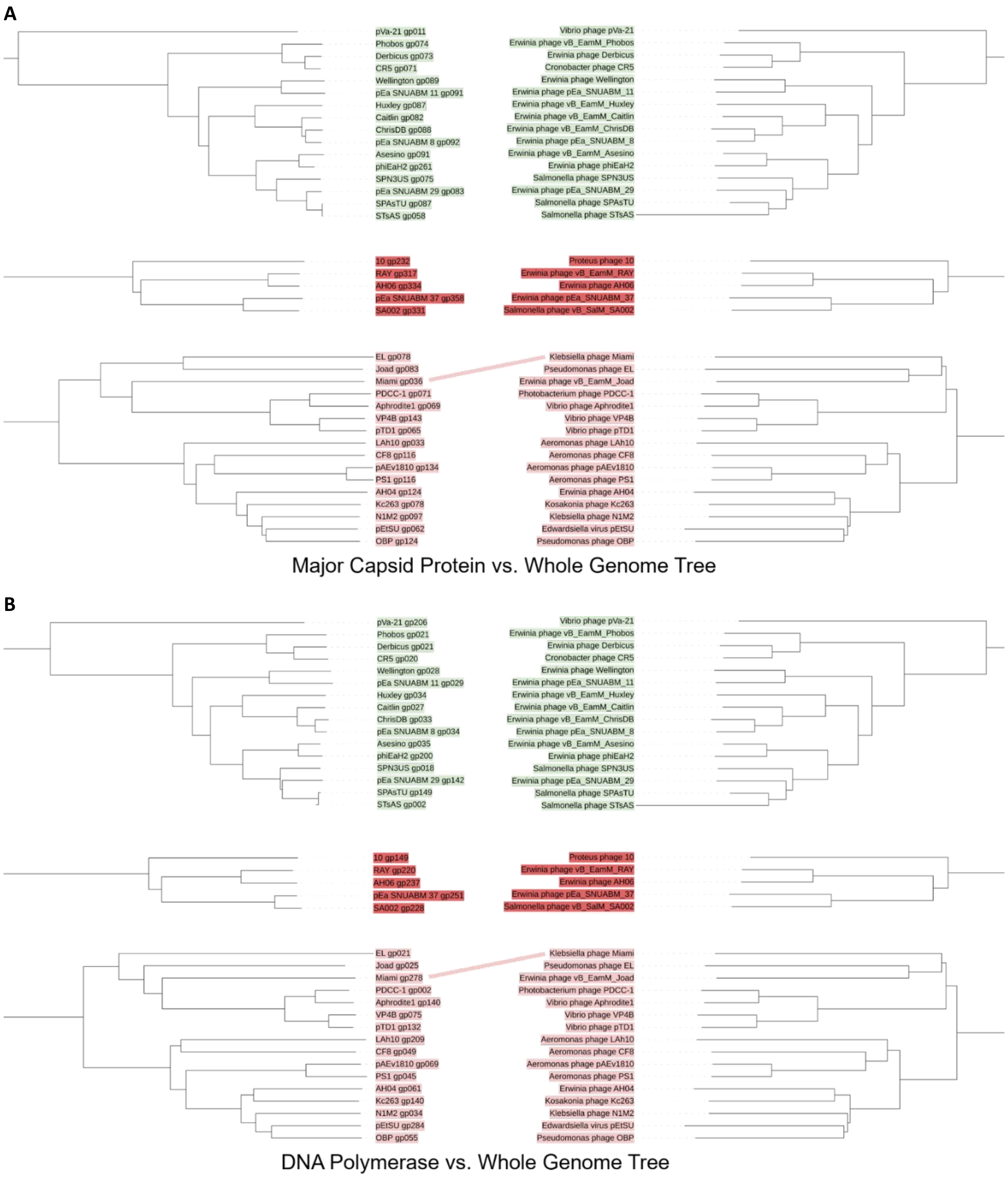
A
pVa-21 gp011
Phobos gp074
Derbicus gp073
CR5 gp071
Wellington gp089
pEa SNUABM 11 gp091
Huxley gp087
Caitlin gp082
ChrisDB gp088
pEa SNUABM 8 gp092
Asesino gp091
phiEaH2 gp261
SPN3US gp075
pEa SNUABM 29 gp083
SPAsTU gp087
STsAS gp058
Vibrio phage pVa-21
Erwinia phage vB_EamM_Phobos
Erwinia phage Derbicus
Cronobacter phage CR5
Erwinia phage Wellington
Erwinia phage pEa_SNUABM_11
Erwinia phage vB_EamM_Huxley
Erwinia phage vB_EamM_Caitlin
Erwinia phage vB_EamM_ChrisDB
Erwinia phage pEa_SNUABM_8
Erwinia phage vB_EamM_Asesino
Erwinia phage phiEaH2
Salmonella phage SPN3US
Erwinia phage pEa_SNUABM_29
Salmonella phage SPAsTU
Salmonella phage STsAS
10 gp232
RAY gp317
AH06 gp334
pEa SNUABM 37 gp358
SA002 gp331
Proteus phage 10
Erwinia phage vB_EamM_RAY
Erwinia phage AH06
Erwinia phage pEa_SNUABM_37
Salmonella phage vB_SalM_SA002
EL gp078
Joad gp083
Miami gp036
PDCC-1 gp071
Aphrodite1 gp069
VP4B gp143
pTD1 gp065
LAh10 gp033
CF8 gp116
pAEv1810 gp134
PS1 gp116
AH04 gp124
Kc263 gp078
N1M2 gp097
pEtSU gp062
OBP gp124
Klebsiella phage Miami
Pseudomonas phage EL
Erwinia phage vB_EamM_Joad
Photobacterium phage PDCC-1
Vibrio phage Aphrodite1
Vibrio phage VP4B
Vibrio phage pTD1
Aeromonas phage LAh10
Aeromonas phage CF8
Aeromonas phage pAEv1810
Aeromonas phage PS1
Erwinia phage AH04
Kosakonia phage Kc263
Klebsiella phage N1M2
Edwardsiella virus pEtSU
Pseudomonas phage OBP
Major Capsid Protein vs. Whole Genome Tree
B
pVa-21 gp206
Phobos gp021
Derbicus gp021
CR5 gp020
Wellington gp028
pEa SNUABM 11 gp029
Huxley gp034
Caitlin gp027
ChrisDB gp033
pEa SNUABM 8 gp034
Asesino gp035
phiEaH2 gp200
SPN3US gp018
pEa SNUABM 29 gp142
SPAsTU gp149
STsAS gp002
Vibrio phage pVa-21
Erwinia phage vB_EamM_Phobos
Erwinia phage Derbicus
Cronobacter phage CR5
Erwinia phage Wellington
Erwinia phage pEa_SNUABM_11
Erwinia phage vB_EamM_Huxley
Erwinia phage vB_EamM_Caitlin
Erwinia phage vB_EamM_ChrisDB
Erwinia phage pEa_SNUABM_8
Erwinia phage vB_EamM_Asesino
Erwinia phage phiEaH2
Salmonella phage SPN3US
Erwinia phage pEa_SNUABM_29
Salmonella phage SPAsTU
Salmonella phage STsAS
10 gp149
RAY gp220
AH06 gp237
pEa SNUABM 37 gp251
SA002 gp228
Proteus phage 10
Erwinia phage vB_EamM_RAY
Erwinia phage AH06
Erwinia phage pEa_SNUABM_37
Salmonella phage vB_SalM_SA002
EL gp021
Joad gp025
Miami gp278
PDCC-1 gp002
Aphrodite1 gp140
VP4B gp075
pTD1 gp132
LAh10 gp209
CF8 gp049
pAEv1810 gp069
PS1 gp045
AH04 gp061
Kc263 gp140
N1M2 gp034
pEtSU gp284
OBP gp055
Klebsiella phage Miami
Pseudomonas phage EL
Erwinia phage vB_EamM_Joad
Photobacterium phage PDCC-1
Vibrio phage Aphrodite1
Vibrio phage VP4B
Vibrio phage pTD1
Aeromonas phage LAh10
Aeromonas phage CF8
Aeromonas phage pAEv1810
Aeromonas phage PS1
Erwinia phage AH04
Kosakonia phage Kc263
Klebsiella phage N1M2
Edwardsiella virus pEtSU
Pseudomonas phage OBP
DNA Polymerase vs. Whole Genome Tree

**C**

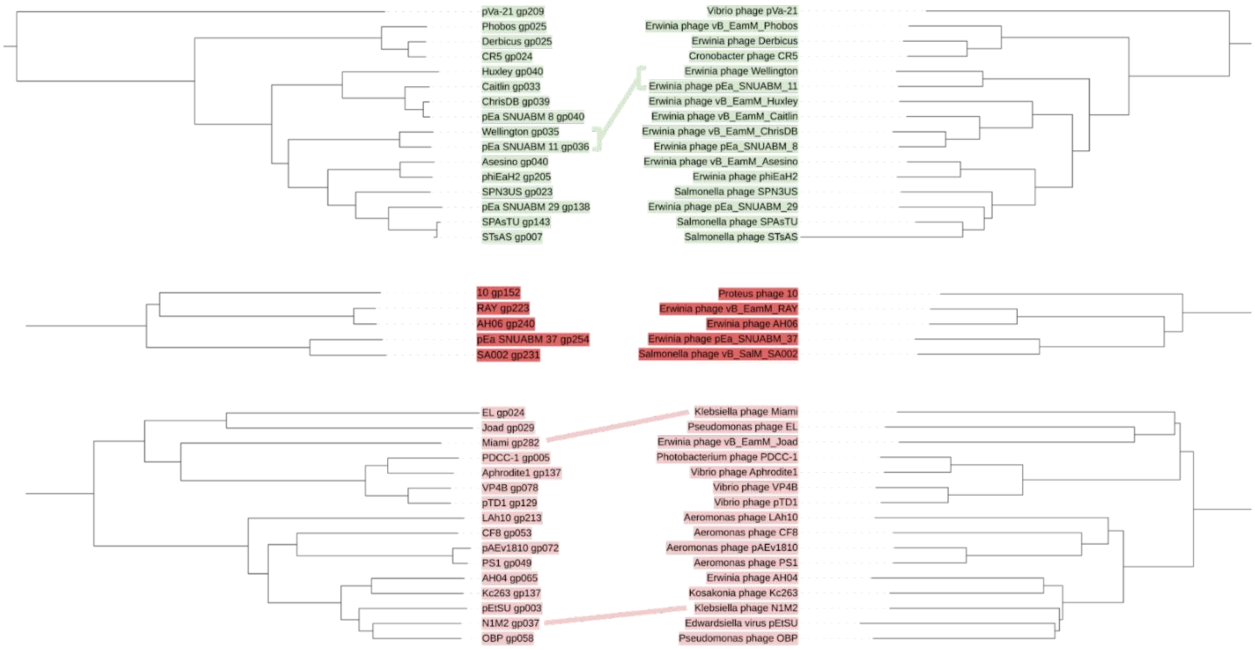


nvRNAP β' Subunit 1 vs. Whole Genome Tree

**D**

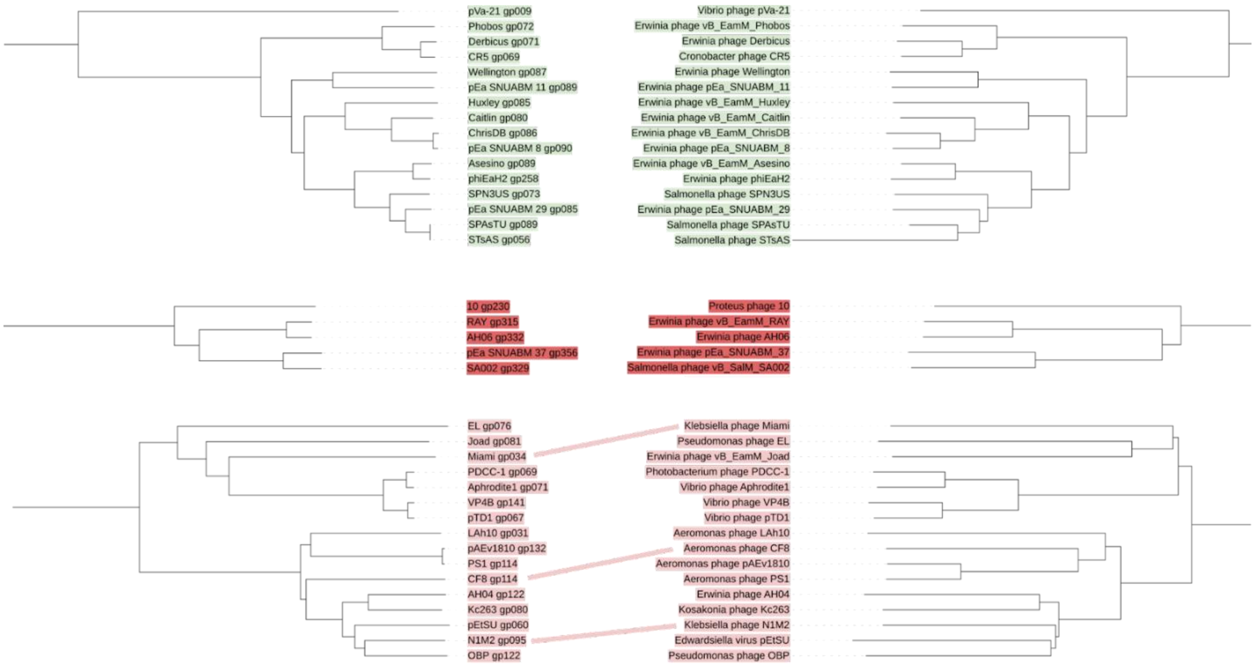


DnaB-like Helicase vs. Whole Genome Tree

**E**

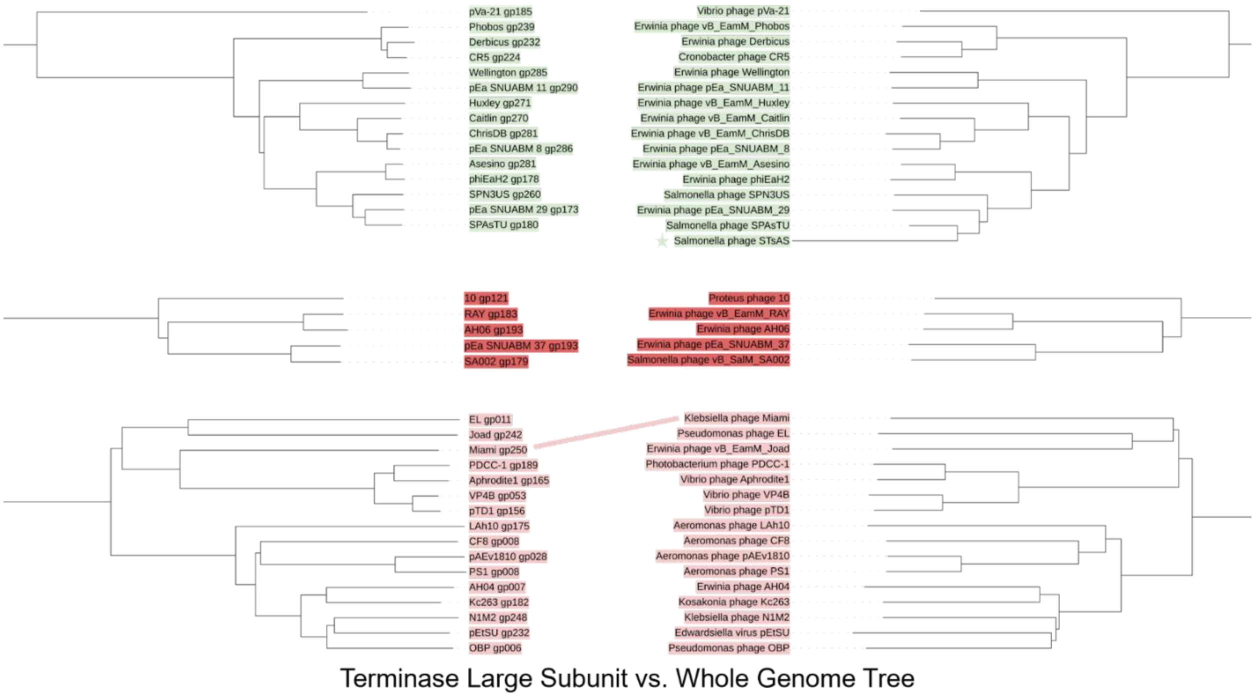


**Figure S2. Congruence between protein and whole genome based phylogenetics trees.** Mirrored trees comparing the whole genome phylogeny (right) with the phylogeny of individual proteins (left) were made for (A) the major capsid protein, (B) the DNA polymerase, (C) an RNA polymerase subunit, (D) the replicative helicase, and (E) the terminase large subunit. Minor discrepancies are pointed out with lines connecting the phage that have different branching patterns. In the case of the terminase, one phage was missing a homolog and is marked with a star. Clades are color-coded by predicted genus as in Figure 1. The light green, red, and light red clades were chosen for comparison because they were the largest clades and had the greatest potential to show how congruent the trees were (light green and light red) or were the clade containing RAY (red).

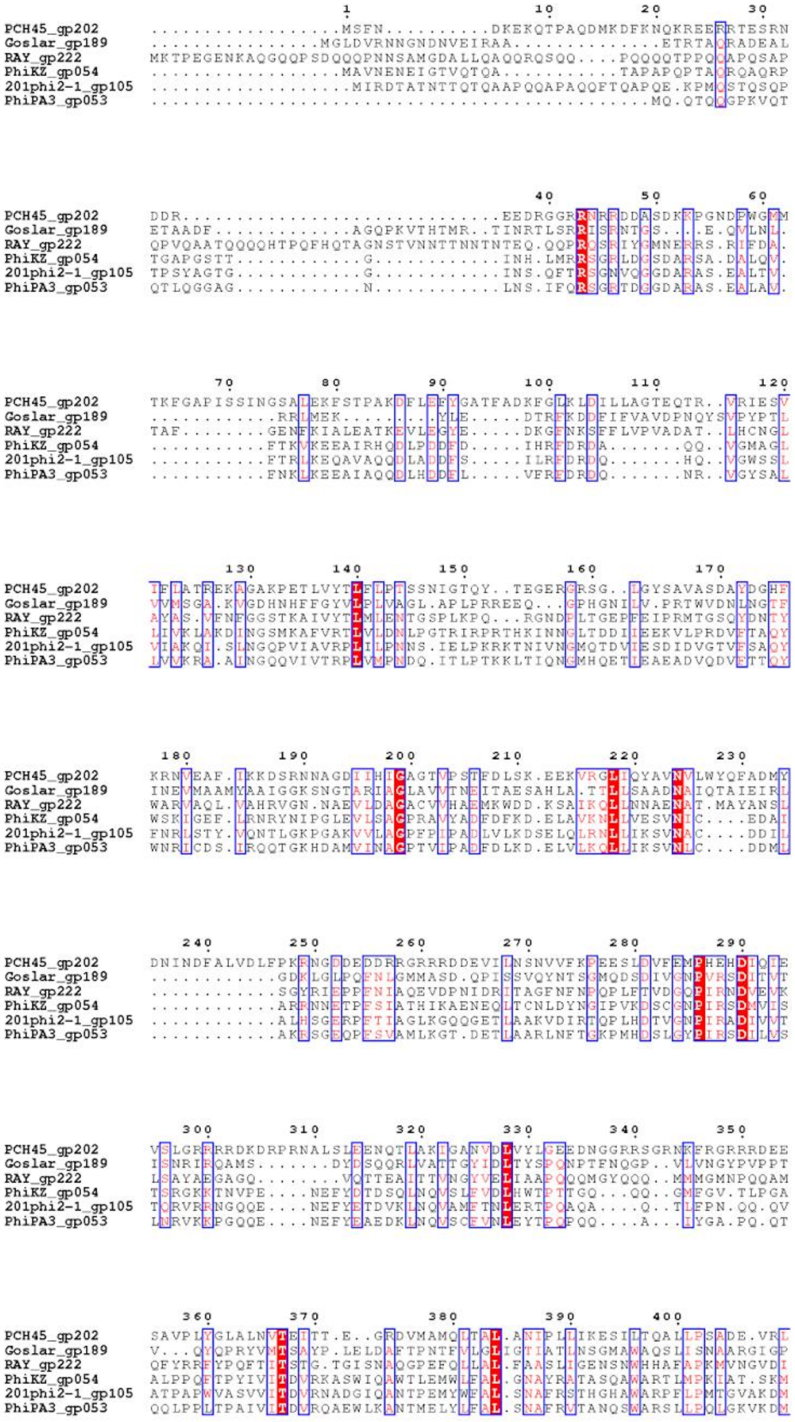
PCH45_gp202
Goslar_gp189
RAY_gp222
PhiKZ_gp054
201phi2-1_gp105
PhiPA3_gp053

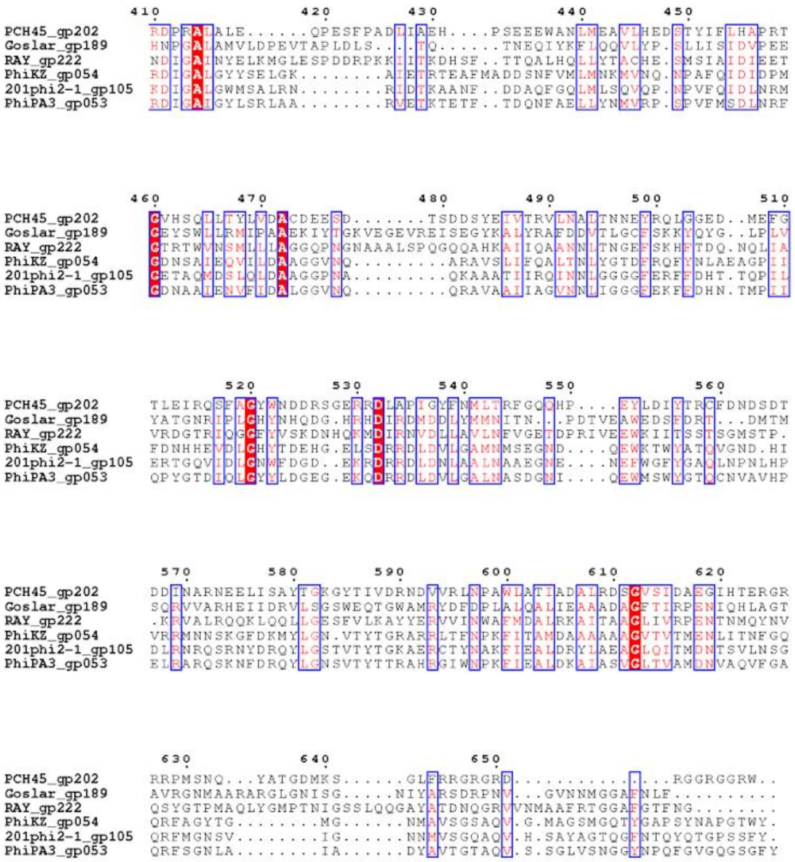


**Figure S3. Chimallin multiple sequence alignment.** Multiple sequence alignment of RAY gp222 with chimallin from previously published nucleus-forming phages.

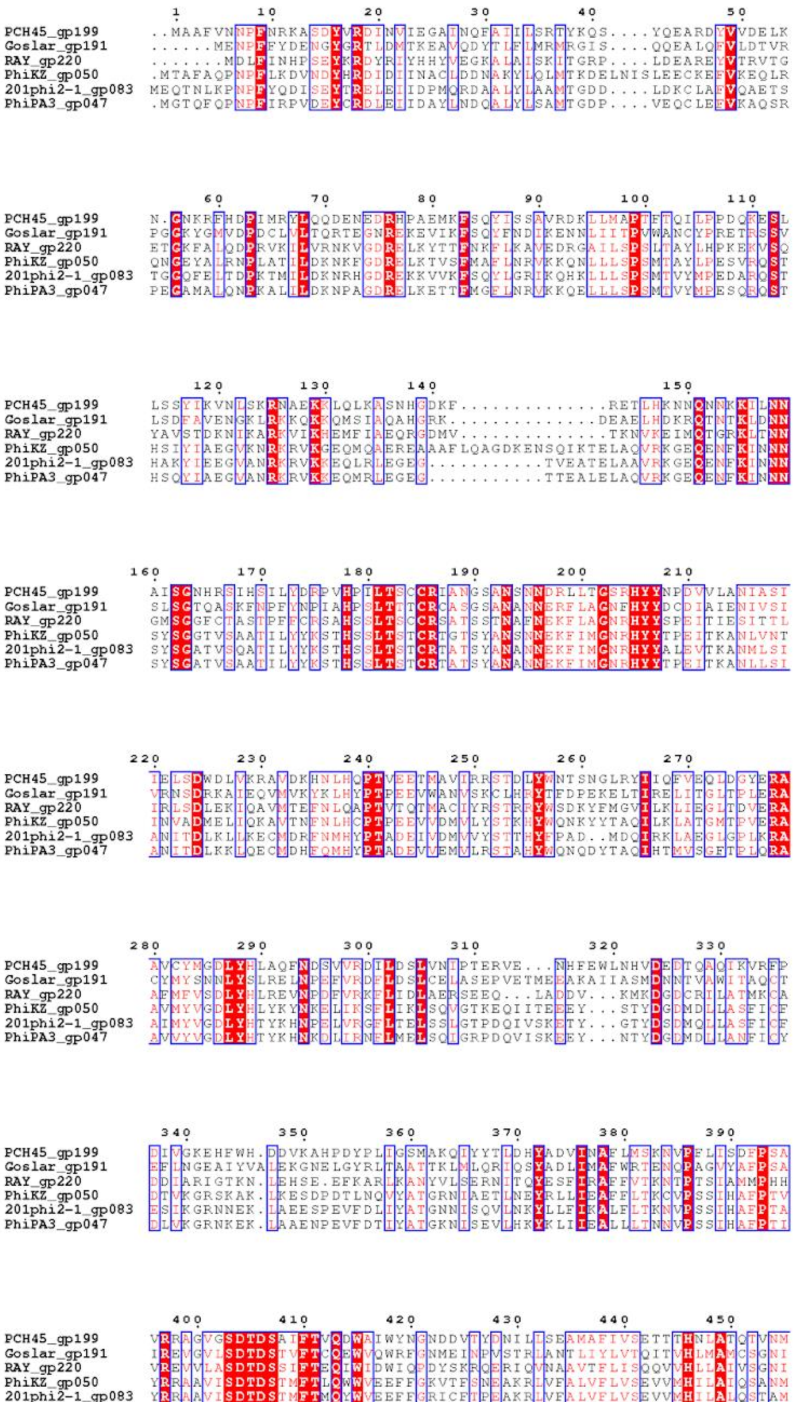
PCH45_gp199
Goslar_gp191
RAY_gp220
PhiKZ_gp050
201phi2-1_gp083
PhiPA3_gp047
1 10 20 30 40 50
60 70 80 90 100 110
120 130 140 150
160 170 180 190 200 210
220 230 240 250 260 270
280 290 300 310 320 330
340 350 360 370 380 390
400 410 420 430 440 450

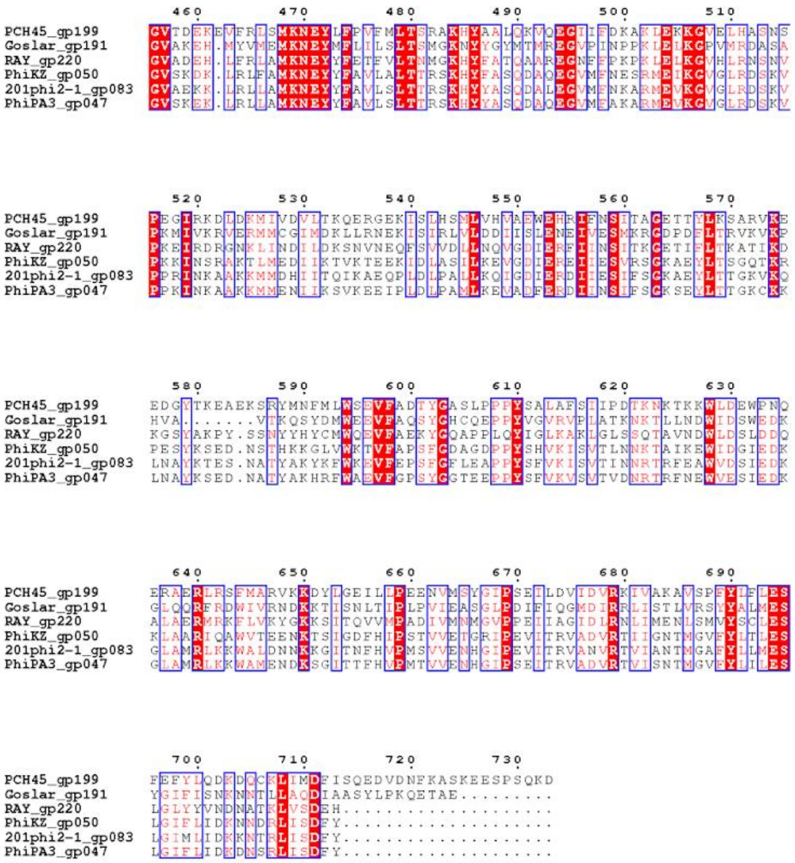

**Figure S4. DNA polymerase multiple sequence alignment.** Multiple sequence alignment of RAY gp220 and DNA polymerases from previously published nucleus-forming phages.

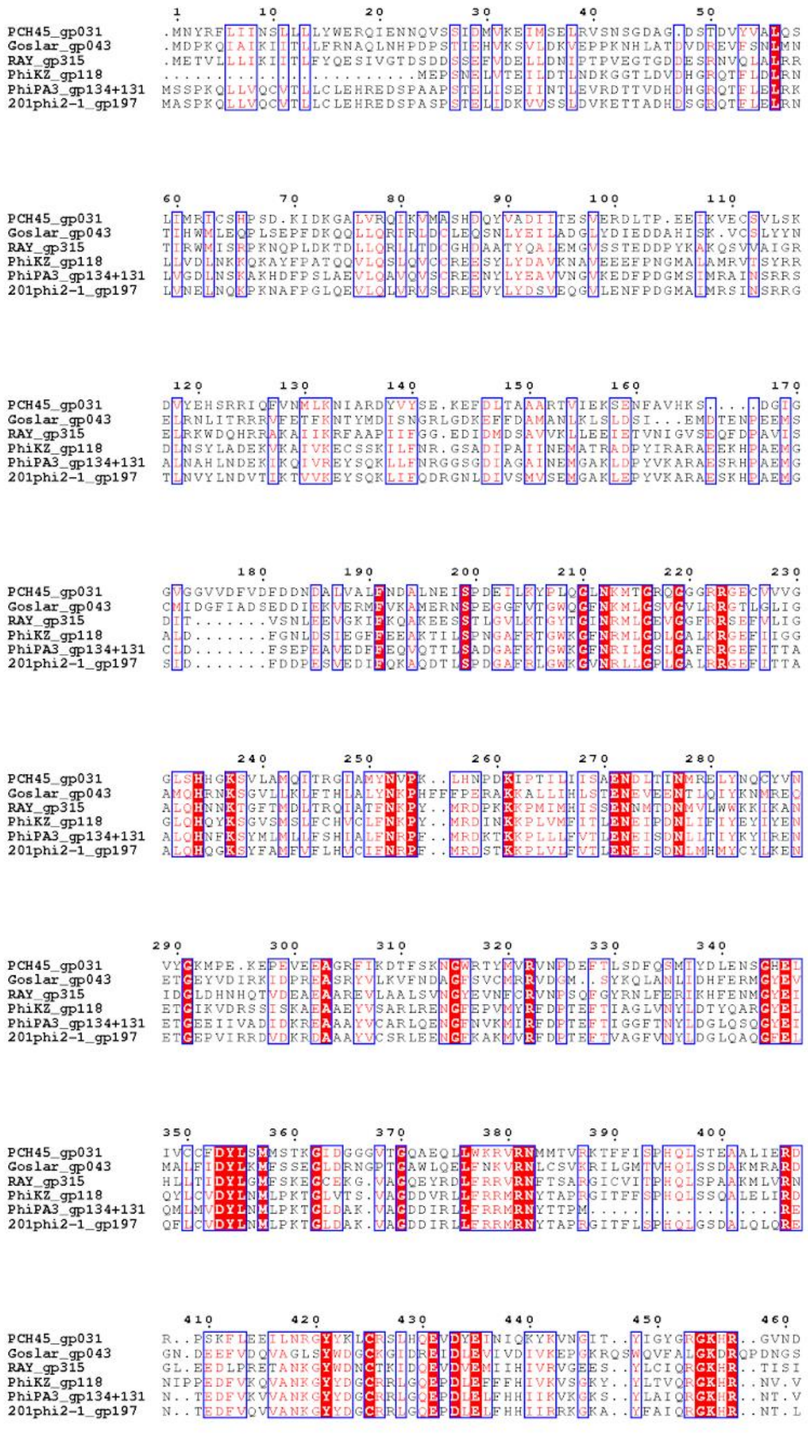
PCH45_gp031
Goslar_gp043
RAY_gp315
PhiKZ_gp118
PhiPA3_gp134+131
201phi2-1_gp197

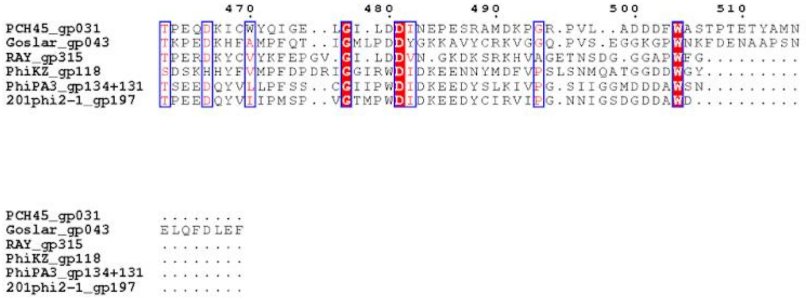


**Figure S5. Replicative (DnaB-like) helicase multiple sequence alignment.** Multiple sequence alignment of RAY gp315 with DNA helicases from previously published nucleus-forming phages.

A

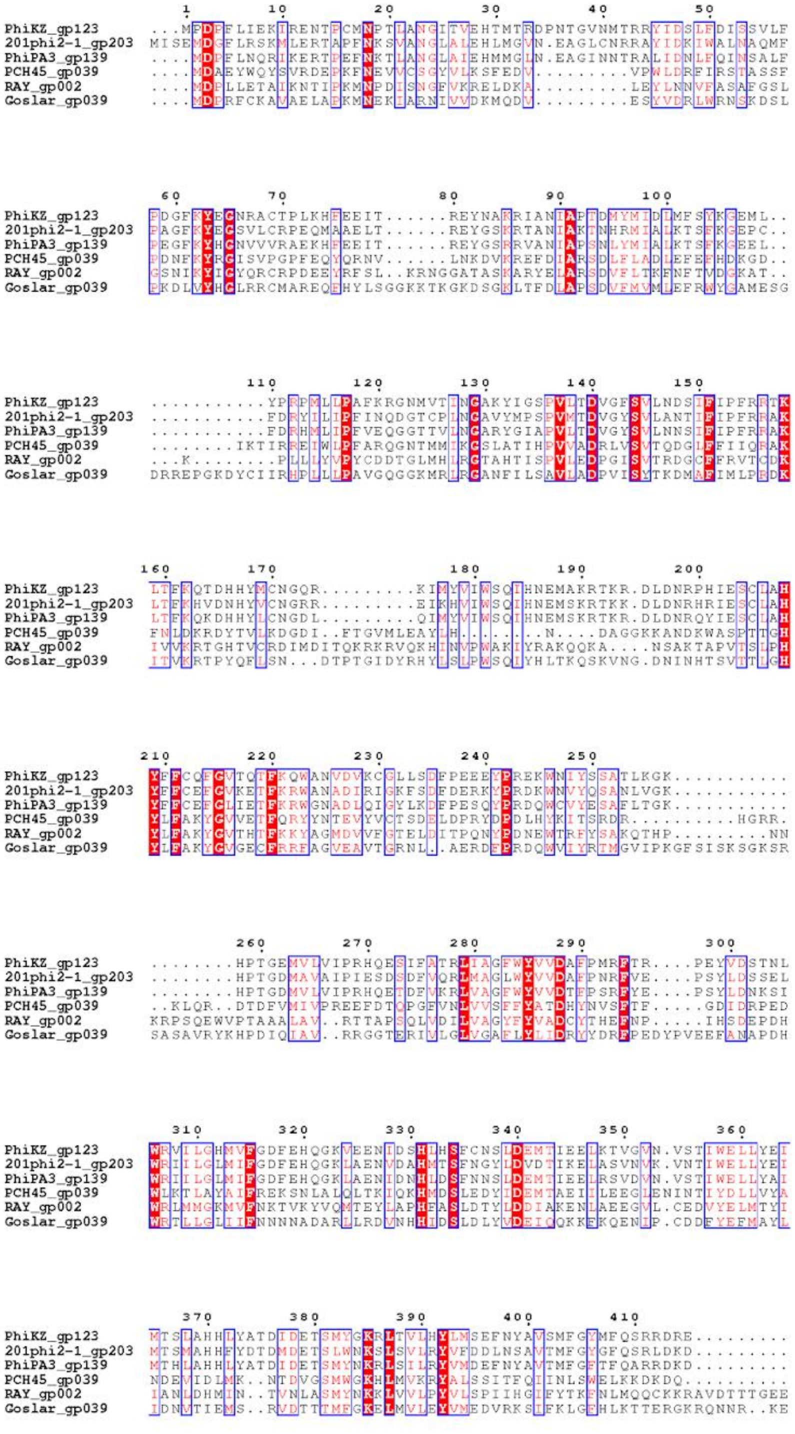
1 10 20 30 40 50
PhiKZ_gp123 ...MFDFFLIEKIRENTFCMNPTLANGITVEHTMTRDPNTGVNMTRRYIDSLFDISSVLF
201phi2-1_gp203 MISEMDGFLRSKMLERTAPENKSVANGLALEHLMGVN.EAGLCNRRAYIDKIWALNAQMF
PhiPA3_gp139 ....MDFFLNQRIKERTFEENKTLANGLAIEHMMGLN.EAGINNTRALIDNLFQINSALF
PCH45_gp039 ....MDAEYWQYSVRDEFKENEVVCGGYVLKSFEDV.........VPWLDRFIRSTASSF
RAY_gp002 ....MDFLLETAIKNTIFKMNPDISNGFVKRELDKA.........LEYLNNVFASAFGSL
Goslar_gp039 ....MDERFCKAVAELAFKMNEKIARNIVVDKMQDV.........ESYVDRLWRNSKDSL
60 70 80 90 100
PhiKZ_gp123 FDGFKYEGNRACTPLKHFEEIT......REYNAKRIANIAPTDMYMIDIMFSYKGEML..
201phi2-1_gp203 FAGFKYEGSVLCRPEQMAAELT......REYGSKRTANIAKTNHRMIALKTSFKGEPC..
PhiPA3_gp139 FEGFKYHGNVVVRAEKHFEEIT......REYGSRRVANIAPSNLYMIALKTSFKGEEL..
PCH45_gp039 FDNFKYRGISVPGPFEQYQRNV......LNKDVKREFDIARSDLFLADLEFEFHDKGD..
RAY_gp002 GSNIKYIGYQRCRPDEEYRFSL..KRNGGATASKARYEIARSDVFLTKFNFTVDGKAT..
Goslar_gp039 FKDLVYHGLRRCMAREQEHYLSGGKKTKGKDSGKLTFDIAPSDVFMVMIEFRWYGAMESG
110 120 130 140 150
PhiKZ_gp123 ...........YPRPMLLPAFKRGNMVTINGAKYIGSPVLTDVGFSVLNDSIFIPFRRTK
201phi2-1_gp203 ...........FDRYILIPFINQDGTCPLNGAVYMPSPVMTDVGYSVLANTIFIPFRRAK
PhiPA3_gp139 ...........FDRHMLIPFVEQGGTTVLNGARYGIAPVLTDVGYSVLNNSIFIPFRRAK
PCH45_gp039 ........IKTIRREIWLPFARQGNTMMIKGSLATIHPVVADRLVSVTQDGLFFIIQRAK
RAY_gp002 ...K........PLLLYVPYCDDTGLMHLRGTAHTISPVLEDPGISVTRDGCFFRVTGDK
Goslar_gp039 DRREPGKDYCIIRHPLLLPAVGQGGKMRLRGANFILSAVLADPVISYTKDMAFIMLPEDK
160 170 180 190 200
PhiKZ_gp123 LTFKQTDHHYMCNGQR.........KIMYVIWSQIHNEMAKRTKR.DLDNRPHIESCLAH
201phi2-1_gp203 LTFKHVDNHYVCNGRR.........EIKHVIWSQIHNEMSKRTKK.DLDNRHRIESCLAH
PhiPA3_gp139 LTFKQKDHHYLCNGDL.........QIMYVIWSQIHNEMSKRTKR.DLDNRQYIESCLAH
PCH45_gp039 FNLDKRDYTVLKDGDI..FTGVMLEAYLH.......N....DAGGKKANDKWASPTTGH
RAY_gp002 IVVKRTGHTVCRDIMDITQKRKRVQKHINVPWAKIYRAKQQKA....NSAKTAPVTSLPH
Goslar_gp039 ITVKRTPYQFLSN...DTPTGIDYRHYLSLPWSQIYHLTKQSKVNG.DNINHTSVTTLGH
210 220 230 240 250
PhiKZ_gp123 YFFCQFGVTQTFKQWANVDVKCGLLSDFPEEEYPREKWNIYSSATLKGK...........
201phi2-1_gp203 YFFCEFGVKETFKRWANADIRIGKFSDFDERKYPRDKWNVYQSANLVGK...........
PhiPA3_gp139 YFFCEFGLIETFKRWGNADLQIGYLKDFPESQYPRDQWCVYESAFLTGK...........
PCH45_gp039 YLFAKYGVVETFQRYYNTEVYVCTSDELDPRYDPDLHYKITSRDR..........HGRR.
RAY_gp002 YLFAKYGVTHTFKKYAGMDVVFGTELDITPQNYPDNEWTRFYSAKQTHP.........NN
Goslar_gp039 YIFAKYGVGECFRRFAGVEAVTGRNL..AERDFPRDQWVIYRTMGVIPKGFSISKSGKSR
260 270 280 290 300
PhiKZ_gp123 ........HPTGEMVLVIPRHQESIFATRLIAGFWYVVDAFPMRFTR....PEYVDSTNL
201phi2-1_gp203 ........HPTGDMAVAIPIESDSDFVQRLMAGLWYVVDAFPNRFVE....PSYLDSSEL
PhiPA3_gp139 ........HPTGDMVLVIPRHQETDFVKRLVAGFWYVVDTFPSRFYE....PSYLDNKSI
PCH45_gp039 ..KLQR..DTDFVMIVPREEFDTQPGFVNLVVSFPYATDHYNVSFTF.....GDIDRPED
RAY_gp002 KRPSQEWVPTAAALAV..RTTAPSQLVDILVAGYFYVADCYTHEFNP.....IHSDEPDH
Goslar_gp039 SASAVRYKHPDIQIAV..RRGGTERIVLGLVGAFLYLIDRYYDREPEDYPVEEFANAPDH
310 320 330 340 350 360
PhiKZ_gp123 WRVILGHMVFGDFEHQGKVEENIDSHIHSFCNSIDEMTIEELKTVGVN.VSTIWELLYEI
201phi2-1_gp203 WRIILGLMIFGDFEHQGKLAENVDAHMTSFNGYIDVDTIKELASVNVK.VNTIWELLYEI
PhiPA3_gp139 WRIILGLMIFGDFEHQGKLAENIDNHIDSFNNSIDEMTIEELRSVDVN.VSTIWELLYAI
PCH45_gp039 WLKTLAYAIFREKSNLALQLTKIQKHMDSLEDYIDEMTAEIIIEEGLENINTIYDLLVYA
RAY_gp002 WRLMMGKMVFNKTVKYVQMTEYLAPHFASLDTYLDDIAKENLAEEGVL.CEDVYELMTYI
Goslar_gp039 WRTLLGLIIFNNNNADARLLRDVNHHIDSLDLYVDEIQQKKEKQENIP.CDDFYEFMAYL
370 380 390 400 410
PhiKZ_gp123 MTSLAHHLYATDIDETSMYGKRLTVLHYLMSEFNYAVSMFGYMFQSRRDRE.........
201phi2-1_gp203 MTSMAHHFYDTDMDETSLWNKSLSVLRYVFDDLNSAVTMFGYGFQSRLDKD.........
PhiPA3_gp139 MTHLAHHLYATDIDETSMYNKRLSILRYVMDEFNYAVTMFGFTFQARRDKD.........
PCH45_gp039 NDEVIDLMK..NTDVGSMWGKHLMVKRYALSSITFQIINLSWELKKDKDQ..........
RAY_gp002 IANLDHMIN..TVNLASMYNKKLVVLPYVLSPIIHGIFYTKFNLMQQCKKRAVDTTTGEE
Goslar_gp039 IDNMTIEMS..RVDTTTMFGKELMVLEYVMEDVRKSIFKLGEHLKTTERGKRQNNR..KE

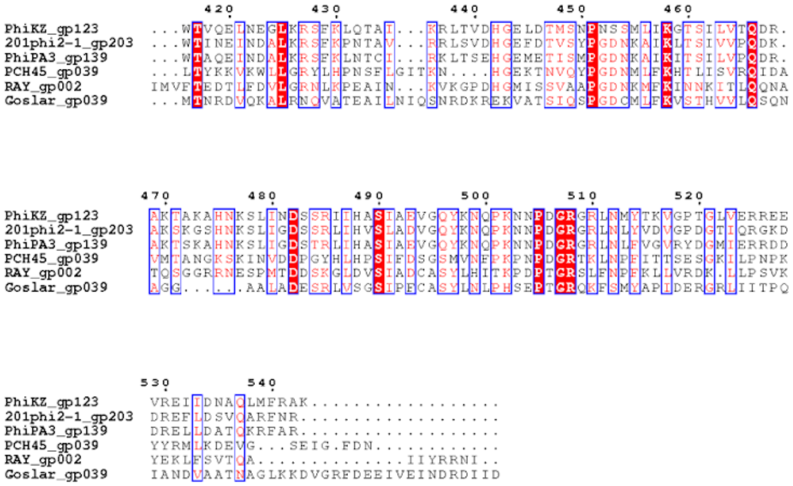
420 430 440 450 460
PhiKZ_gp123 ...WTVQELNEGLKRSFKLQTAI...KRLTVDHGELDTMSNPNSSMLIKGTSILVTQDR.
201phi2-1_gp203 ...WTINEINDALKRSFKPNTAV...RRLSVDHGEFDTVSYPGDNKAIKLTSIVVPQDK.
PhiPA3_gp139 ...WTAQEINDALKRSFKLNTCI...RKLTSEHGEMETISMPGDNKAIKITSILVPQDR.
PCH45_gp039 ...LTYKKVKWLLGRYLHPNSFLGITKN....HGEKTNVQYPGDNMLFKHTLISVRQIDA
RAY_gp002 IMVFTEDTLFDVLGRNLKPEAIN...KVKGPDHGMISSVAAPGDNKMFKINNKITLQQNA
Goslar_gp039 ...MTNRDVQKALRNQVATEAILNIQSNRDKREKVATSIQSPGDCMLFKVSTHVVLQSQN
470 480 490 500 510 520
PhiKZ_gp123 AKTAKAHNKSLINDSSRIIHASIAEVGQYKNQPKNNPDGRGRLNMYTKVGPTGLVERREE
201phi2-1_gp203 AKSKGSHNKSLIGDSSRLIHVSLADVGQYKNQPKNNPDGRGRLNLYVDVGPDGTIQRGKD
PhiPA3_gp139 AKTSKAHNKSLIGDSTRLIHASIAEVGQYKNQPKNNPDGRGRLNLFVGVRYDGMIERRDD
PCH45_gp039 VMTANGKSKINVDDPGYHLHPSIFDSGSMVNFPKPNPDGRTKLNPFITTSESGKILPNPK
RAY_gp002 TQSGGRRNESPMTDDSKGLDVSIADCASYLHITKPDPTGRSLFNPFKLLVRDK.LLPSVK
Goslar_gp039 AGG.....AALADESRLVSGSIPFCASYLNLPHSEPTGRQKFSMYAPIDERGRLIITPQ
530 540
PhiKZ_gp123 VREIIDNAQLMFRAK..................
201phi2-1_gp203 DREFLDSVQARFNR...................
PhiPA3_gp139 DRELLDATQKRFAR...................
PCH45_gp039 YYRMLKDEVG...SEIG.FDN............
RAY_gp002 YEKLFSVTQA..............IIYRRNI..
Goslar_gp039 IANDVAATNAGLKKDVGRFDEEIVEINDRDIID

B

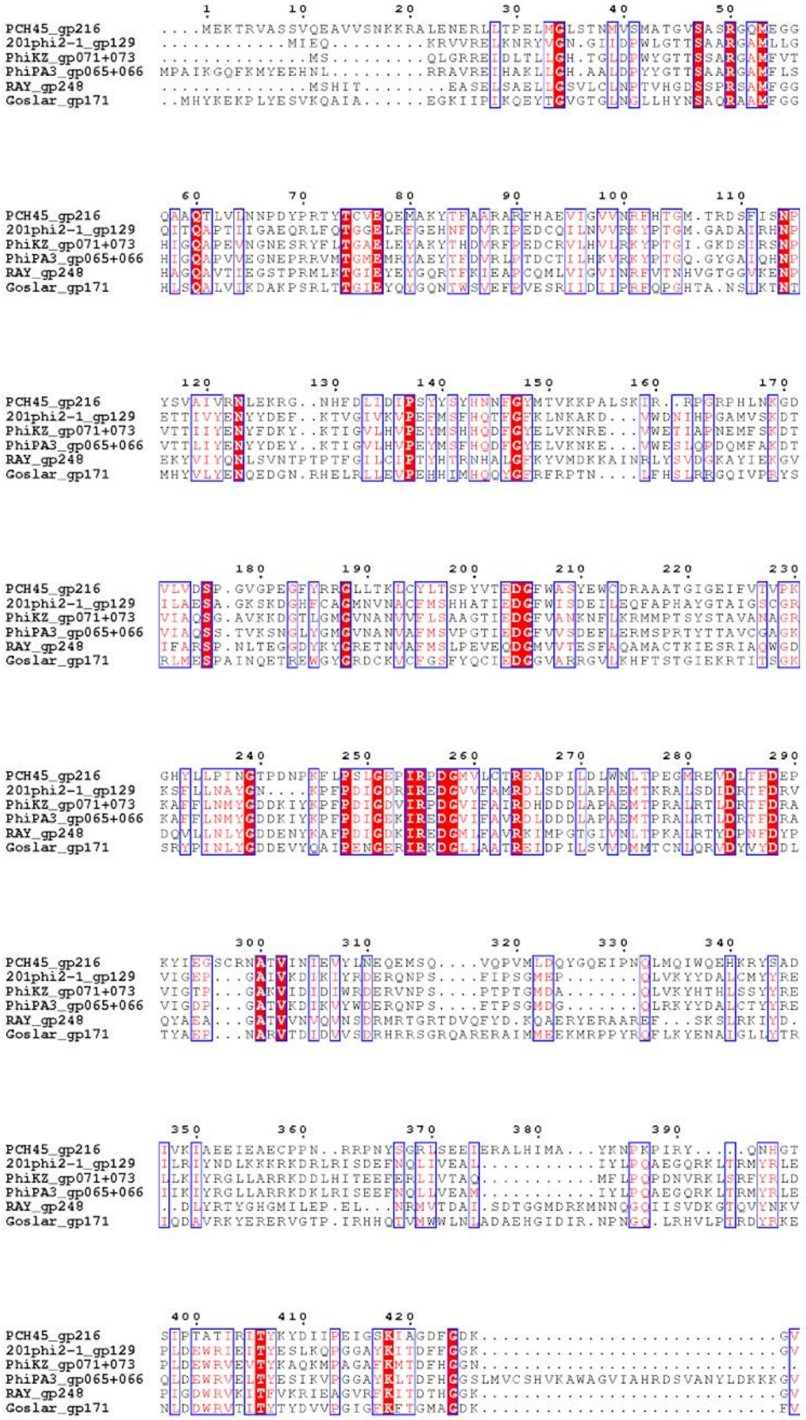
PCH45_gp216 ....MEKTRVASSVQEAVVSNKKRALENERLITPELMGLSTNMVSMATGVSASRGQMEGG
201phi2-1_gp129 ............MIEQ.........KRVVRELKNRYVGN.GIIDPWLGTTSSAARGAMLLG
PhiKZ_gp071+073 .............MS.........QLGRREIDLTLLGH.TGLDPWYGTTSSAARGAMFVT
PhiPA3_gp065+066 MPAIKGQFKMYEEHNL.........RRAVREIHAKLLGH.AALDPYYGTTSAARGAMFLS
RAY_gp248 .............MSHIT........EASELSAELLGSVLCLNPTVHGDSSPRSAMFGG
Goslar_gp171 ..MHYKEKPLYESVKQAIA......EGKIIPIKQEYTGVGTGLNGLLHYNSAQRAAMFGG
PCH45_gp216 QAAQTLVINNPDYPRTYTCVEQEMAKYTFAARARFHAEVIGVVNRFHTGM.TRDSFISNP
201phi2-1_gp129 QITQAPTIIGAEQRLFQTGGELRFGEHNFDVRIPEDCQILNVVRKYPTGM.GADAIRHNP
PhiKZ_gp071+073 HIGQAPEVNGNESRYFITGAELEYAKYTHDVRFPEDCRVLHVLRKYPTGI.GKDSIRSNP
PhiPA3_gp065+066 HIGQAPVVEGNEPRRVMTGMEMRYAEYTFDVRLPTDCTILHKVRKYPTGQ.GYGAIQHNP
RAY_gp248 HAGQAVTIEGSTPRMLKTGIEYEYGQRTFKIEAPCQMLVIGVINRFVTNHVGTGGVKENP
Goslar_gp171 HLSQALVIKDAKPSRLTTGIEYQYGQNTWSVEFPVESRIIDLIPRFQPGHTA.NSIKTNT
PCH45_gp216 YSVAIVRNLEKRG..NHFDLIDIPSYYSYHNNFGYMTVKKPALSKIR..RPGRPHLNKGD
201phi2-1_gp129 ETTIVYENYYDEF..KTVGIVKVPEFMSFHQTFGFKLNKAKD...VWDNIHPGAMVSKDT
PhiKZ_gp071+073 VTTIIYENYFDKY..KTIGVLHVPEYMSHHQDFGYELVKNRE...VWETIAPNEMFSKDT
PhiPA3_gp065+066 VTTLIYENYYDEY..KTIGVLHVPEYMSFHQDFGYELVKNKE...VWESLQPDQMFAKDT
RAY_gp248 EKYVIYQNLSVNTPTPTFGILGIPTYHTRNHALGFKYVMDKKAINRLYSVDGKAYIEKGV
Goslar_gp171 MHYVLYENQEDGN.RHELRLLEVPEHHIMHQQYGERFRPTN....IFHSLRRGQIVPRYS
PCH45_gp216 VLVDSP.GVGPEGFYRRGLLTKLCYLTSPYVTEDGFWASYEWCDRAAATGIGEIFVTVPK
201phi2-1_gp129 ILAESA.GKSKDGHFCAGMNVNACFMSHHATIEDGFWISDEILEQFAPHAYGTAIGSCGR
PhiKZ_gp071+073 VIAQSG.AVKKDGTLGMGVNANVVFLSAAGTIEDGFVANKNFLKRMMPTSYSTAVANAGR
PhiPA3_gp065+066 VIAQSS.TVKSNGLYGMGVNANVAFMSVPGTIEDGFVVSDEFLERMSPRTYTTAVCGAGK
RAY_gp248 IFARSP.NLTEGGDYKYGRETNVAFMSLPEVEQDGMVVTESFAQAMACTKIESRIAQWGD
Goslar_gp171 RLMESPAINQETREWGYGRDCKVCFGSFYQCIEDGGVARRGVLKHFTSTGIEKRTITSGK
PCH45_gp216 GHYLLPINGTPDNPKFLPSLGEPIRPDGMVLCTREADPILDLWNLTPEGMREVDLTFDEP
201phi2-1_gp129 KSFLLNAYGN....KPEPDIGDRIREDGVVFAMRDLSDDLAPAEMTKRALSDIDRTFDRV
PhiKZ_gp071+073 KAFFLNMYGDDKIYKPEPDIGDVIRPDGVIFAIRDHDDDLAPAEMTPRALRTLDRTFDRA
PhiPA3_gp065+066 KAFFLNMYGDDKIYKPEPDIGDKIRPDGVIFAVRDLDDDLAPAEMTPRALRTLDRTFDRA
RAY_gp248 DQVLLNLYGDDENYKAFPDIGDKIREDGMLFAVRKIMPGTGIVNLTPKALRTYDPNFDYP
Goslar_gp171 SRYPINLYGDDEVYQAIPENGERIRKDGLLAATREIDPILSVVDMMTCNLQRVDYVYDDL
PCH45_gp216 KYIEGSCRNATVINIEVYLNEQEMSQ....VQPVMLDQYGQEIPNQLMQIWQEHKRYSAD
201phi2-1_gp129 VIGEP...GAIVKDIKIYRDERQNPS....FIPSGMEP.......QLVKYYDALCMYYRE
PhiKZ_gp071+073 VIGTP...GAKVIDIDIWRDERVNPS....PTPTGMDA.......QLVKYHTHLSSYYRE
PhiPA3_gp065+066 VIGDP...GATVKDIKVYWDERQNPS....FTPSGMDG.......QLRKYYDALCTYYRE
RAY_gp248 QYAEA...GATVVNVQVNSDRMRTGRTDVQFYD.KQAERYERAAREF...SKSLRKIYD.
Goslar_gp171 TYAEP...NARVTDIDVVSDRHRRSGRQARERAIMMEEKMRPPYRQFLKYENAIGLLYTR
PCH45_gp216 IVKIAEEIEAECPPN..RRPNYSGRLSEEIERALHIMA...YKNPKPIRY....QNHGT
201phi2-1_gp129 ILRIYNDLKKKRKDRLRISDEFNQLIVEAL...........IYLPQAEGQRKLTRMYRLE
PhiKZ_gp071+073 LLKIYRGLLARRKDDLHITEEFERLIVTAQ...........MFLPQPDNVRKLSRFYRLD
PhiPA3_gp065+066 IIKIYRGLLARRKDKLRISEEFNQLLVEAM...........IYLPQAEGQRKLTRMYRLE
RAY_gp248 ..DLYRTYGHGMILEP.EL...NRMVTDAI.SDTGGMDRKMNNQGQIISVDKGTQVYNKV
Goslar_gp171 IQDAVRKYERERVGTP.IRHHQTVMWWLNLADAEHGIDIR.NPNGQ.LRHVLPTRDYRKE
PCH45_gp216 SIPTATIRITYKYDIIPEIGSKIAGDFGDK...........................GV
201phi2-1_gp129 PLDEWRIEITYESLKQPGGAYKITDFFGGK...........................GV
PhiKZ_gp071+073 PLDEWRVEVTYKAQKMPAGAFKMTDFHGGN.............................
PhiPA3_gp065+066 QLDEWRVELTYESIKVPGGAYKLTDFHGGSLMVCSHVKAWAGVIAHRDSVANYLDKKKGV
RAY_gp248 PIGDWRVKITFVKRIEAGVRFKITDTHGGK...........................GV
Goslar_gp171 NLDDWRVTITYTYDVVPGIGFKFTGMAGDK...........................FV

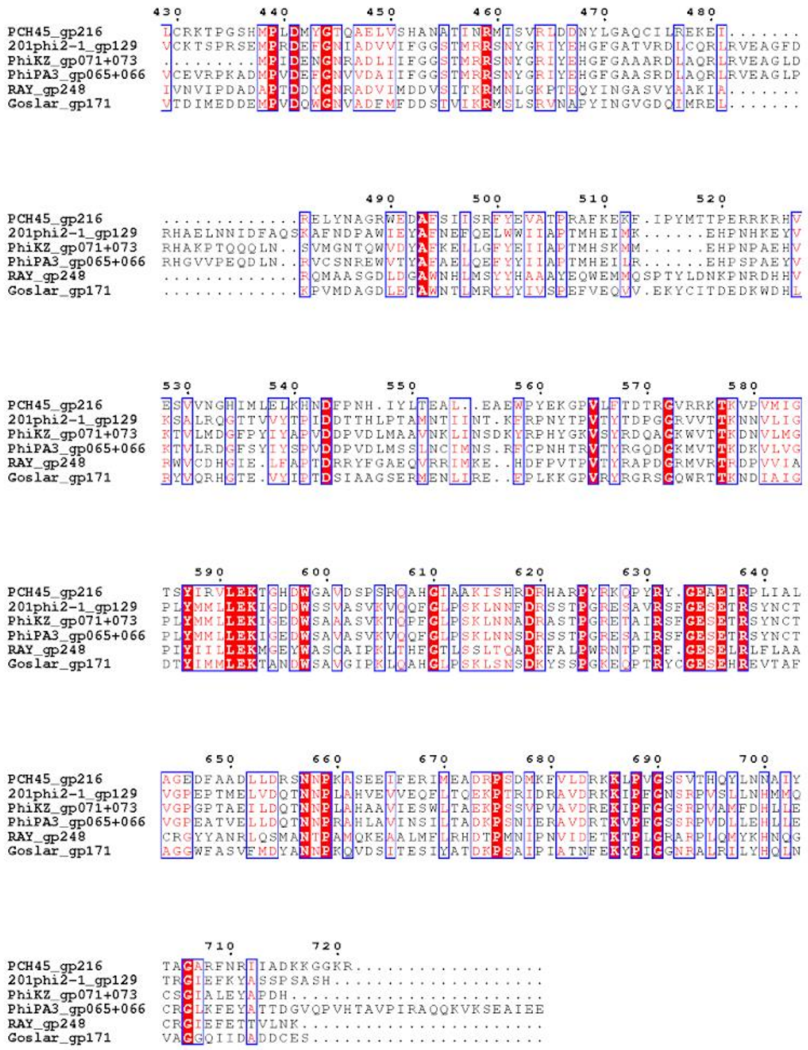
430 440 450 460 470 480
PCH45_gp216 LCRKTPGSHMPLDMYGTQAELVSHANATINRMISVRLDDNYLGAQCILREKEI.......
201phi2-1_gp129 VCKTSPRSEMPRDEFGNIADVVIFGGSTMRRSNYGRIYEHGFGATVRDLCQRLRVEAGFD
PhiKZ_gp071+073 .........MPIDENGNRADLIIFGGSTMRRSNYGRIYEHGFGAAARDLAQRLRVEAGLD
PhiPA3_gp065+066 VCEVRPKADMPVDEFGNVVDAIIFGGSTMRRSNYGRIYEHGFGAASRDLAQRLRVEAGLP
RAY_gp248 IVNVIPDADAPTDDYGNRADVIMDDVSITKRMNLGKPTEQYINGASVYAAKIA.......
Goslar_gp171 VTDIMEDDEMPVDQWGNVADFMFDDSTVIKRMSLSRVNAPYINGVGDQIMREL.......
490 500 510 520
PCH45_gp216 .............RELYNAGRWEDAFSIISRFYEVATPRAFKEKF.IPYMTTPERRKRHV
201phi2-1_gp129 RHAELNNIDFAQSKAFNDPAWIEYAFNEFQELWWIIAPTMHEIMK......EHPNHKEYV
PhiKZ_gp071+073 RHAKPTQQQLN..SVMGNTQWVDYAFKELLGFYEIIAPTMHSKMM......EHPNPAEHV
PhiPA3_gp065+066 RHGVVPEQDLN..RVCSNREWVTYAFAELQEFYYIIAPTMHEILR......EHPSPAEYV
RAY_gp248 .............RQMAASGDLDGAWNHLMSYYHAAAYEQWEMMQSPTYLDNKPNRDHHV
Goslar_gp171 .............KPVMDAGDLETAWNTLMRYYYIVSPEFVEQVV.EKYCITDEDKWDHL
530 540 550 560 570 580
PCH45_gp216 ESVVNGHIMLELKHNDFPNH.IYLTEAL..EAEWPYEKGPVLFTDTRGVRRKTKVPVMIG
201phi2-1_gp129 KSALRQGTTVVYTPIDDTTHLPTAMNTIINT.KFRPNYTPVTYTDPGGRVVTTKNNVLIG
PhiKZ_gp071+073 KTVLMDGFPYIYAPVDDPVDLMAAVNKLINSDKYRPHYGKVSYRDQAGKWVTTKDNVLMG
PhiPA3_gp065+066 KTVLRDGFSYIYSPVDDPVDLMSSLNCIMNS.RFCPNHTRVTYRGQDGKMVTTKDKVLVG
RAY_gp248 RWVCDHGIE.LFAPTDRRYFGAEQVRRIMKE..HDFPVTPVTYRAPDGRMVRTRDPVVIA
Goslar_gp171 RYVQRHGTE.VYIPTDSIAAGSERMENLIRE..FPLKKGPVRYRGRSGQWRTTKNDIAIG
590 600 610 620 630 640
PCH45_gp216 TSYIRVLEKTGHDWGAVDSPSRQAHGIAAKISHRDRHARPYRKQPYRY.GEAEIRPLIAL
201phi2-1_gp129 PLYMMLLEKIGDDWSSVASVKVQQFGLPSKLNNFDRSSTPGRESAVRSFGESETRSYNCT
PhiKZ_gp071+073 PLYMMLLEKIGEDWSAAASVKTQPFGLPSKLNNADRASTPGRETAIRSFGESETRSYNCT
PhiPA3_gp065+066 PLYMMLLEKIGEDWSAVASVKVQQFGLPSKLNNSDRSSTPGRESAIRSFGESETRSYNCT
RAY_gp248 PIYIILLEKMGEYWASCAIPKLTHFGTLSSLTQADKFALPWRNTPTRF.GESELRLFLAA
Goslar_gp171 DTYIMMLEKTANDWSAVGIPKLQAHGLPSKLSNSDKYSSPGKEQPTRYCGESEHREVTAF
650 660 670 680 690 700
PCH45_gp216 AGEDFAADLLDRSNNPKASEEIFERIMEADRPSDMKFVLDRKKLPVGSSVTHQYLNNAIY
201phi2-1_gp129 VGPEPTMELVDQTNNPLAHVEVVEQFLTQEKPTRIDRAVDRKKIPFGNSRPVSLLNHMMQ
PhiKZ_gp071+073 VGPGPTAEILDQTNNPLAHAAVIESWLTAEKPSSVPVAVDREKIPFGGSRPVAMFDHLLE
PhiPA3_gp065+066 VGPEATVELLDQTNNPRAHLAVINSILTADKPSNIERAVDRTKVPFGSSRPVDLLEHLLE
RAY_gp248 CRGYYANRLQSMANTPAMQKEAALMFLRHDTPMNIPNVIDETKTPLGRARPLQMYKHNQG
Goslar_gp171 AGGWFASVFMDYANNPKQVDSITESIYATDKPSAIPIATNFEKYPIGGNRALRILYHQLN
710 720
PCH45_gp216 TACARFNRIIADKKGGKR..................
201phi2-1_gp129 TRGIEFKYASSPSASH....................
PhiKZ_gp071+073 CSGIALEYAPDH........................
PhiPA3_gp065+066 CRGLKFEYATTDGVQPVHTAVPIRAQQKVKSEAIEE
RAY_gp248 CRGIEFETTVLNK.......................
Goslar_gp171 VAGGQIIDADDCES......................

C

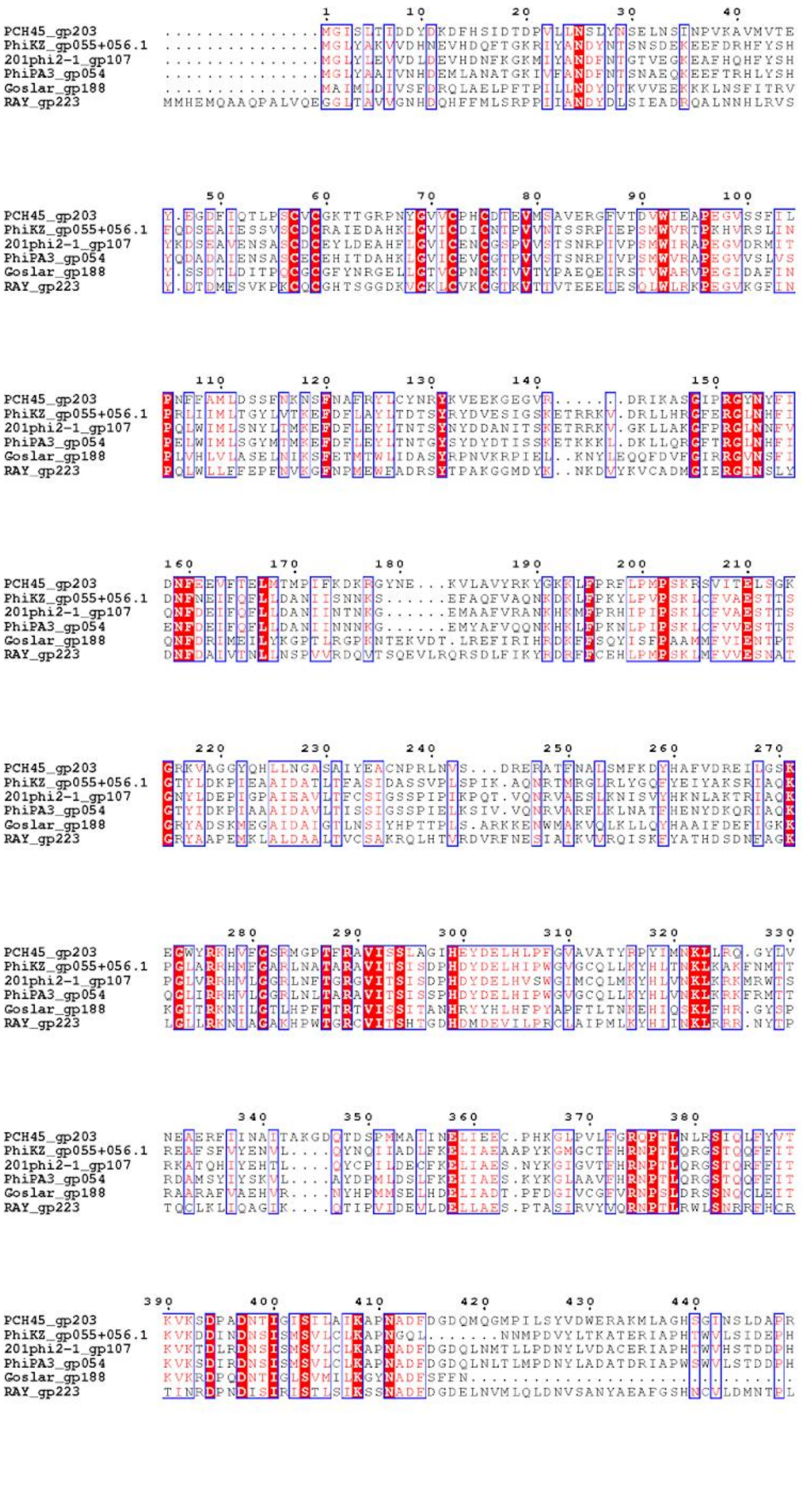
PCH45_gp203
PhiKZ_gp055+056.1
201phi2-1_gp107
PhiPA3_gp054
Goslar_gp188
RAY_gp223

```
                      450        460        470
                        .          .          .
PCH45_gp203          SLNRNLVMPDTDFSTLNNFMYGNSR...........
PhiKZ_gp055+056.1    EISGNLELQGPVVETIINWAHEKYLPPLEEWLKAK.
201phi2-1_gp107      QLSGNLELQGPVVETVVNYGHEDYLPPLEEWLAGH.
PhiPA3_gp054         EISGNLELQGPVVETIINFAHEDYLPPLEEWLKMAA
Goslar_gp188         ....................................
RAY_gp223            KISGDVGLPGTLISTMNRWMYSND............
```

D

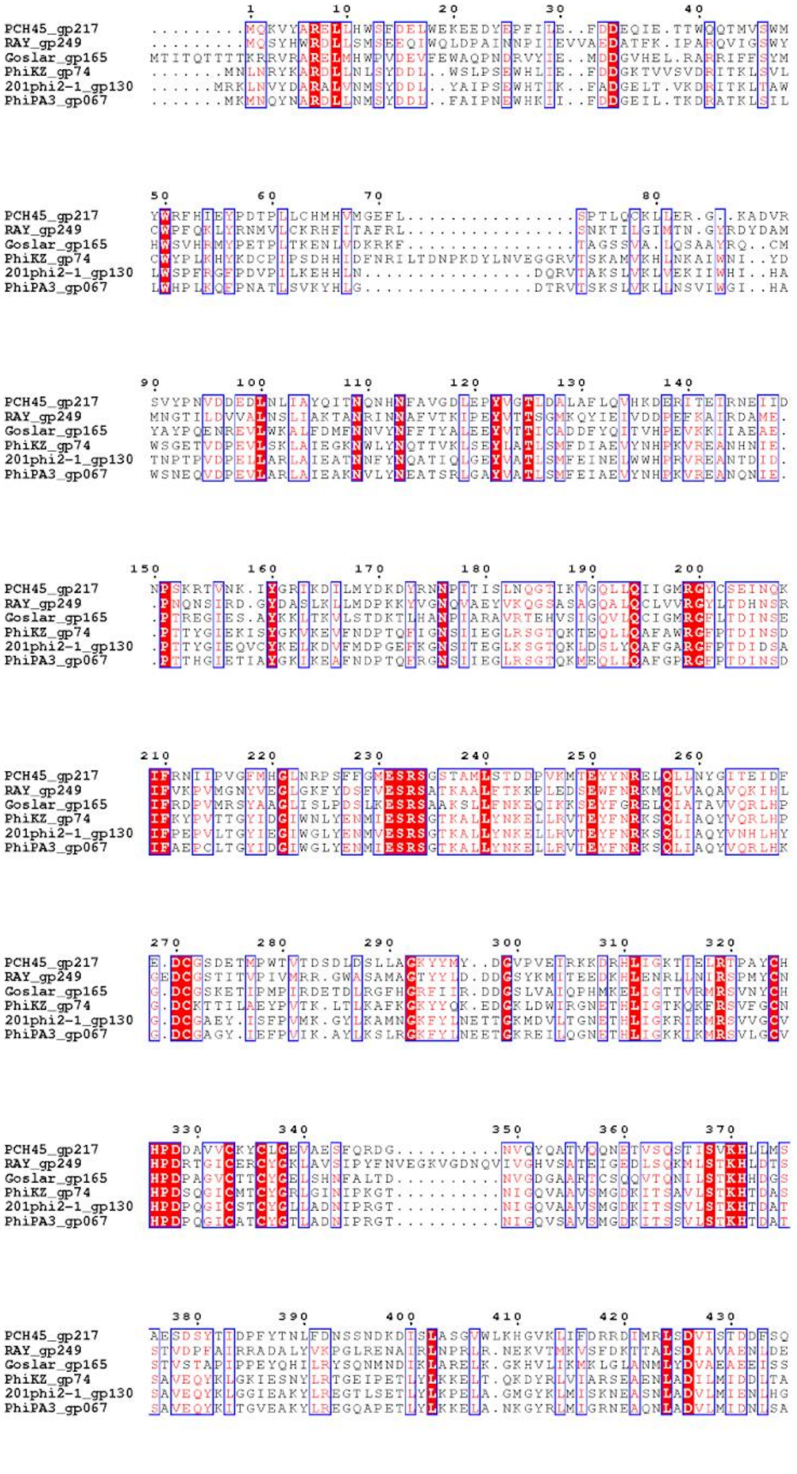
PCH45_gp217
RAY_gp249
Goslar_gp165
PhiKZ_gp74
201phi2-1_gp130
PhiPA3_gp067

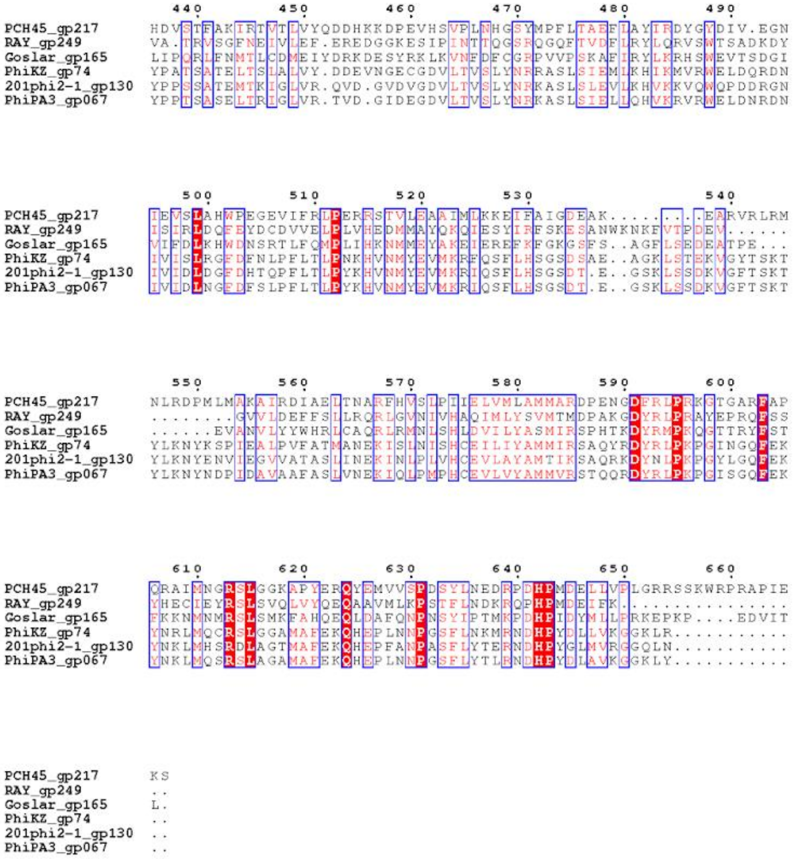

**Figure S6. Non-virion RNA polymerase multiple sequence alignment.** Multiple sequence alignments of RAY gp002 (A), gp248 (B), gp223 (C), and gp249 (D) with nvRNAP subunits from previously published nucleus-forming phages. These RAY proteins are homologs of known phage msRNAP subunits ΦKZ gp123, gp71-73, gp55-56.1, and gp74, respectively (de Martín Garrido et al., 2021).

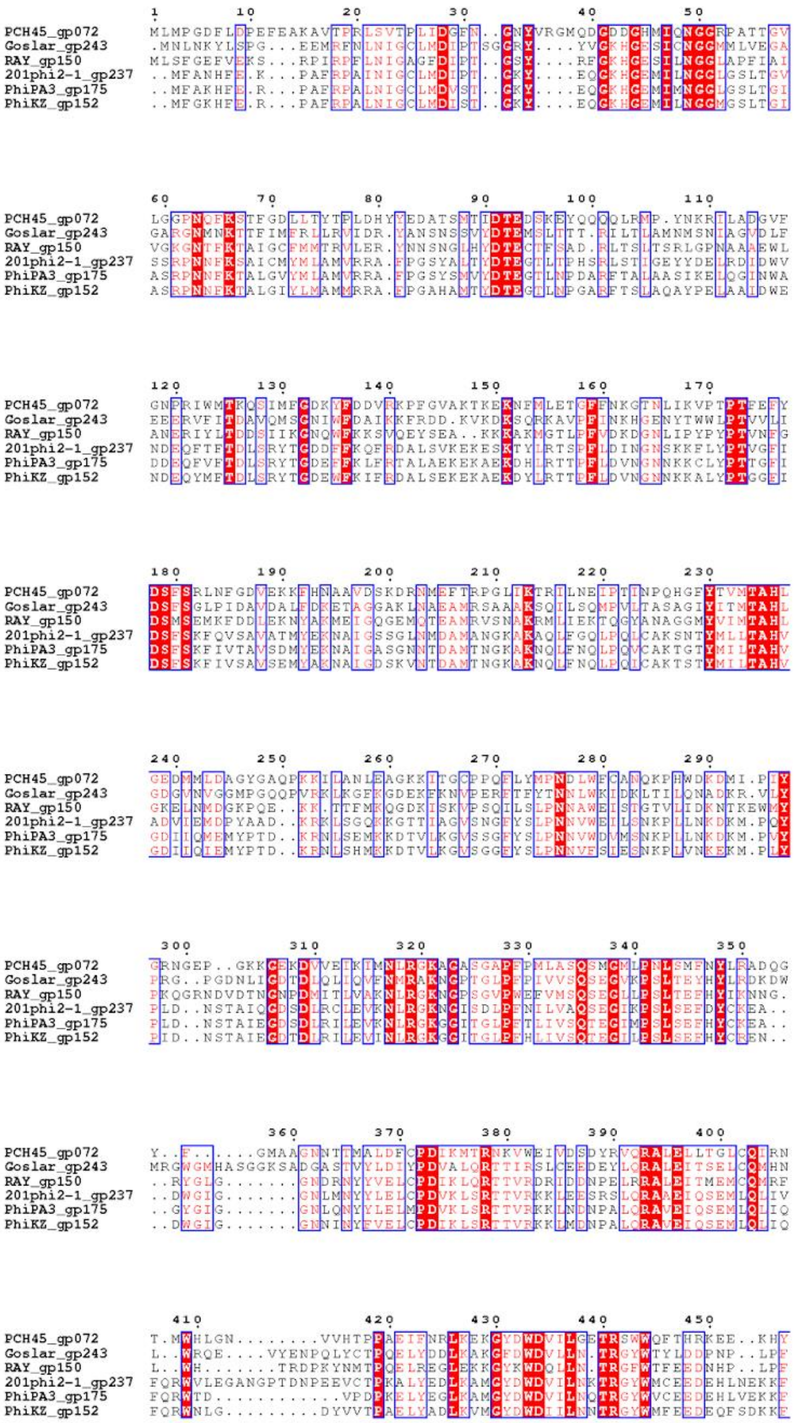
PCH45_gp072
Goslar_gp243
RAY_gp150
201phi2-1_gp237
PhiPA3_gp175
PhiKZ_gp152

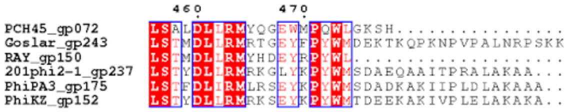


**Figure S7. UvsX/RecA multiple sequence alignment.** Multiple sequence alignment of RAY gp150 with UvsX from previously published nucleus-forming phages. UvsX is a RecA homolog.

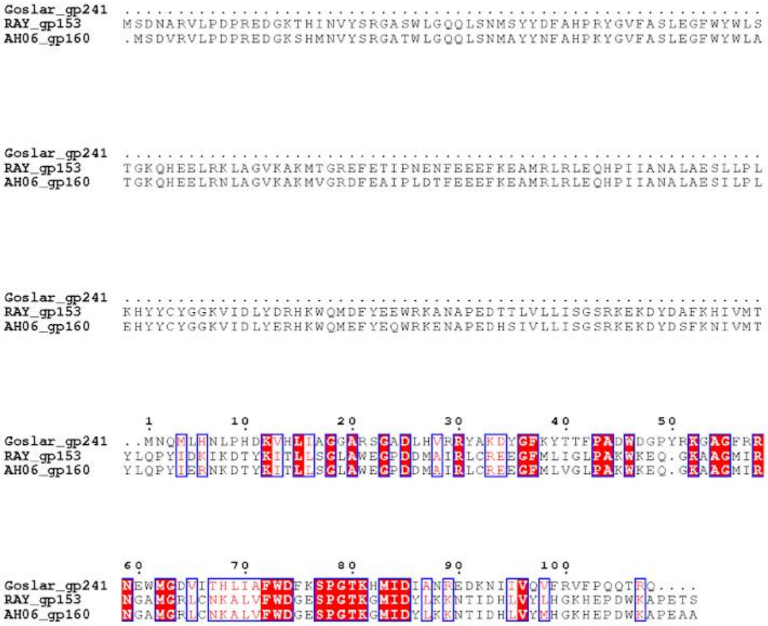


**Figure S8. DprA multiple sequence alignment.** Multiple sequence alignment of RAY gp153 with DprA from previously published nucleus-forming phage Goslar and RAY close relative AH06. DprA is a RecA-associated protein involved in recombination. Since it is not part of the core genome, it was not present in most of the previously-studied nucleus-forming phages used for other MSA analysis.

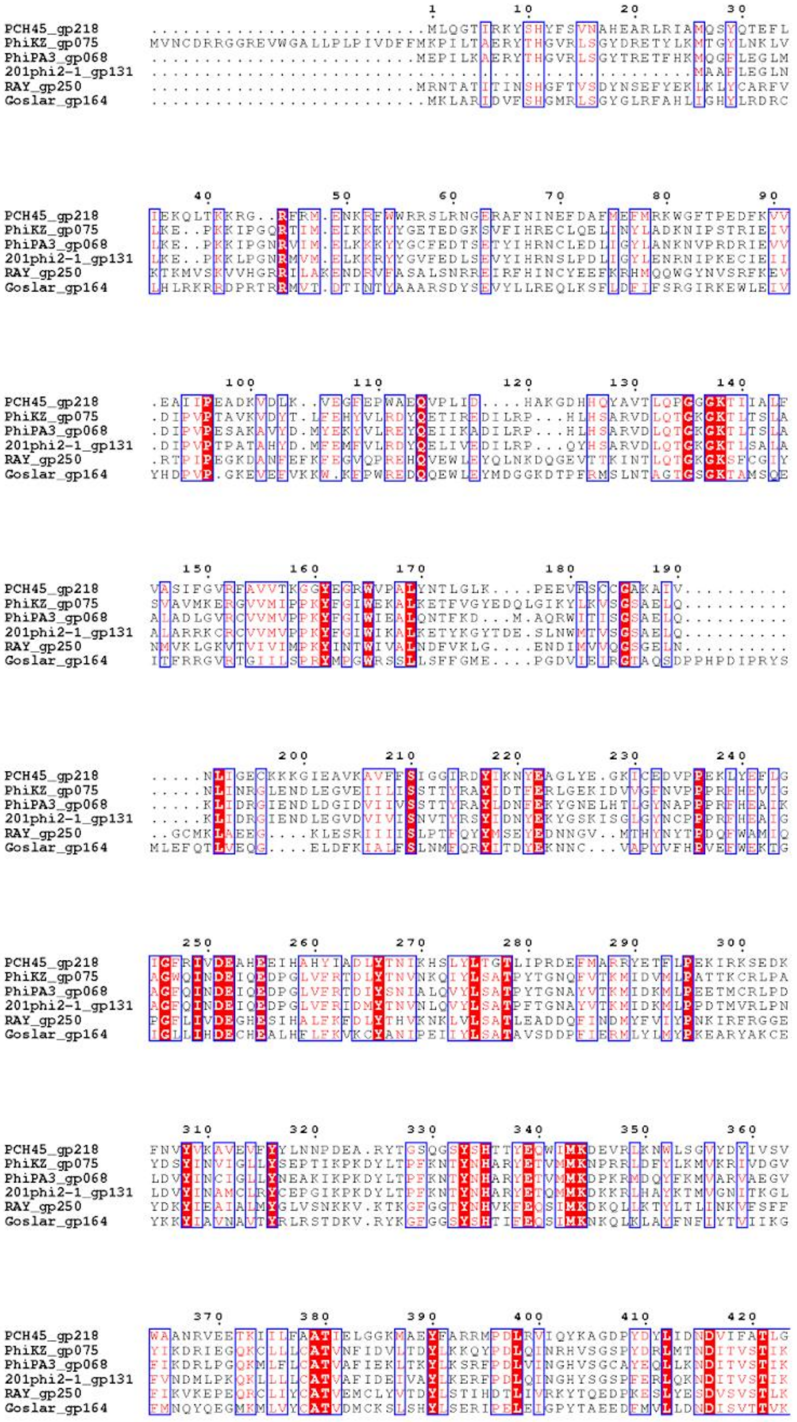
PCH45_gp218
PhiKZ_gp075
PhiPA3_gp068
201phi2-1_gp131
RAY_gp250
Goslar_gp164

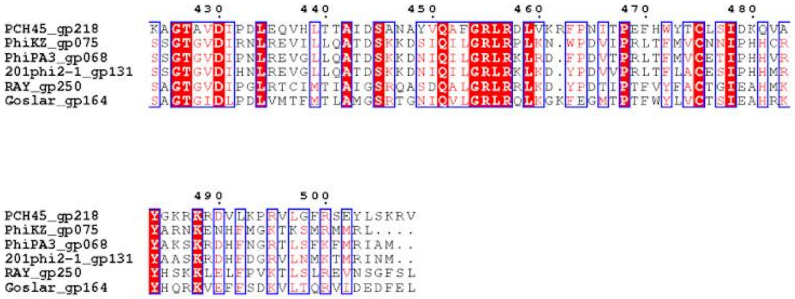


**Figure S9. Non-virion SF2 helicase multiple sequence alignment.** Multiple sequence alignment of RAY gp250 with SF2 helicase homologs from previously published nucleus-forming phages. RAY gp250 localized in the phage nucleus (Fig. 4B) as opposed to RAY gp131, another SF2 helicase homolog which localized similarly to capsids (Fig. 5B), leading us to call gp250 and its homologs the "non-virion" SF2 helicase.

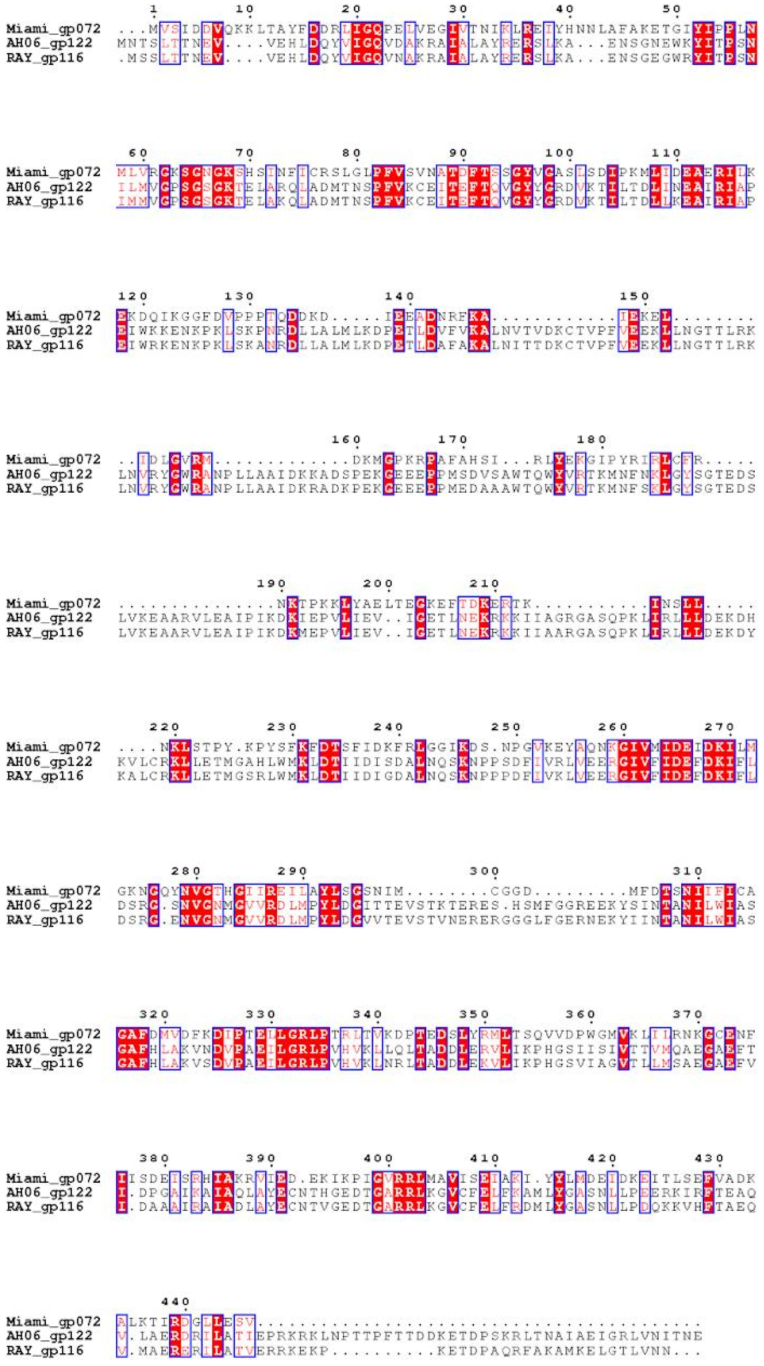

**Figure S10. HslU-like protease multiple sequence alignment.** Multiple sequence alignment of RAY gp116 with HslU-like protease homologs from RAY's close relative AH06 and its distant relative Miami. HslU is not part of the core genome and was not present in any of the previously-studied nucleus-forming phages used for other MSA analysis.

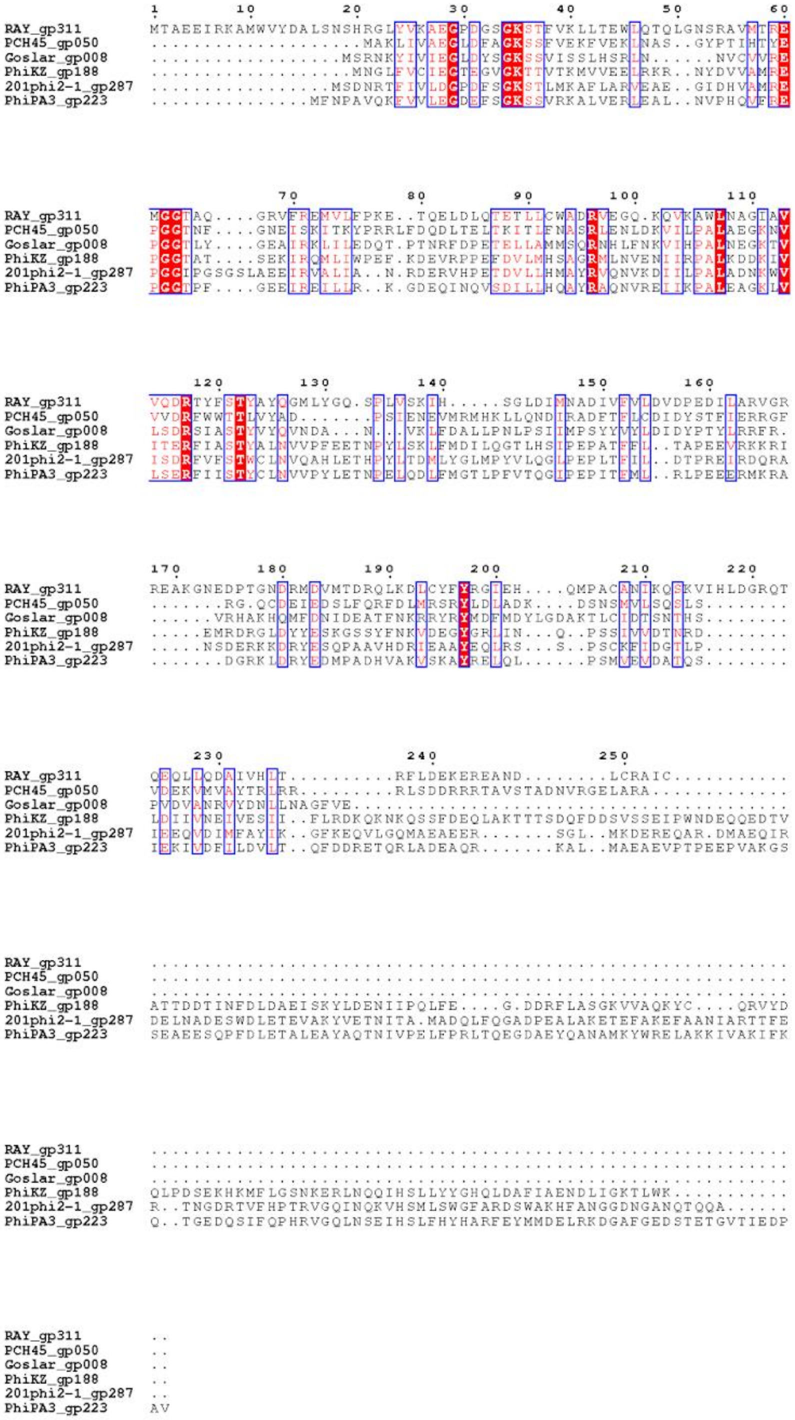

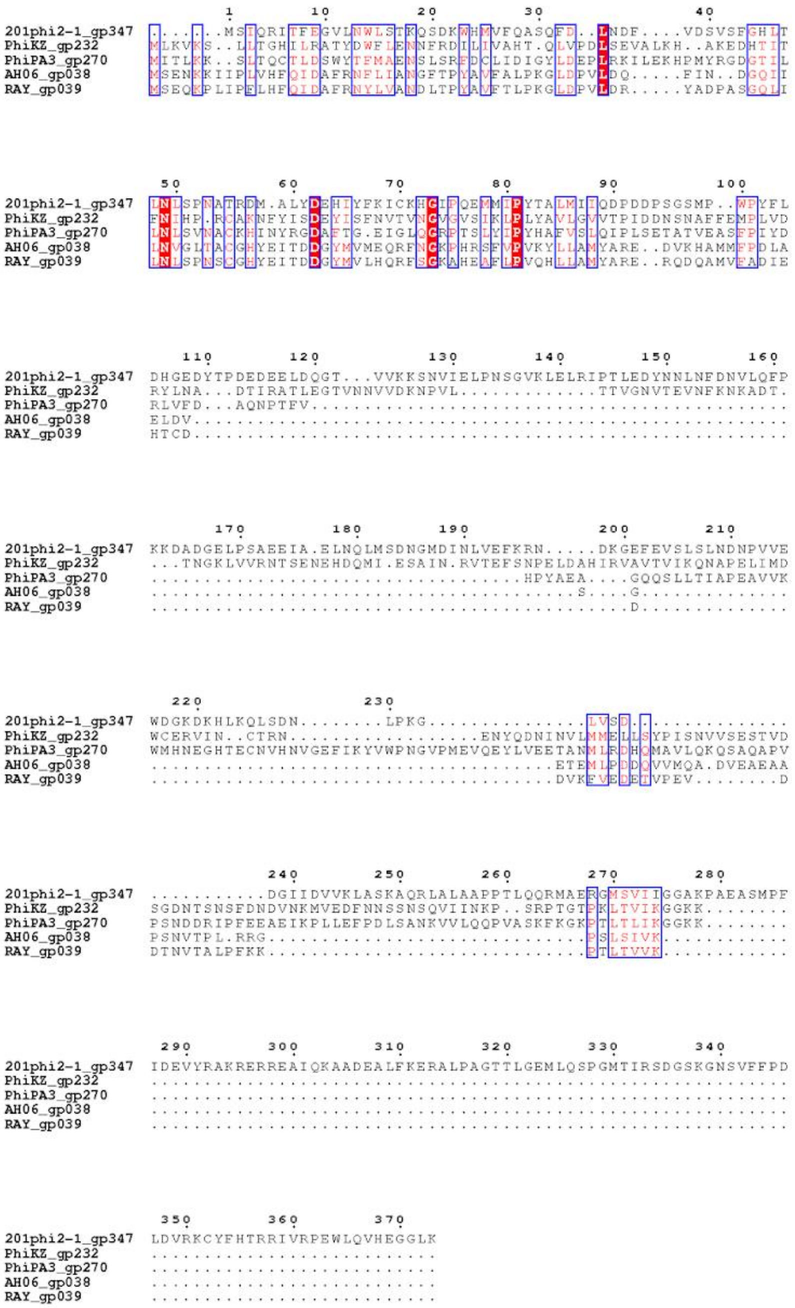

**Figure S12. SspB multiple sequence alignment.** Multiple sequence alignment of RAY gp039 with SspB from previously published nucleus-forming phages and RAY close relative AH06. Since SspB is not part of the core genome, it was not present in some of the previously-studied nucleus-forming phages used for other MSA analysis.

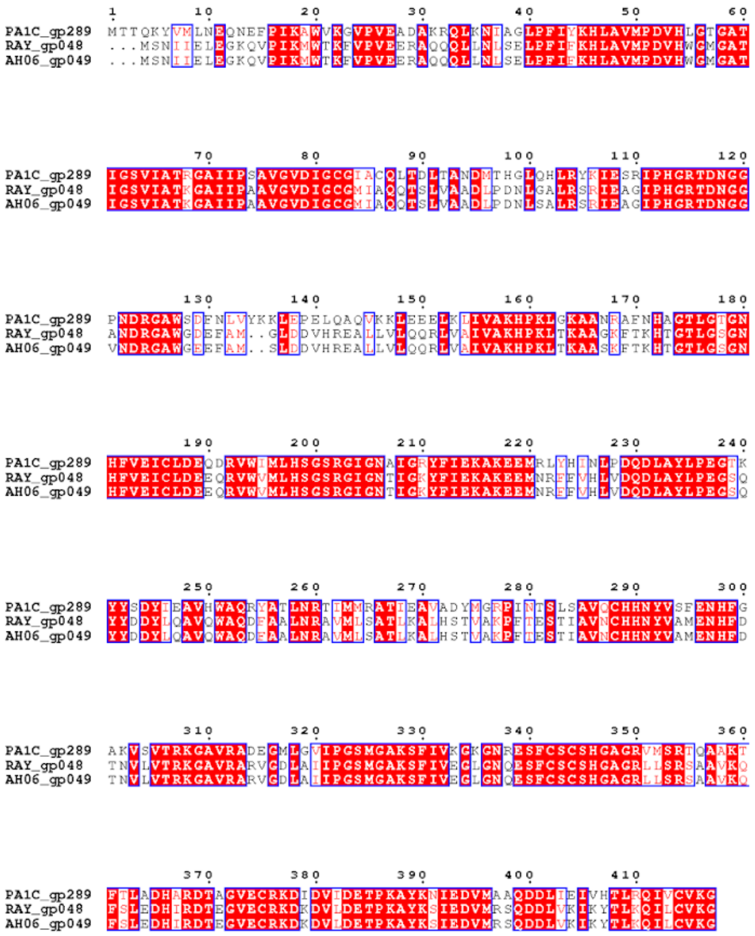

**Figure S13. RtcB multiple sequence alignment.** Multiple sequence alignment of RAY gp048 with RtcB from previously RAY close relative AH06 and ΦPA3 close relative *Pseudomonas* phage PA1C. RtcB is not part of the core genome and was not present in any of the previously-studied nucleus-forming phages used for other MSA analysis. However, despite its uncommon occurrence in Chimalliviridae accessory genomes, its amino acid sequence conservation is remarkably high, even between distantly-related phages like RAY and PA1C.

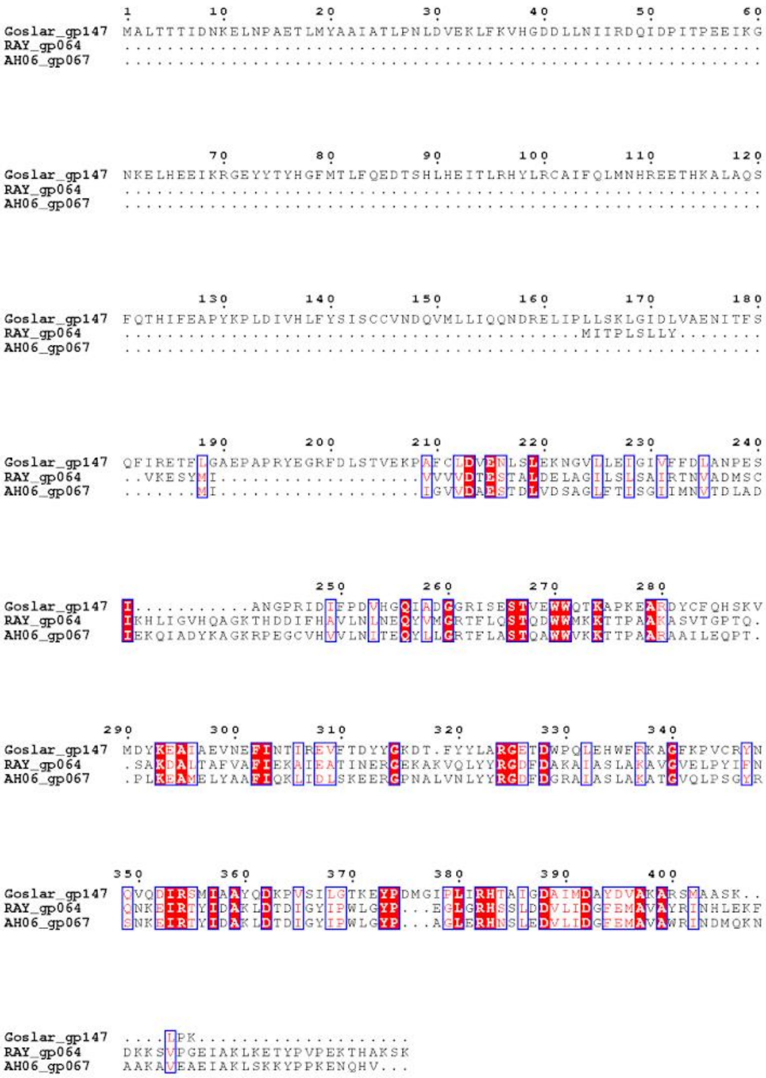


**Figure S14. Exonuclease multiple sequence alignment.** Multiple sequence alignment of RAY gp064 with related exonucleases from previously published nucleus-forming phage Goslar and RAY close relative AH06. Since this exonuclease is not part of the core genome, it was not present in most of the previously-studied nucleus-forming phages used for other MSA analysis.

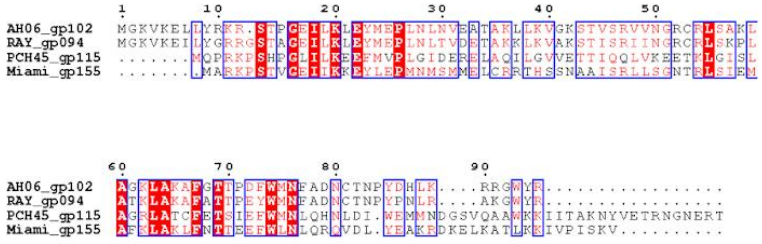

**Figure S15. XRE-like transcriptional regulator multiple sequence alignment.** Multiple sequence alignment of RAY gp094 with XRE superfamily transcriptional repressors from previously published nucleus-forming phage PCH45, RAY close relative AH06, and RAY distant relative Miami. Since this transcriptional regulator is not part of the core genome, it was not present in most of the previously-studied nucleus-forming phages used for other MSA analysis.

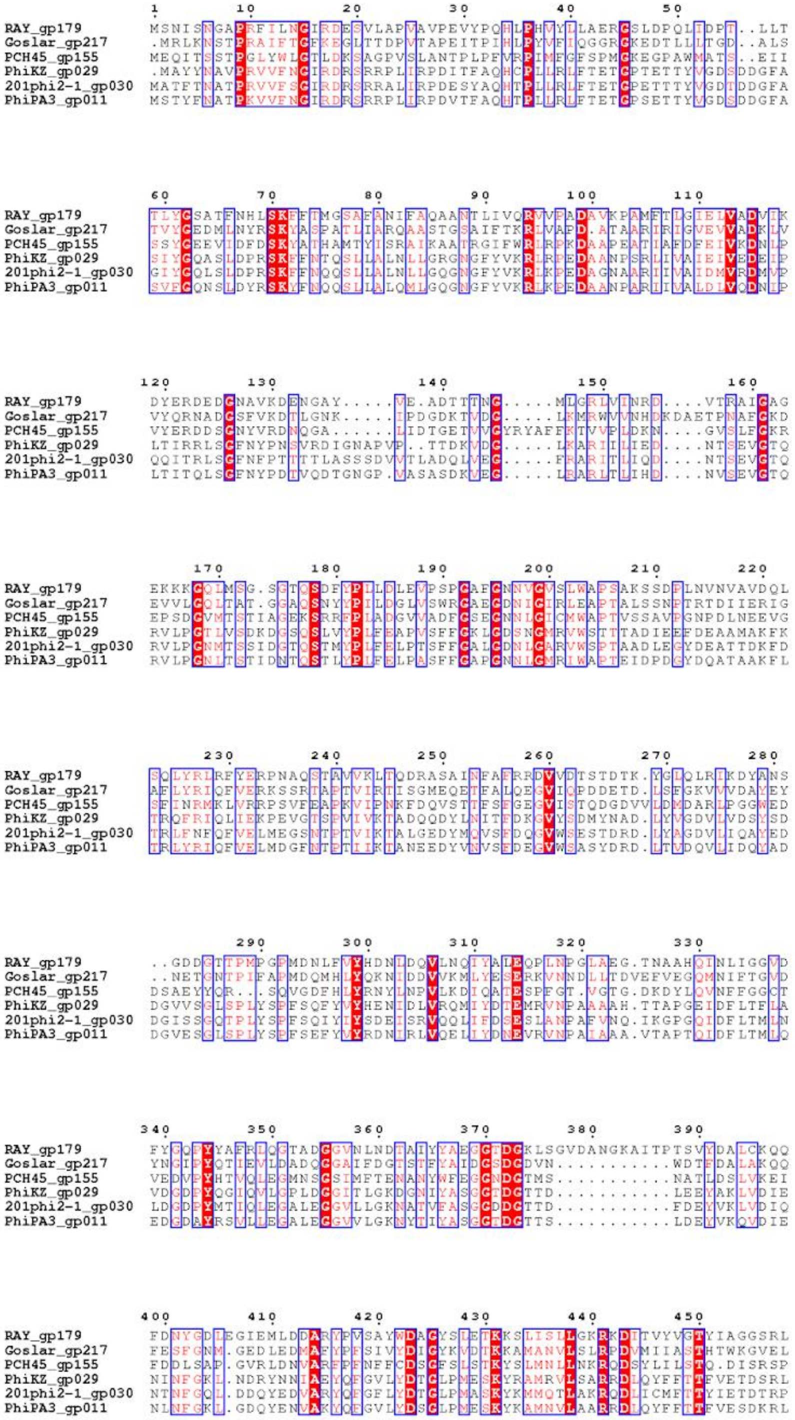
1 10 20 30 40 50
RAY_gp179 MSNISNGAPRFILNGIRDESVLAPVAVPEVYPQHLPHVYLLAERGSLDPQLIDPT..LLT
Goslar_gp217 .MRLKNSTPRAIFTGFKEGLTTDPVTAPEITPIHLPYVFIQGGRGKEDTLLLTGD..ALS
PCH45_gp155 MEQITSSTPGLYWLGTLDKSAGPVSLANTPLPFVRPIMFGFSPMGKEGPAWMATS..EII
PhiKZ_gp029 .MAYYNAVPRVVFNGIRDRSRRPLIRPDITFAQHCPLLRLFTETGPTETTYVGDSDDGFA
201phi2-1_gp030 MATFTNATPRVVFSGIRDRSRRALIRPDESYAQHTPLLRLFTETGPETTTYVGDTDDGFA
PhiPA3_gp011 MSTYFNATPKVVFNGIRDRSRRPLIRPDVTFAQHTPLLRLFTETGPSETTYVGDSDDGFA
60 70 80 90 100 110
RAY_gp179 TLYGSATFNHLSKFFTMGSAFANIFAQAANTLIVQRVVPADAVKPAMFTLGIELVADVIK
Goslar_gp217 TVYGEDMLNYRSKYASPATLIARQAASTGSAIFTKRLVAPD.ATAARIRIGVEVVADKLV
PCH45_gp155 SSYGEEVIDFDSKYATHAMTYISRAIKAATRGIFWRLRPKDAAPEATIAFDFEIVKDNLP
PhiKZ_gp029 SIYGQASLDPRSKFFNTQSLLALNLLGRGNGFYVKRLRPEDAANPSRLIVAIEIVEDEIP
201phi2-1_gp030 GIYGQLSLDPRSKFFNQQSLLALNLLGQGNGFYVKRLKPEDAGNAARIIVAIDMVRDMVP
PhiPA3_gp011 SVFGQNSLDYRSKYFNQQSLLALQMLGQGNGFYVKRLKPEDAANPARIIVALDLVQDNIP
120 130 140 150 160
RAY_gp179 DYERDEDGNAVKDENGAY.....VE.ADTTTNG.....MLGRLVINRD....VTRAIGAG
Goslar_gp217 VYQRNADGSFVKDTLGNK.....IPDGDKTVDG.....LKMRWVVNHDKDAETPNAFGKD
PCH45_gp155 VYERDDSGNYVRDNQGA......LIDTGETVVGYRYAFFKTVVPLDKN....GVSLFGKR
PhiKZ_gp029 LTIRRLSGFNYPNSVRDIGNAPVP..TTDKVDG.....LKARIILIED....NTSEVGTQ
201phi2-1_gp030 QQITRLSGFNFPTTTTLASSSDVVTLADQLVEG.....FRARITLIQD....NTSEVGTQ
PhiPA3_gp011 LTITQLSGFNYPDTVQDTGNGP.VASASDKVEG.....LRARLTLIHD....NVSEVGTQ
170 180 190 200 210 220
RAY_gp179 EKKKGQLMSG.SGTQSDFYPLLDLEVPSPGAFGNNVGVSLWAPSAKSSDPLNVNVAVDQL
Goslar_gp217 EVVLGQLTAT.GGAQSNYYPILDGLVSWRGAEGDNIGIRLEAPTALSSNPTRTDIIERIG
PCH45_gp155 EPSDGVMTSTIAGEKSRRFPLADGVVADFGSEGNNLGICMWAPTVSSAVPGNPDLNEEVG
PhiKZ_gp029 RVLPGTLVSDKDGSQSLVYPLFEAPVSFFGKLGDSNGMRVWSTTTADIEEFDEAAMAKFK
201phi2-1_gp030 RVLPGNMTSSIDGTQSTMYPLFELPTSFFGALGDNLGARVWSPTAADLEGYDEATTDKFD
PhiPA3_gp011 RVLPGNLTSTIDNTQSTLYPLFELPASFFGAPGNNLGMRIWAPTEIDPDGYDQATAAKFL
230 240 250 260 270 280
RAY_gp179 SQLYRLRFYERPNAQSTAVVKLTQDRASAINFAFRRDVVDTSTDTK.YGLQLRIKDYANS
Goslar_gp217 AFLYRIQFVERKSSRTAPTVIRTISGMEQETFALQEGVIQPDDETD.LSFGKVVVDAYEY
PCH45_gp155 SFINRMKLVRRPSVFEAPKVIPNKFDQVSTTFSFGEGVISTQDGDVVLDMDARLPGGWED
PhiKZ_gp029 TRQFRIQLIEKPEVGTSPVIVKTADQQDYLNITFDKGVYSDMYNAD.LYVGDVLVDSYSD
201phi2-1_gp030 TRLFNFQFVELMEGSNTPTVIKTALGEDYMQVSFDQGVWSESTDRD.LYAGDVLIQAYED
PhiPA3_gp011 TRLYRIQFVELMDGFNTPTIIKTANEEDYVNVSFDEGVWSASYDRD.LTVDQVLIDQYAD
290 300 310 320 330
RAY_gp179 ..GDDGTTPMPGPMDNLFVYHDNLDQVLNQIYALEQPLNPGLAEG.TNAAHQINLIGGVD
Goslar_gp217 ..NETGNTPIFAPMDQMHLYQKNIDDVVKMLYESERKVNNDLLTDVEFVEGQMNIFTGVD
PCH45_gp155 DSAEYYQR...SQVGDFHLYRNYLNPVLKDIQATESPFGT.VGTG.DKDYLQVNFFGGCT
PhiKZ_gp029 DGVVSGLSPLYSPFSQFYVYHENIDLVRQMIYDTEMRVNPAAAAH.TTAPGEIDFLTFLA
201phi2-1_gp030 DGISSGQTPLYSPFSQIYIYSDEISRVQQLIFDSESLANPAFVNQ.IKGPGQIDFLTMLN
PhiPA3_gp011 DGVESGLSPLYSPFSEFYVYRDNIRLVQELIYDNEVRVNPAIAAA.VTAPTQIDFLTMLQ
340 350 360 370 380 390
RAY_gp179 FYGQPYYAFRLQGTADGGVNLNDTAIYYAEGGTDGKLSGVDANGKAITPTSVYDALCKQQ
Goslar_gp217 YNGIPYQTIEVLDADQGGAIFDGTSTFYAIDGSDGDVN...........WDTFDALAKQQ
PCH45_gp155 VEDVPYHTVQLEGMNSGSIMFTENANYWFEGGNDGTMS...........NATLDSLVKEI
PhiKZ_gp029 VDGDPYQGIQVLGPLDGGITLGKDGNIYASGGTDGTTD...........LEEYAKLVDIE
201phi2-1_gp030 LDGDPYMTIQLEGALEGGVLLGKNATVFASGGDDGTTD...........FDEYVKLVDIQ
PhiPA3_gp011 EDGDAYRSVLLEGALEGGVVLGKNYTIYASGGTDGTTS...........LDEYVKQVDIE
400 410 420 430 440 450
RAY_gp179 FDNYGDLEGIEMLDDARYPVSAYWDAGYSLETKKSLISLLGKRKDITVYVGTYIAGGSRL
Goslar_gp217 FESFGNM.GEDLEDMAFYPFSIVYDIGYKVDTKKAMANVLSLRPDVMIIASTHTWKGVEL
PCH45_gp155 FDDLSAP.GVRLDNVARFPFNFFCDSGFSLSTKYSLMNLLNKRQDSYLILSTQ.DISRSP
PhiKZ_gp029 NINFGKL.NDRYNNIAEYQFGVLYDTGLPMESKYRAMRVLSARRDLQYFFTTFVETDSRL
201phi2-1_gp030 NTNFGQL.DDQYEDVARYQFGFLYDTGLPMASKYKMMQTLAKRQDLICMFTTYIETDTRP
PhiPA3_gp011 NLNFGKL.GDQYENVAKYQFGVLYDSGLPMESKYKAMNVLAARRDLQYFFTTFVESDKRL

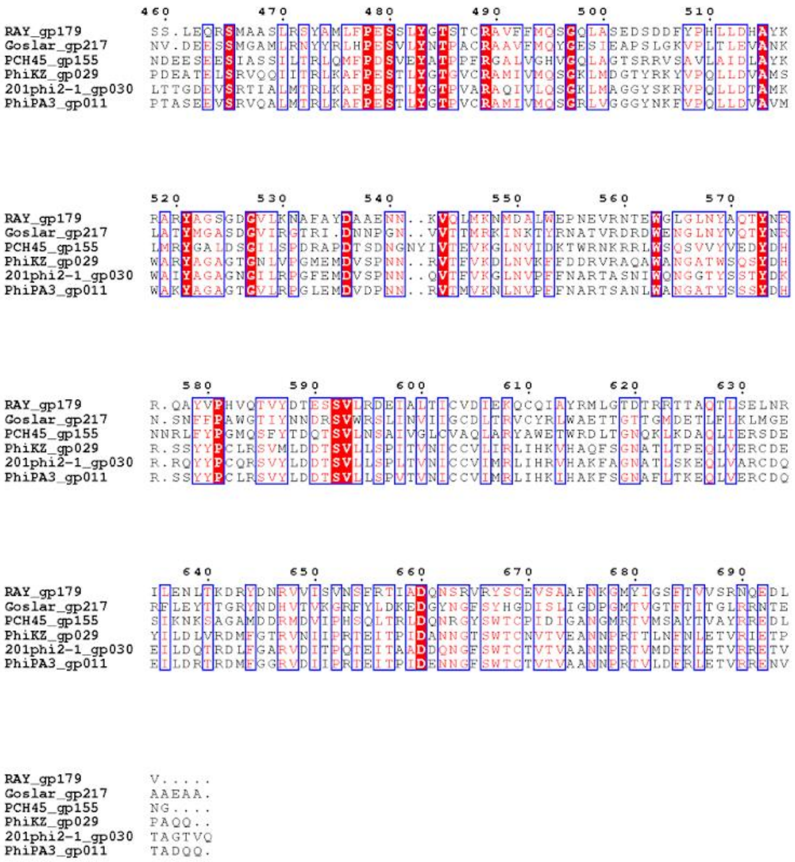

**Figure S16. Tail sheath multiple sequence alignment.** Multiple sequence alignment of RAY gp179 with tail sheath proteins from previously published nucleus-forming phages.

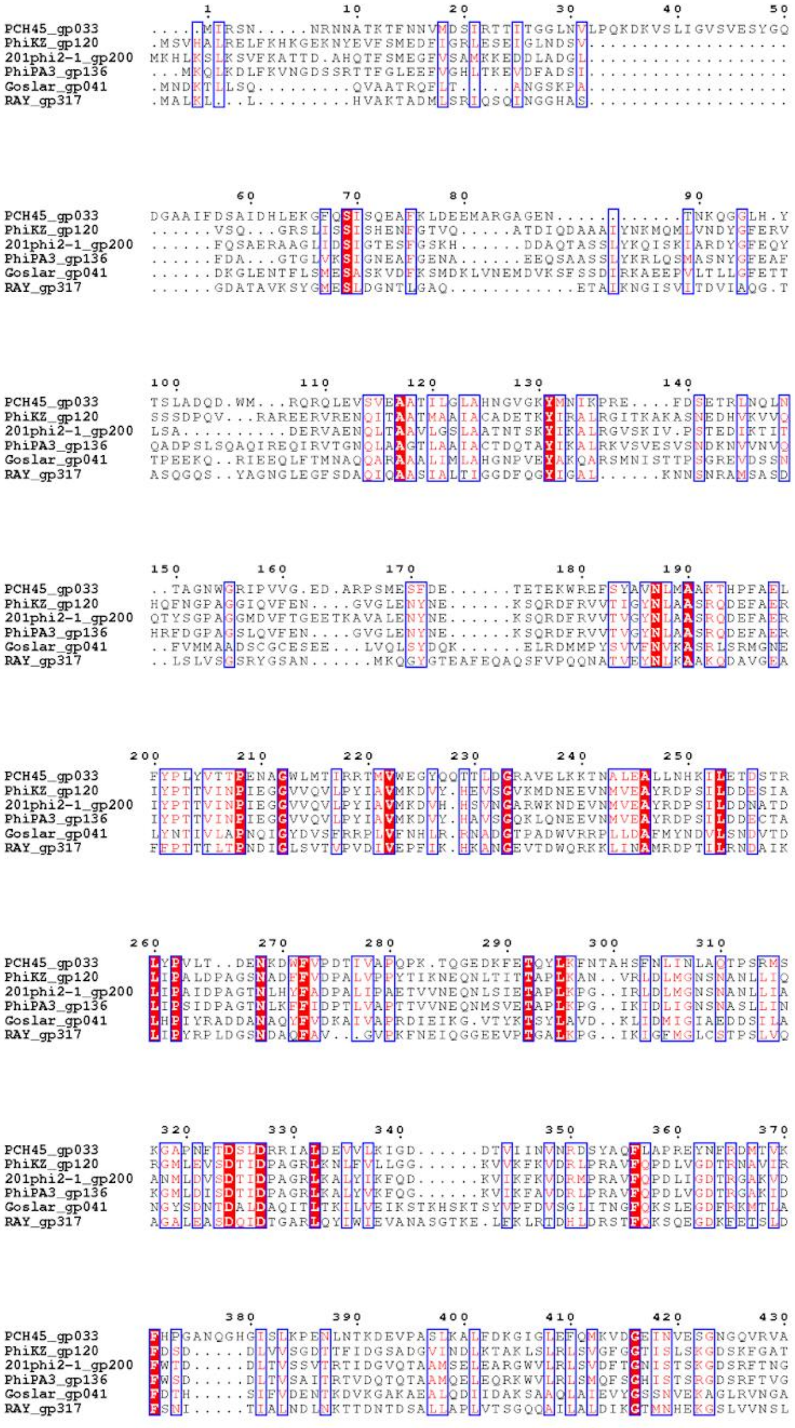
PCH45_gp033
PhiKZ_gp120
201phi2-1_gp200
PhiPA3_gp136
Goslar_gp041
RAY_gp317

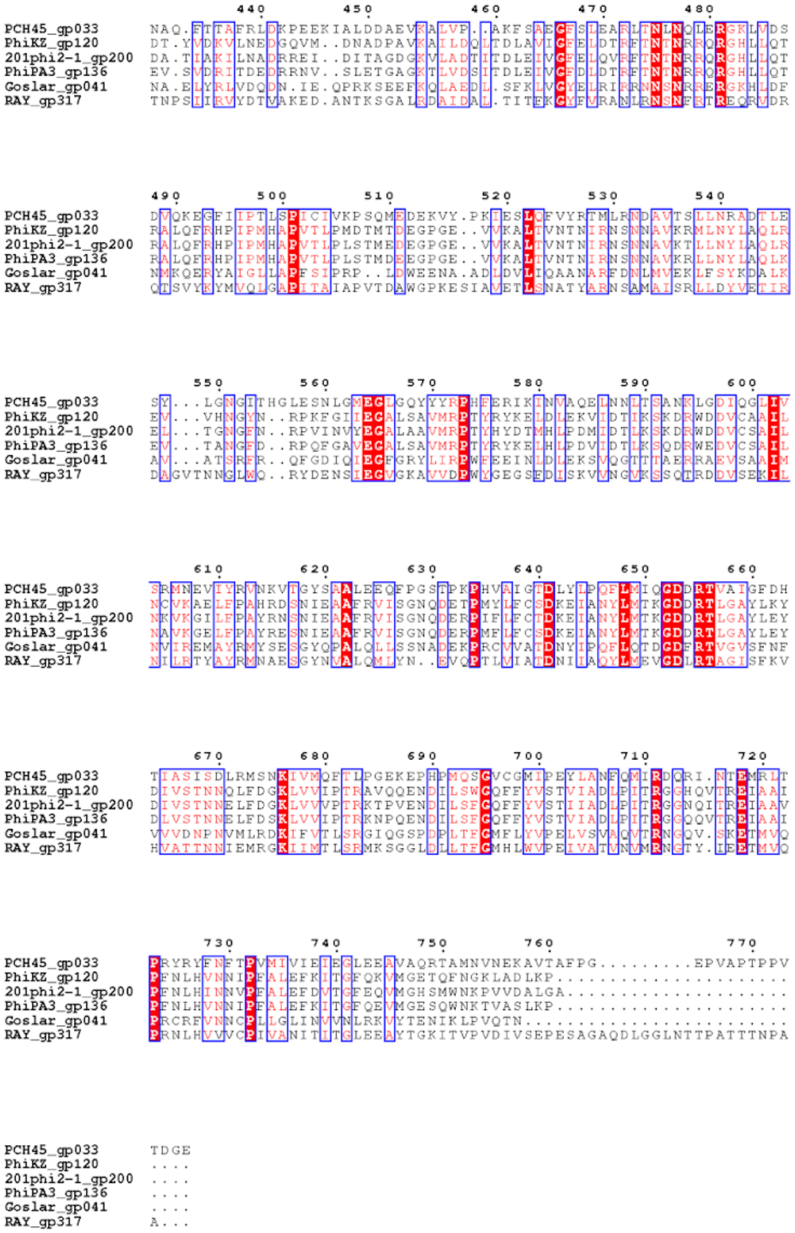


**Figure S17. Major capsid protein (MCP) multiple sequence alignment.** Multiple sequence alignment of RAY gp317 with MCP from previously published nucleus-forming phages.

A

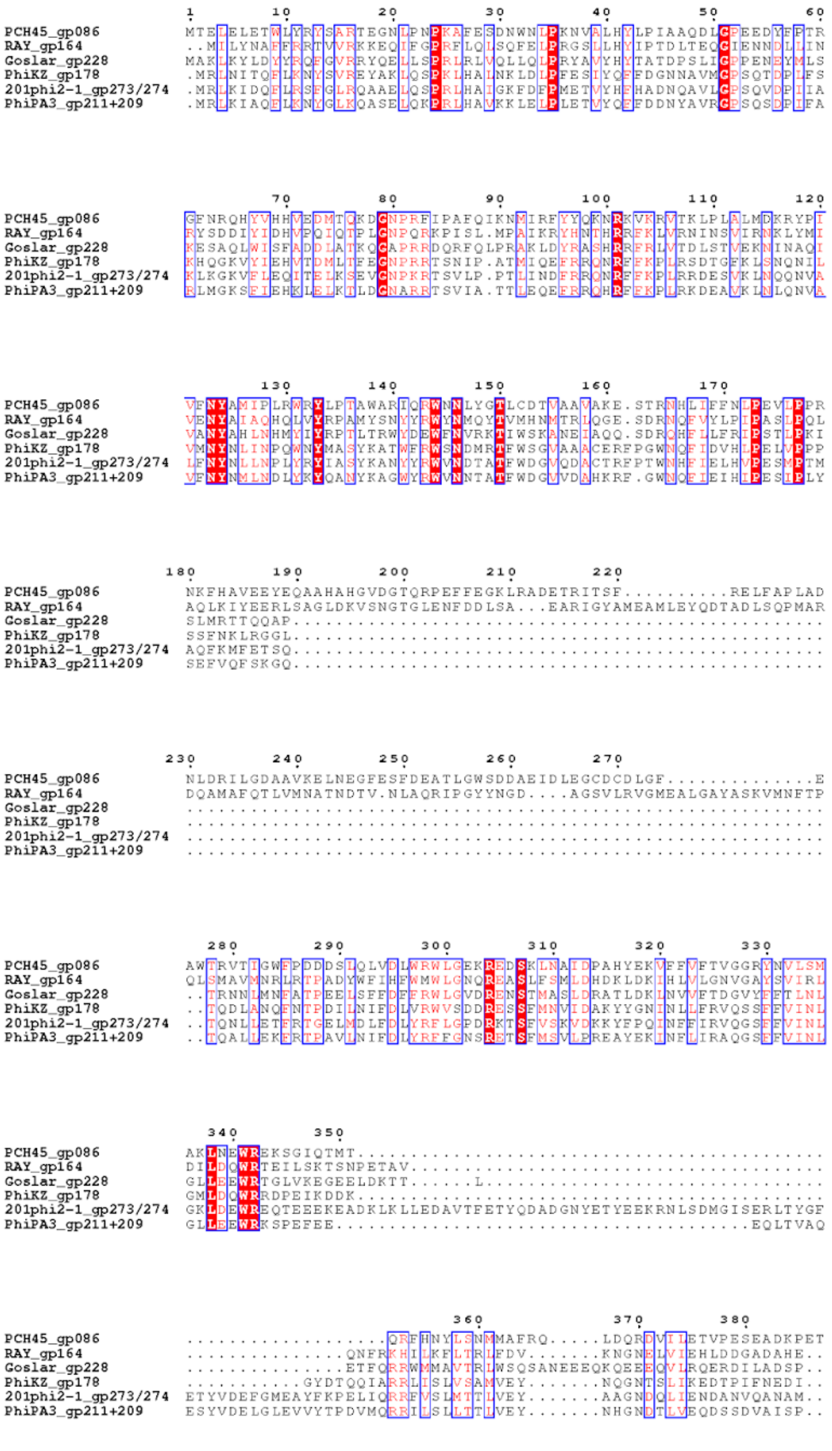
PCH45_gp086
RAY_gp164
Goslar_gp228
PhiKZ_gp178
201phi2-1_gp273/274
PhiPA3_gp211+209

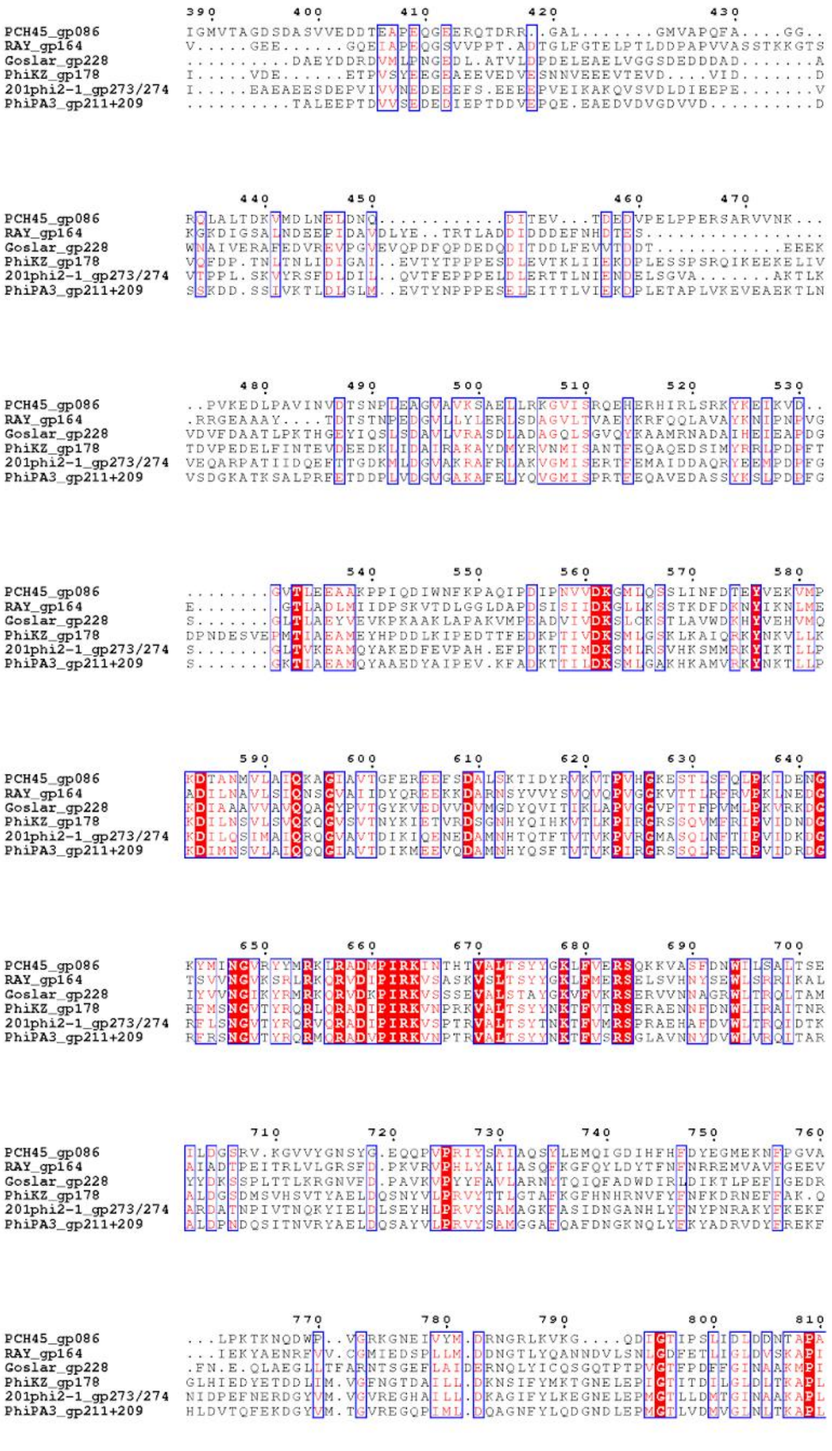
PCH45_gp086
RAY_gp164
Goslar_gp228
PhiKZ_gp178
201phi2-1_gp273/274
PhiPA3_gp211+209

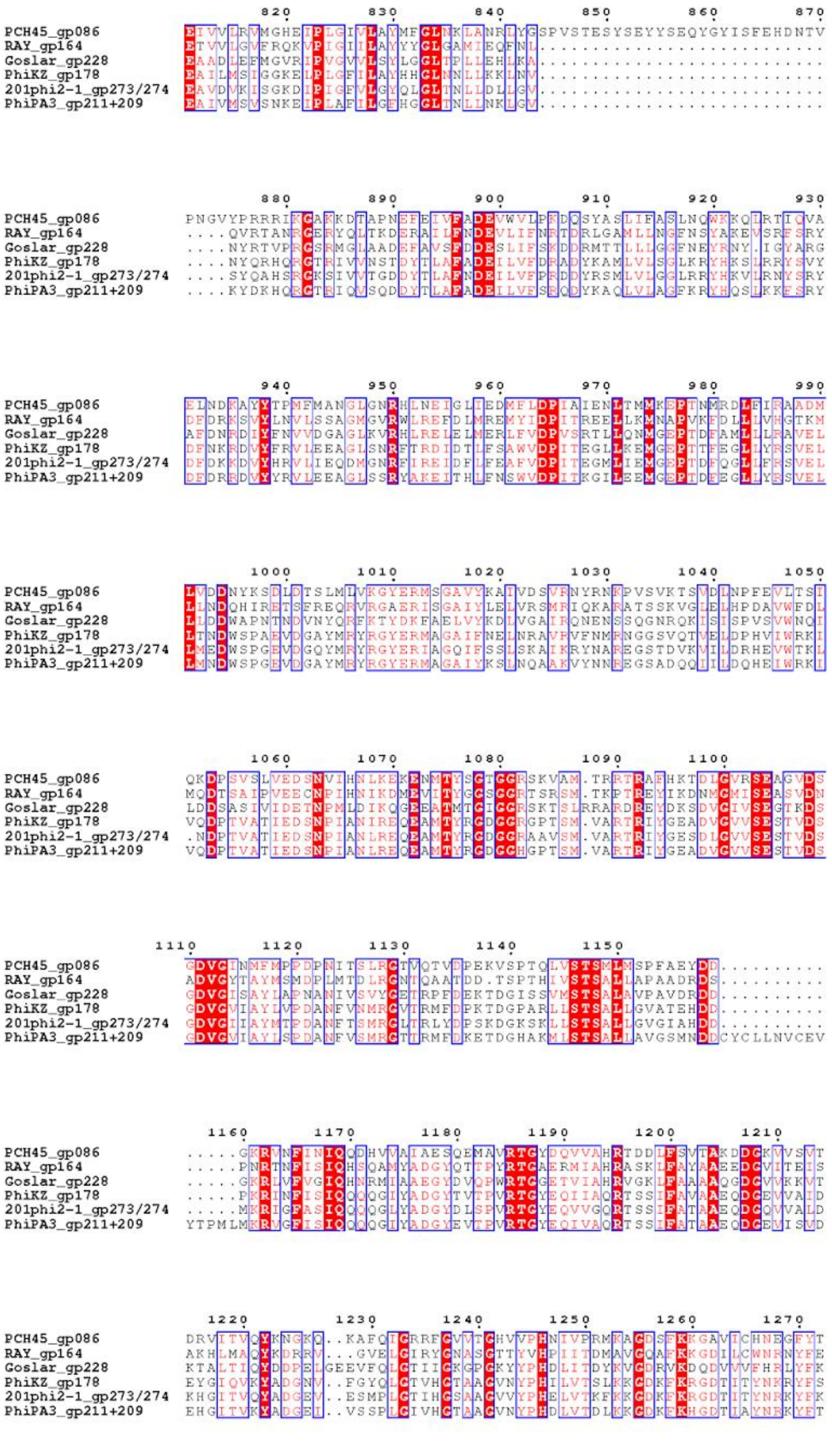
PCH45_gp086
RAY_gp164
Goslar_gp228
PhiKZ_gp178
201phi2-1_gp273/274
PhiPA3_gp211+209

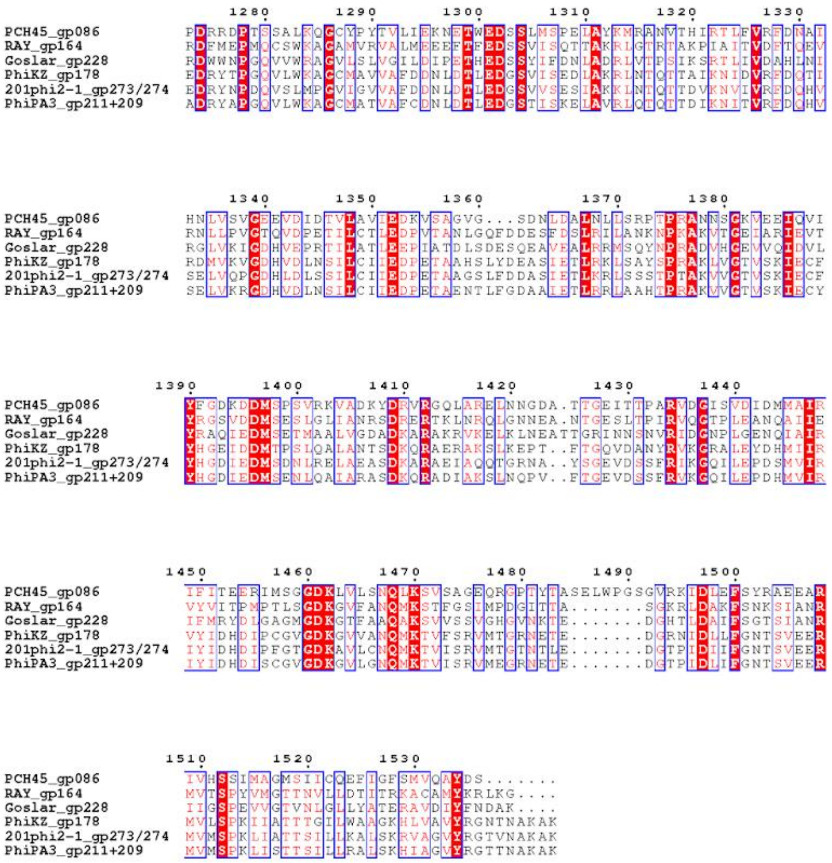

PCH45_gp086
RAY_gp164
Goslar_gp228
PhiKZ_gp178
201phi2-1_gp273/274
PhiPA3_gp211+209

B

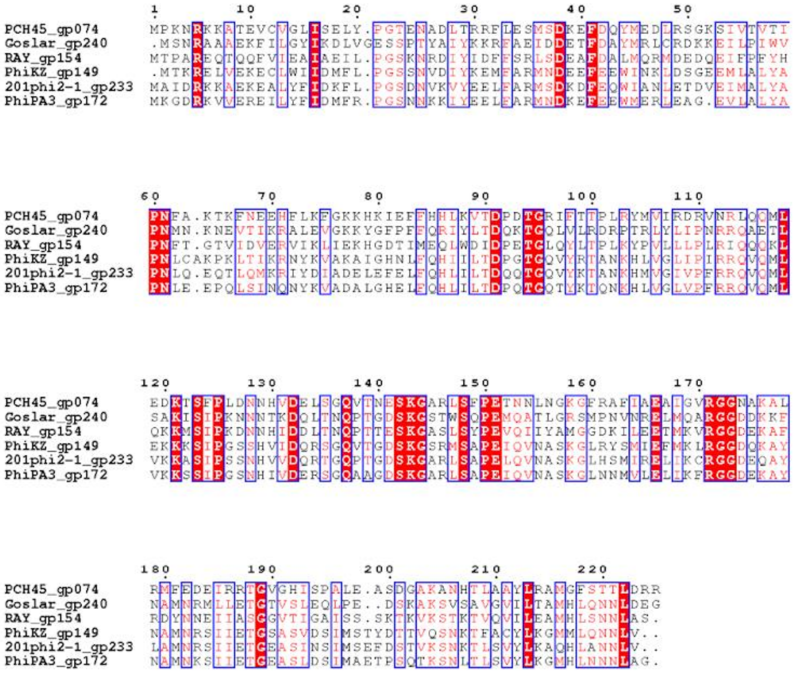
PCH45_gp074
Goslar_gp240
RAY_gp154
PhiKZ_gp149
201phi2-1_gp233
PhiPA3_gp172

C

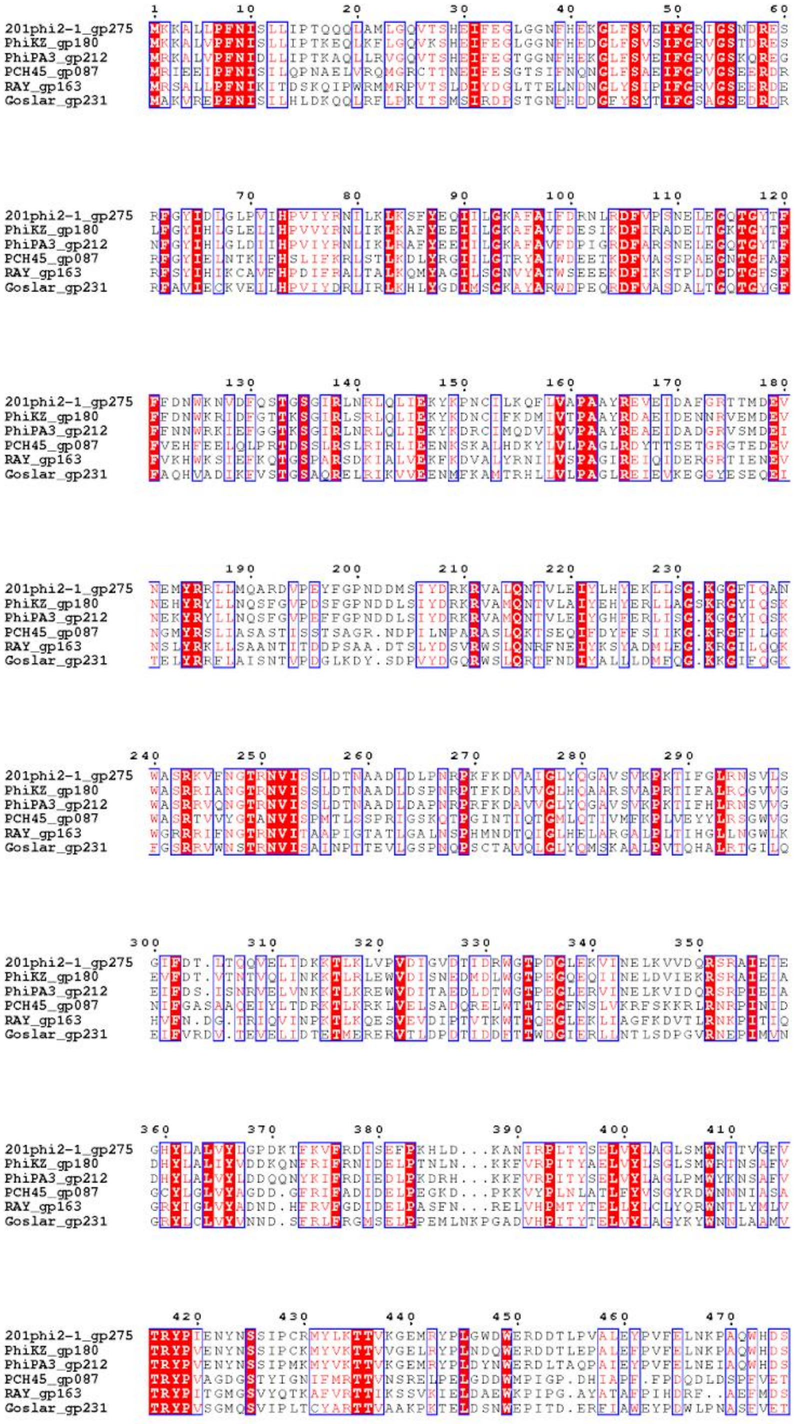
201phi2-1_gp275
PhiKZ_gp180
PhiPA3_gp212
PCH45_gp087
RAY_gp163
Goslar_gp231

```
                       480        490        500        510        520        530
201phi2-1_gp275  ISLPPATLAPLTADFDGDTT.SWIACYSEESIAECNKFFKSRLAYIRAGGGIAFSLNIHT
PhiKZ_gp180      VSISPSILAPLGAD..............................................
PhiPA3_gp212     TSVSPAVLAPLTADFDGDTV.SFNAVYSKEAIEESDKFFKSRLAYIKAGGGLAFSINIHT
PCH45_gp087      MSPHTARIKAQGGDFDGDRESC.IGAMSKEARDEYQRLKSKRDWYVGADNKLVLSAANDN
RAY_gp163        LSVHPSRLGLLGGDYDGD.TMSATILLGDDVLRECAMLLSRRESYITGRGEFLIDVTNET
Goslar_gp231     TIVPAPYLAGLGGDQGKANKMPVFWQI.................................

                    540        550
201phi2-1_gp275  LNLTLRYMTGDPIPRK
PhiKZ_gp180      ................
PhiPA3_gp212     LNLTLRYMTGDPVARD
PCH45_gp087      IELLLLQITGGPTK..
RAY_gp163        LDFTLKALTS......
Goslar_gp231     ................
```

D

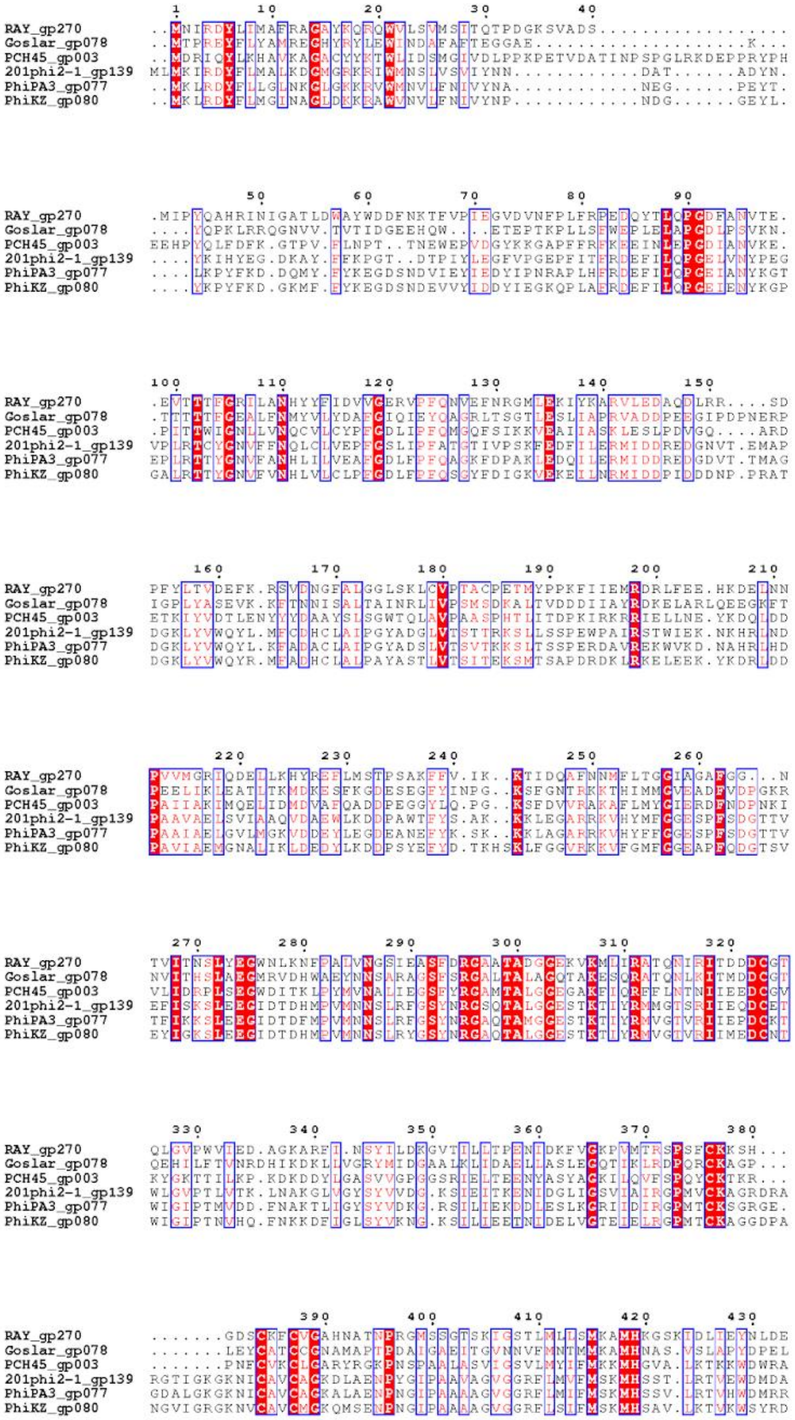
RAY_gp270
Goslar_gp078
PCH45_gp003
201phi2-1_gp139
PhiPA3_gp077
PhiKZ_gp080

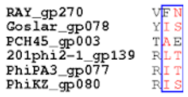


**Figure S18. Virion RNA polymerase multiple sequence alignment.** Multiple sequence alignments of RAY gp164 (A), gp154 (B), gp163 (C), and gp270 (D) with vRNAP subunits from previously published nucleus-forming phages. These RAY proteins are homologs of known phage msRNAP subunits ΦKZ gp178, gp149, gp180, and gp80, respectively (Ceyssens et al., 2014).

```
                   1        10        20        30        40        50
Miami_gp214        MLN....YIFGPDVKQG.PTITEINHVNFRNFNDDIQTAYGTSTPGRYMFRRVGSNKFVT
Goslar_gp160       MLDRINRLMNRITVKNQSNNSVHIGGIDAWRLSDDIQRIWGTSRFMKHMFRNFSSSGLSL
RAY_gp131          MFDYLRRALGFINVEEK.NDIITITGFNAPLATRDILKVWKTSKIAGYLFREVTQNKISF
AH06_gp136         MFDYLRRALGFINVEEK.NDVITITGFNAPLATRDILKVWKTSKVAGYLFREVTQNKISF
PhiKZ_gp203        MIDSFRKLMGSLSITET.DQEIIISGFDGAAFIRDINKYWRTTKLATQLFNTVSRRSISF
201phi2-1_gp300    MLDQFRNVFGGVTVKET.NTEIVVSGIRAKDIVRDMDKHWKTTRITQNIFNTVSGNSFSF
PhiPA3_gp233       MFDTFRTLFSGLDVKET.DTEIIISGVAANEIIRDMDKFWKTTKLTGNMFNNVSRRSMSF
```

```
                     60        70        80        90       100       110
Miami_gp214        DNFFLLELDHLITKILQSRNTWSSRKNLNELQIKLRTDTWIKDTVTPTAYPIDK...ASF
Goslar_gp160       HNFYLLDFVYILETIVEAKNTRSNKRMLKHLIEVLLQETWLQNTTIEQPAILDK...ARM
RAY_gp131          NSFFAIEVEYILKQLYEHDKTWSDRRGLGKILELLRKNTWMRNLEEQ....YEDIIDLGQ
AH06_gp136         NSFFAIEVEYIFKQLLDHDKTWTDRRGIGKILDLLRKNTWMRNLDEQ....YEDIIDLSQ
PhiKZ_gp203        YKFFAPEIYYMLEAVKNYRSRYISIKTVNAIREAMLQYTWLKNTRPVDTNSVPGRLNFKM
201phi2-1_gp300    YKFFAPEIAYILDGIKKYRHRWTSIRAINGIIEGLNEHTWLKGTIPPDTTTIKGRLDFRK
PhiPA3_gp233       YKFFAPEVMYMLENIKHYRNRWTSIKAINAIREALMEHTWLKGTIPVDPNTVPGRLDLRK
```

```
                        120       130       140       150       160
Miami_gp214        LKQFKIKPYDTQWGFLSTYKMVKKSHHLKGLLLDAKAGSGKTFTSLMWTRL.........
Goslar_gp160       NKMLTLSLFDYQDEFIDIYNRNVPKYRLKGYVCAMGAGAGKTITALA.............
RAY_gp131          LKYVKKTPLPEQRNWLNYYNTVIPRYGLRGALLSAAPGAGKTLCSIM.............
AH06_gp136         LKLVKKSPLPEQRNWLNYYNTAVPRYSLRGALLSAAPGAGKTLCSIM.............
PhiKZ_gp203        LDKLTFTPDESQKAYFENYNYRIDQYGLRGDIVAGKPGTGKTF.................
201phi2-1_gp300    LKNLHFTPKPYQMEYFKNYSYRLDQYNLKGDLIAAAAGTGKTYSM...............
PhiPA3_gp233       LQNLHFDAMTYQMEYFQNYSYRLDQYGLKGDLIAAAAGTGKAQPLTSMVKVPGGWKAMGN
```

```
Miami_gp214        ............................................................
Goslar_gp160       ............................................................
RAY_gp131          ............................................................
AH06_gp136         ............................................................
PhiKZ_gp203        ............................................................
201phi2-1_gp300    ............................................................
PhiPA3_gp233       IQVGDVVTAWDGTPTKVTGVYPQGKKQTFTVTFKDGRTTKACDEHLWNVYCQDWTRYGGT
```

```
Miami_gp214        ............................................................
Goslar_gp160       ............................................................
RAY_gp131          ............................................................
AH06_gp136         ............................................................
PhiKZ_gp203        ............................................................
201phi2-1_gp300    ............................................................
PhiPA3_gp233       GWRVINTLELFGRIQSGKQRLYVQLCKSEEGIDVELPIDPYNLGVILGDGCISSNCVSVT
```

```
Miami_gp214        ............................................................
Goslar_gp160       ............................................................
RAY_gp131          ............................................................
AH06_gp136         ............................................................
PhiKZ_gp203        ............................................................
201phi2-1_gp300    ............................................................
PhiPA3_gp233       SGDPQLFTEFAKALPENLELITRDDITMGVINKKGERNPYTSALREMGLLGENSLTKFIP
```

```
Miami_gp214        ............................................................
Goslar_gp160       ............................................................
RAY_gp131          ............................................................
AH06_gp136         ............................................................
PhiKZ_gp203        ............................................................
201phi2-1_gp300    ............................................................
PhiPA3_gp233       QNYLMASTAQRLALVQGLMDTDGTVDVNSSLSFSTSSYMLAKQFQYLIRSLGGIAKISFK
```

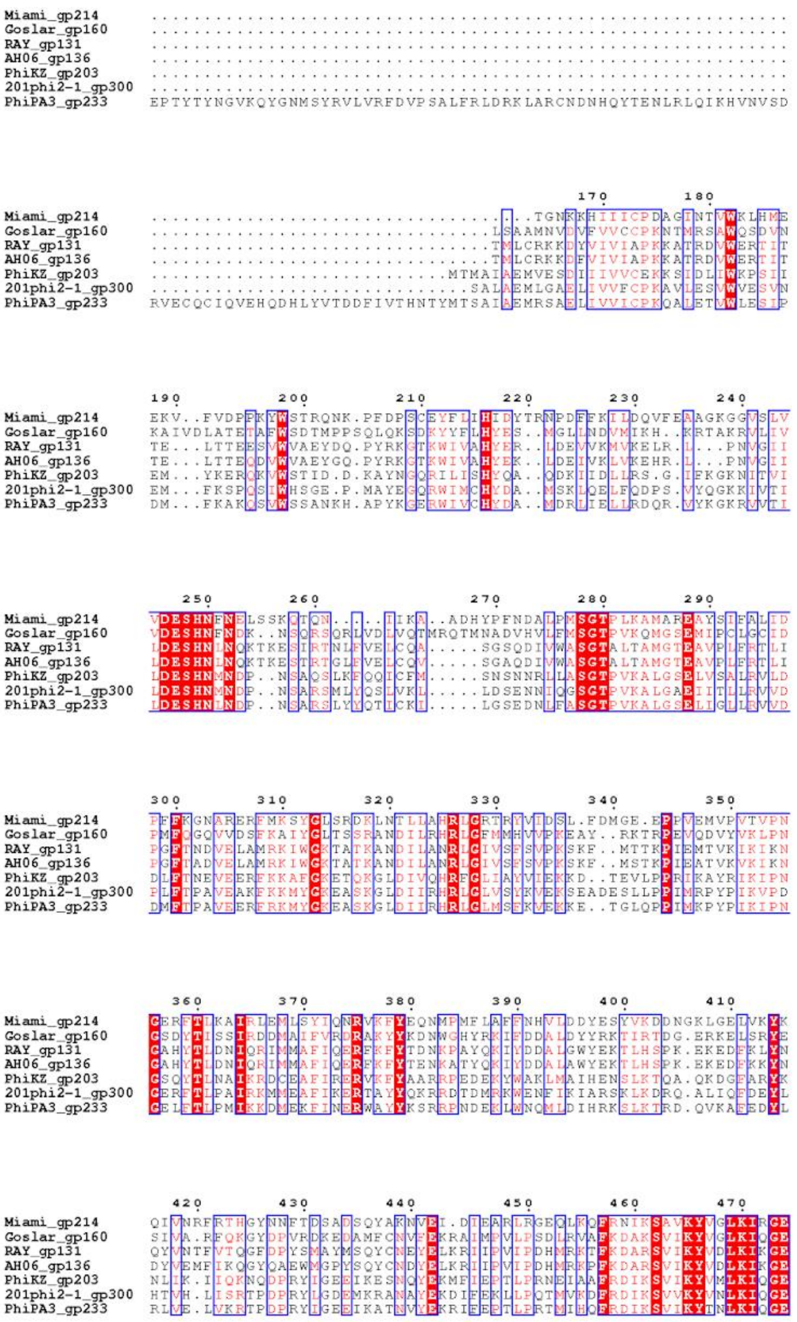
Miami_gp214
Goslar_gp160
RAY_gp131
AH06_gp136
PhiKZ_gp203
201phi2-1_gp300
PhiPA3_gp233
170
180
190
200
210
220
230
240
250
260
270
280
290
300
310
320
330
340
350
360
370
380
390
400
410
420
430
440
450
460
470

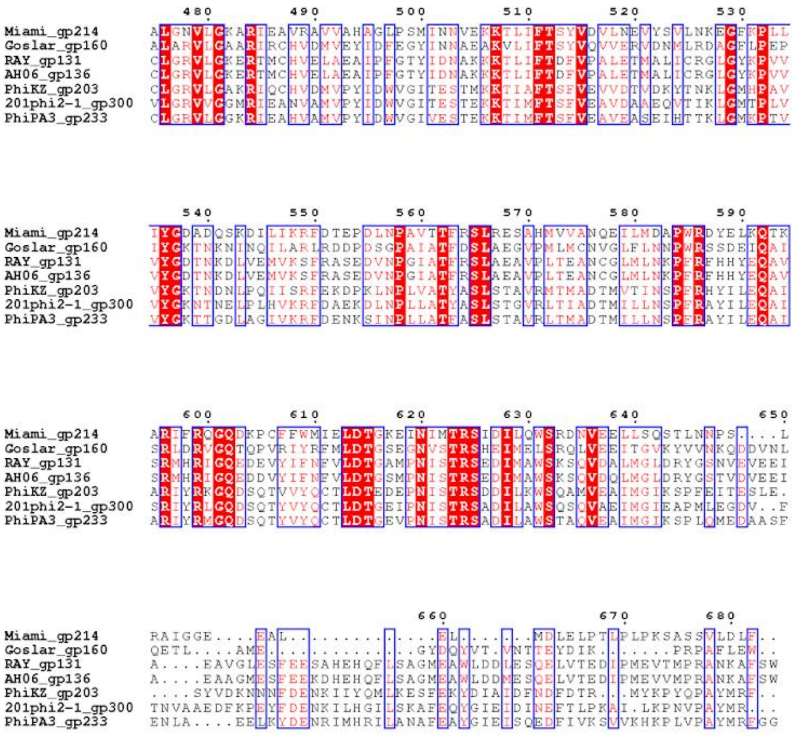

**Figure S19. Virion SF2 helicase multiple sequence alignment.** Multiple sequence alignment of RAY gp131 with SF2 helicase homologs from previously published nucleus-forming phages, as well as RAY close relative AH06 and RAY distant relative Miami. RAY gp131 localized similarly to capsids (Fig. 5B) and was seen in virion mass spec experiments (Thomas et al., 2012) as opposed to RAY gp250, another SF2 helicase homolog which localized in the phage nucleus (Fig. 4B), leading us to call gp131 and its homologs the "virion" SF2 helicase. While this virion SF2 helicase is not part of the core genome (and not found in PCH45), it is widely conserved in a majority of Chimalliviridae.

```
                1         10        20        30        40        50        60
AH06_gp315    MTTMSELGPAHTTLLAEEKDVQKDAATDATETALAVLKKTSKSKPEKDDEDEPTISFGAE
RAY_gp299     MSTMTELGPSQGTLLADEKAVHDDAATEATEVALHVLKKIGKKKEKDDEDEEVVLSFGSE
Miami_gp025   ....................................................MSTPYTQK

              70 80 90 100 110
AH06_gp315    GYTMLTSMSFDLTE.NTDLA......IQELLDKISKIVIEANGNLTDKDLEAIKN.VGNE
RAY_gp299     GYAMLTSMSFDLTE.NSDLA......VQELLDKITTIVIEANGNLTQKDLEKIKS.VGNE
Miami_gp025   GMSMSLAEEYEGDKKLDRYIEGTPDNVERYKKVAAVIGKEPGHVTDDK.DTIDNTAGTE

              120 130 140 150
AH06_gp315    .............ALLDAGNEDYKEVMNLVMRKIMQFIRWAINQGKEGIKRISDRLSRLS
RAY_gp299     .............SLLGDGNEDYKEVVTMVMKRIMQFIRWAINQGKEGIKRVADRLSRLS
Miami_gp025   EFDIDEAVLNFPEDENVAGMEDFVEFAKKAIVRLMRMIKNAAIFIYELVFNKVSRIKHIA

              160 170 180 190 200 210
AH06_gp315    VKAMYVERKIDVSADNSLPNDKFVLPRTYPLIMMANKPPANAMEVINVLNRTKYLFTTLH
RAY_gp299     VKTMYVERKIDISTDNSLPTDKFVLPRSYPMLMMVNKPPANAMEVINAVNRTKYLFTTLH
Miami_gp025   RKAEYQLRT.....QSLIPT.LVPYPKGTRKIFIGPALQTNANWTLKSIQQVQLFYTRLM

              220 230 240 250 260 270
AH06_gp315    NDYQNFQNLFKSAVATGSRADTLEMINSYLTSLASKLSARPNPLFDNRLSF.........
RAY_gp299     NDYQGFQALFKAAVATGSRADTLEMINNYLNGLISKLSARPNPLFDNRNTF.........
Miami_gp025   KAHGN...LLDAA.KSG..................NWRNNPNFIFETAATFANEICQNHH

              280 290 300 310 320
AH06_gp315    ......NQLPGG..YRLVFSQGQAFADCSATLTRTSERYDMAPAVQRPDKASLMRLVAEV
RAY_gp299     ......NQLPGG..YRLVFSEGQSFADCGATILRTPEKYEAAPAVQRPDKASLVRLVSEI
Miami_gp025   SDYSETQVLPGGKIYRVNYSKGSGKNRPGVSLSQKL.SVQLKDKSFKPDRAIADDLMTAL

              330 340 350 360 370
AH06_gp315    KTYLRIINEVY...GRVSGRLEADFRAIVKSAEREVKS.FDSTADIRTASTTIEWFTEQQ
RAY_gp299     KTYLRVINEVY...GKVSSRLEVDFRNIVKAAEKEVKG.FDSTADIRTASTTIEWFTEQQ
Miami_gp025   QGFADKIGNDHRSQTSTSRKMEAAIR........DLERYYVDPEAKRHAAAYFTWLIDFQ

              380 390 400 410 420 430
AH06_gp315    SKLYTRSMMLSCTVLNASLDYCLSAIGAKPAAGTESFDITETSSISYAIESLDEQLERLD
RAY_gp299     SRLYTRSMMLSCTVLNAALDYCLGAIGAKPAVGTESFDILDTSSMGYAIESLGEQMERLD
Miami_gp025   R...........TVLSTGLQYVFGTLFGYAEFISANVK......................

              440 450 460 470 480 490
AH06_gp315    AGLCSLEIDSRSMQSIADVEELVDVDNDQVINLLMNQTPASYYNPLYGISFAGLQGVLEE
RAY_gp299     AGLCELEIDARTMQSITDVKEMVDVDNDHVIQLLMQQRPSSYFNPPDGLSEYSLGDAFNG
Miami_gp025   ............................................................

              500 510 520 530 540 550
AH06_gp315    KKGAEYIASRLGGIVDITKQLDNTAEVIKQIMTDLPDGEVGAVKPGLK......PTVADV
RAY_gp299     SATARYIGRRLTSIVNLTNQLSGTTDLMKGLLAEMPTGEIK...PGSDADVIERNSLLDR
Miami_gp025   ............................................................

              560 570 580 590 600 610
AH06_gp315    EFTSDHPWCAFLHRVDRQSIYASDVVRYVGDTYEKLRGVNNTLSVLAEQMTKTVVDGEMN
RAY_gp299     EMFEGHPLCAFLHRVDREGIYATDVIRYVTDTYEKLRQVRTTLAVQAEQMTLTVADGEMN
Miami_gp025   ............................................................
```

```
                        620       630       640       650       660       670
                          .         .         .         .         .         .
AH06_gp315     VSLLKEYMFNAPTEPFTASSLCGGFTISPVAEKLGKLVVRGYVLESPPPLKPMVTIHRPS
RAY_gp299      VSLLKEFLFNAPQEPFTDSVLCGGFRLQPVAEKLGELVIHGAVMDSPPPLKPMTTIHRPS
Miami_gp025    ............................................................

                        680       690       700       710       720       730
                          .         .         .         .         .         .
AH06_gp315     YDEYNEINSLAARFNAEQARLAEITRLLQIGTGHLRYITASVADNLTADGLKGGGDGWFS
RAY_gp299      YDEYNGVNQLASLFGNELRELAKIAERLQIGTGYLRYITTSVADNLNEDGLKGGGDGWFN
Miami_gp025    ............................................................

                        740       750       760       770       780
                          .         .         .         .         .
AH06_gp315     TALEYLAISARQYRWMYRLTVQLAIYQRTMISALRQYNNEGGWDGDKQ
RAY_gp299      TALEYLAVSARQYRWMYRLTVQLAVYEHTMINALRHYNNSGGWHD...
Miami_gp025    ................................................
```

**Figure S20. Capsid-associated protein multiple sequence alignment.** Multiple sequence alignment of RAY gp299 with homologs from RAY's close relative AH06 and its distant relative Miami. This protein is found in capsids (Thomas et al., 2012) but is not part of the core genome and was not present in any of the previously-studied nucleus-forming phages used for other MSA analysis. As there is very little information about this protein, we are not able to make any hypotheses concerning its potential function, and it remains an area of future study.

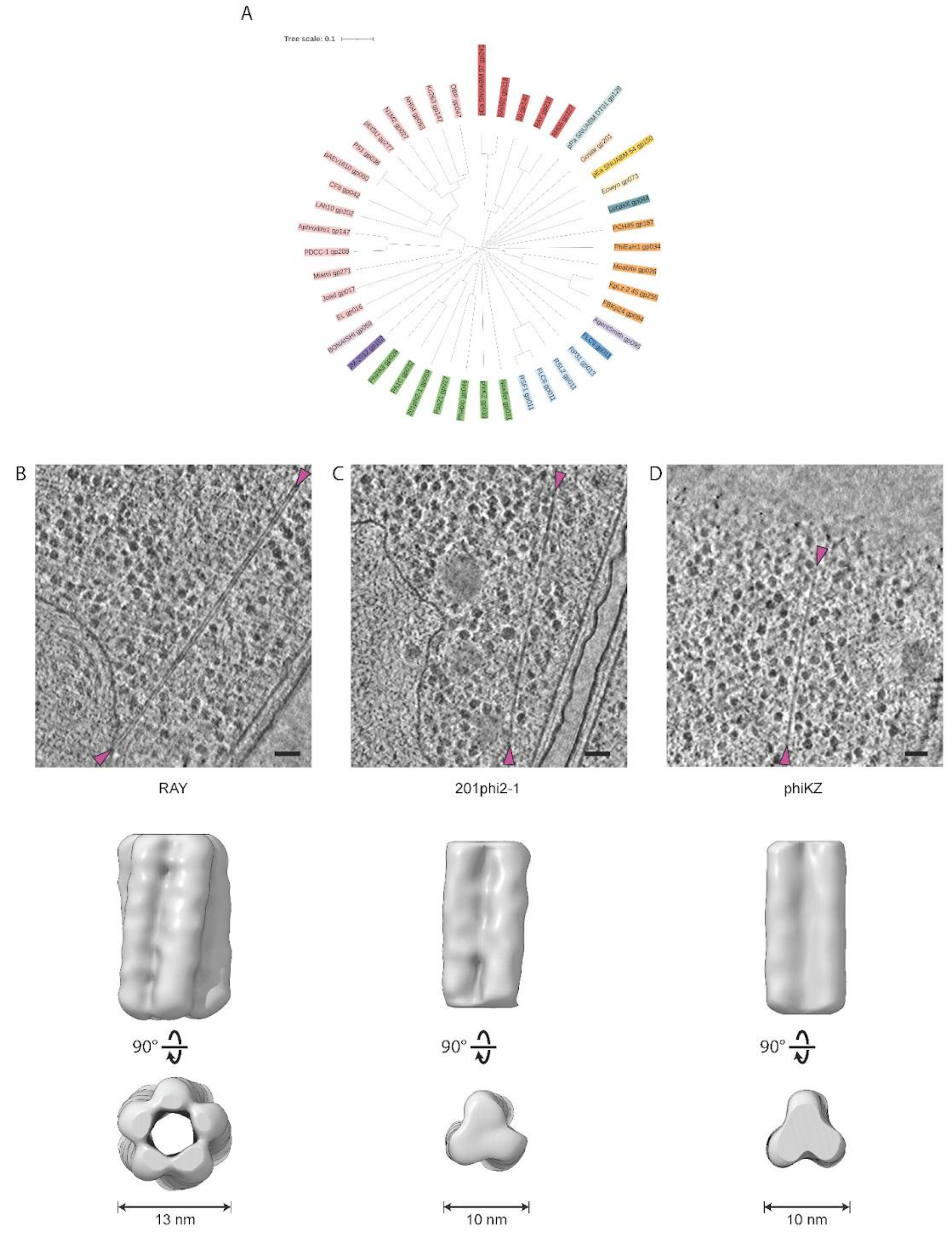


**Figure S21. PhuZ protein phylogeny and filament structural comparison.** (A) A protein tree of PhuZ homologs from Chimalliviridae. While PhuZ is not a member of the core genome, it is present in a majority of Chimalliviridae. Chimalliviridae are color-coded by predicted genus as in Figure 1. (B,C,D) Top, tomographic slices from jumbo phage infected host cells with the ends of putative PhuZ filaments marked with magenta arrows. Bottom, orthogonal views of subtomogram averages from the corresponding tomographic datasets for (B) RAY, (C) 201ϕ2-1 (EMPIAR-10859), and (D) ΦKZ. Scale bars: 50 nm.

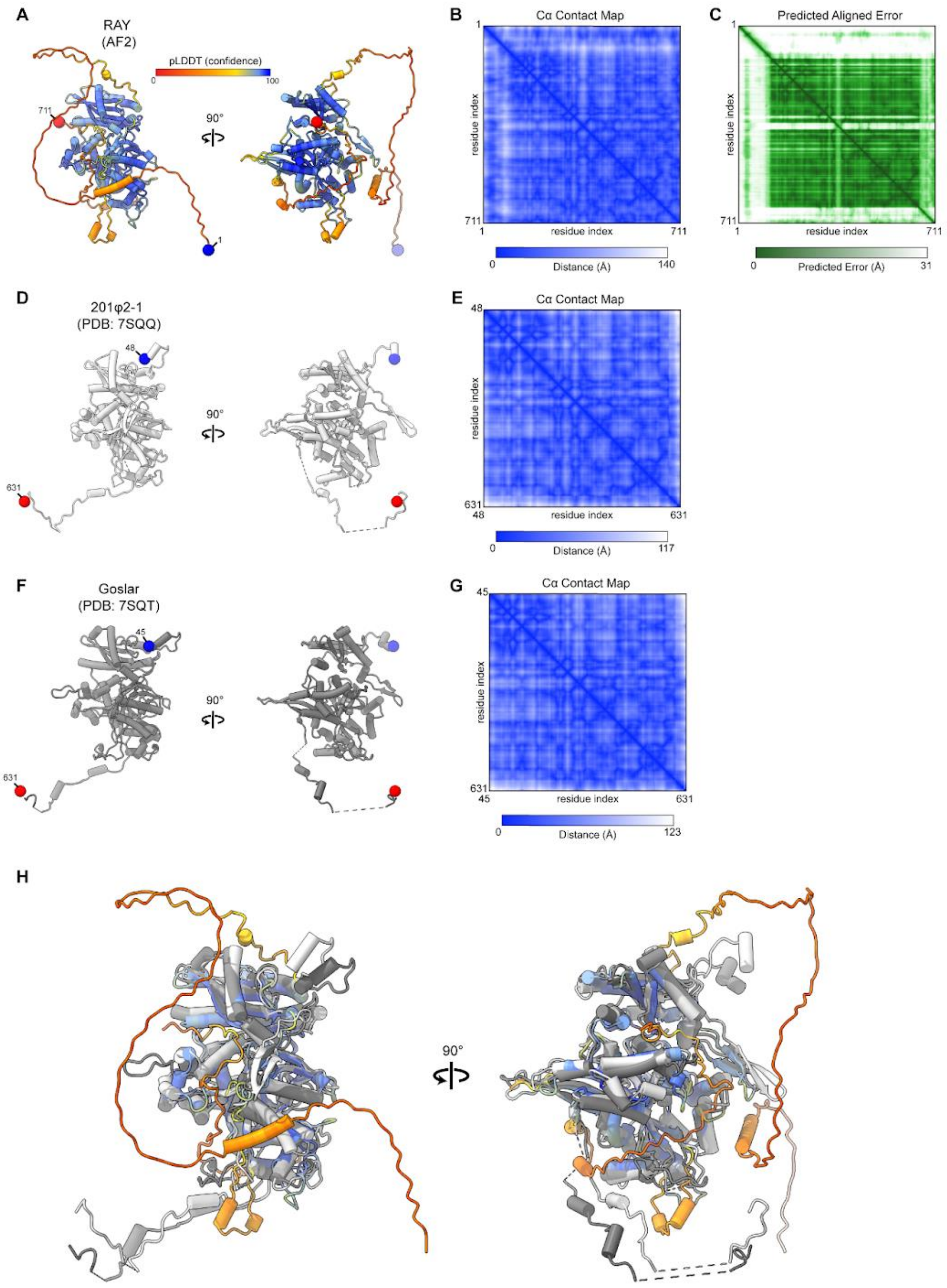


**Figure S22. Structural comparison of RAY, 201ϕ2-1, and Goslar chimallin protomers.** (A) Orthogonal views of the AF2 predicted RAY chimallin protomer model colored by pLDDT. N- and C-termini are labeled and shown as blue and red spheres, respectively. (B) Pairwise C-alpha distance map and (C) predicted aligned error plot for the RAY chimallin protomer model. (D) Orthogonal views of the 201ϕ2-1 chimallin protomer model (PDB ID: 7SQQ) colored white with termini indicated as in A and (E) corresponding pairwise C-alpha distance map. (F) Orthogonal views of the Goslar chimallin protomer model (PDB ID: 7SQT) colored gray with termini indicated as in A and (G) corresponding pairwise C-alpha distance map. (H) Orthogonal views of superimposed chimallin protomer models from A,D, and F.

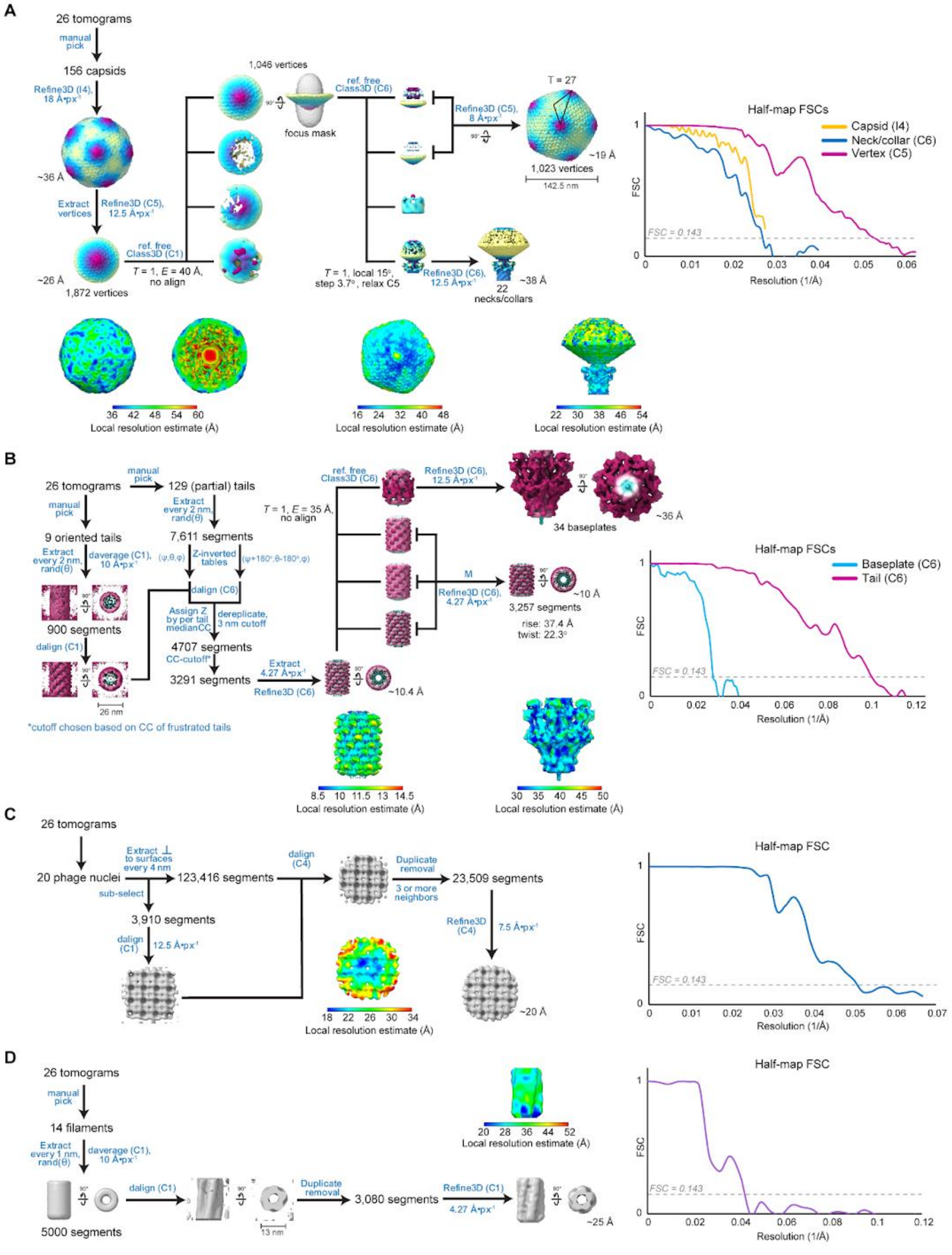

**Figure S23. Subtomogram analysis workflows of RAY components.** Workflow schematics, local resolution estimates, and half-map Fourier shell correlation curves for the RAY (A) capsid and collar, (B) tail sheath and baseplate, (C) chimallin, (D) putative PhuZ filament.

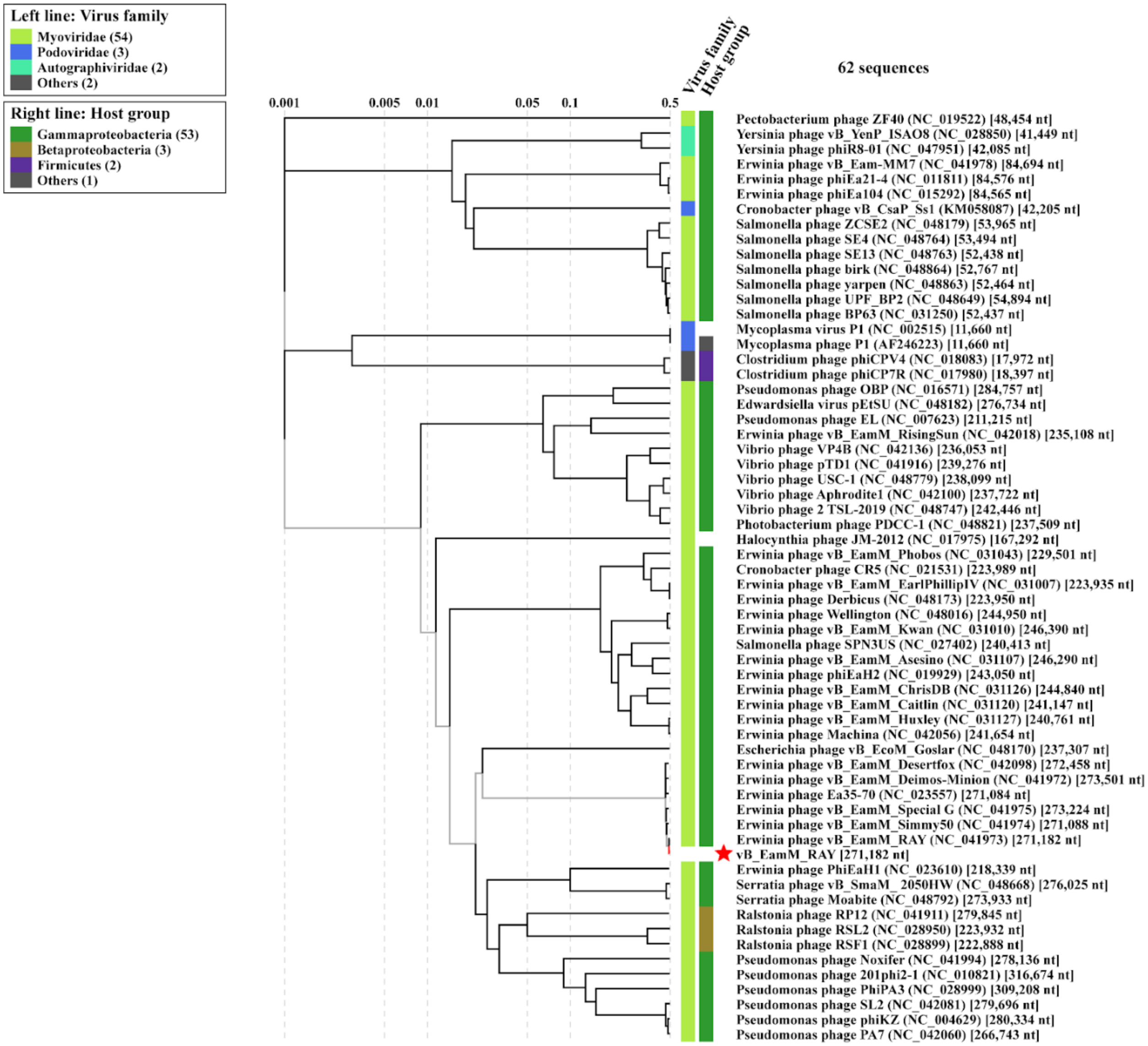


**Figure S24. ViPTree-predicted relatives of RAY.** Relatives of RAY were attempted to be found using ViPTree to identify similar genomes. This whole genome tree represents possible RAY relatives found by ViPTree (Nishimura et al., 2017).
